# A selectable system to evaluate synthetic gene optimization features

**DOI:** 10.1101/825976

**Authors:** Brevin A. Smider, Jacob Gil, Vaughn V. Smider

## Abstract

Heterologous gene expression – transferring genes from a natural cell or origin to another host cell, often across species – is a fundamental technique in biological research as well as biotechnological and pharmaceutical manufacturing. Optimization of gene sequences, therefore, is a critical factor when enhanced protein yield is needed. For any given protein, an enormous diversity of nucleotide sequences comprising the vast combinatorics of possible codon combinations could theoretically be used to encode the same amino acid sequence. The process of “codon optimization” typically replaces certain codons thought to be suboptimal with a more optimal codon encoding the same amino acid. However, such methods cannot address the enormity of possible nucleotide sequences possible to encode a given protein, or to decipher potential factors in addition to codon usage that may affect gene expression. Here we utilize the *Sh ble* gene encoding zeocin resistance in a system comprising bioinformatic synthetic gene production and an antibiotic selection platform. We find that supposedly codon optimized genes do not produce enhanced antibiotic resistance, suggesting that other factors are more important in synthetic gene optimization and heterologous gene expression.

## Introduction

Heterologous gene expression is a major experimental technique in nearly all areas of biological research, including development and commercialization of products in the biotechnology industry. In particular, optimized gene expression is important where milligrams to grams of recombinant product may be required, for example in structural biology studies, as well as for immunogen or recombinant drug production. Furthermore, optimal gene expression is especially critical in biopharmaceutical manufacturing, where recombinant proteins are produced for therapeutic use. Despite this importance, the mathematical magnitude of testing variants of gene expression constructs often precludes identification of the optimal gene sequence for heterologous expression. With recent computational biology advancements and cost reductions in parallel gene synthesis, the ability to test many gene variants simultaneously is now more feasible. The ultimate goal would be to computationally predict an optimal gene DNA sequence based on its protein amino acid sequence in the context of its expression host cell.

The challenge with designing heterologous gene constructs is that the degeneracy of the genetic code provides for an enormous number of possible nucleic acid sequences when a given protein sequence is reverse translated into DNA. The number of possible sequences is a function of both the length of the protein as well as the content of individual amino acids. In this regard, some amino acids like serine, leucine, and arginine are encoded by six different codons, whereas methionine and tryptophan are only encoded by one. As an example of the vast sequence space the genetic code provides, Table 1 illustrates the number of possible different nucleic acid sequences for several representative proteins. Of note, exenatide, which is an important peptide used clinically in diabetes (Byetta, Astra-Zeneca) is only 39 amino acids long, however the number of possible DNA sequences which could encode it is >10^19^. The largest protein known, titin, comprising over 26,000 amino acids, could theoretically be encoded by an astronomical 10^13,483^ different DNA sequences. Even for short proteins, the ability to produce and test the entire sequence landscape encoding them is out of reach with current expression techniques. Smaller libraries, however comprising thousands to over millions of sequences are constructible in a “one-pot” format. In this regard, a vast sequence space can be tested as long as an effective selection strategy is in place to identify “winner” sequences in the pool of variants. In this regard, phage display and other molecular evolution approaches have been utilized to engineer recombinant antibody fragments and enzymes for several years [REFS].

**Table 1.** Possible codon combinations for representative proteins

| Protein | Length (AA) | Number of Sequences |
| --- | --- | --- |
| Exenatide | 39 | $6.648 \times 10^{19}$ |
| Human insulin | 110 | $5.165 \times 10^{57}$ |
| Sh ble | 124 | $2.642 \times 10^{61}$ |
| erythropoietin | 166 | $1.134 \times 10^{89}$ |
| GCSF (neupogen/filgrastim) | 175 | $2.822 \times 10^{94}$ |
| Trastuzumab LC | 214 | $2.288 \times 10^{111}$ |
| Trastuzumab HC | 450 | $3.248 \times 10^{229}$ |
| Green Fluorescent Protein (GFP) | 238 | $2.645 \times 10^{109}$ |
| Titin | 26,926 | $2.618 \times 10^{13483}$ |

Historically, codon optimization was thought to be a critical component to gene expression optimization. Fundamentally, codon optimization presumes that the relative abundance of tRNAs encoding synonymous codons would impact translation rates, and thus protein expression. Indeed, several studies correlated highly expressed genes with the codon adaptation index (CAI) [REFS]. The CAI is a measure of codon usage bias defined by assessing the relative merits of each synonymous codon in a gene with a reference set of highly expressed genes from that species[1]. The CAI has been utilized as a quantitative measure to predict gene expression, and several companies provide services for ‘codon optimization’ which utilize the CAI to enhance gene expression [REFS]. Many codon optimization algorithms have been examined, with mixed results in terms of predicting gene expression [REFS]. Some studies, however, show that codon optimization is not necessarily a major factor associated with gene expression. In particular, a study by Kudla, et.al. examined several sequence variants of GFP and found that protein expression was not correlated with CAI, but was more correlated with mRNA stability at the 5’ end of the gene[2]. Other factors like mRNA turnover, the role of translation speed with protein folding and stability, also likely impact protein expression [REFS]. Thus, it is unclear what the relative importance of these factors are to heterologous gene optimization, or whether other unknown sequence intrinsic mechanisms may additionally impact gene expression.

In order to assess the sequence space associated with optimal gene expression, a system to quantitatively select libraries of gene sequences encoding the identical protein would be a useful advance, in that a synthetic gene library encoding a single protein could be used under selective pressure to identify sequences that outperform the remaining library pool. In this regard, winning sequences could be examined for features associated with their optimal expression. In initial studies to develop such a platform, here we utilize the *Streptoalloteichus hindustanus ble* gene (*sh ble*) encoding bleomycin and zeocin resistance in a selectable system to evaluate optimal gene features.

## Results

To establish a system to evaluate multiple different DNA sequences encoding an identical protein for optimal expression properties, we designed software capable of reverse translation of protein sequences, with the input of various filters for sequence features (Figure 1). At a high level, a user input for the number of sequences to be output, as well as threshold values for other features such as CAI and %GC in desired output sequences were included. We chose the *sh ble* gene as proof of concept for two significant reasons: (i) it is a relatively small gene, encoding an amino acid sequence of only 124 residues, therefore the sequence space to be explored is relatively smaller than for larger genes, and (ii) it provides a readily selectable function of antibiotic resistance. Importantly, *sh ble* encodes a protein which directly binds and neutralizes bleomycin or zeocin at a 1:1 ratio. Therefore, unlike catalytic antibiotic resistance proteins like β-lactamase, the amount of antibiotic resistance for *sh ble* should be directly proportional to the amount of gene expression.

**Figure 1.**
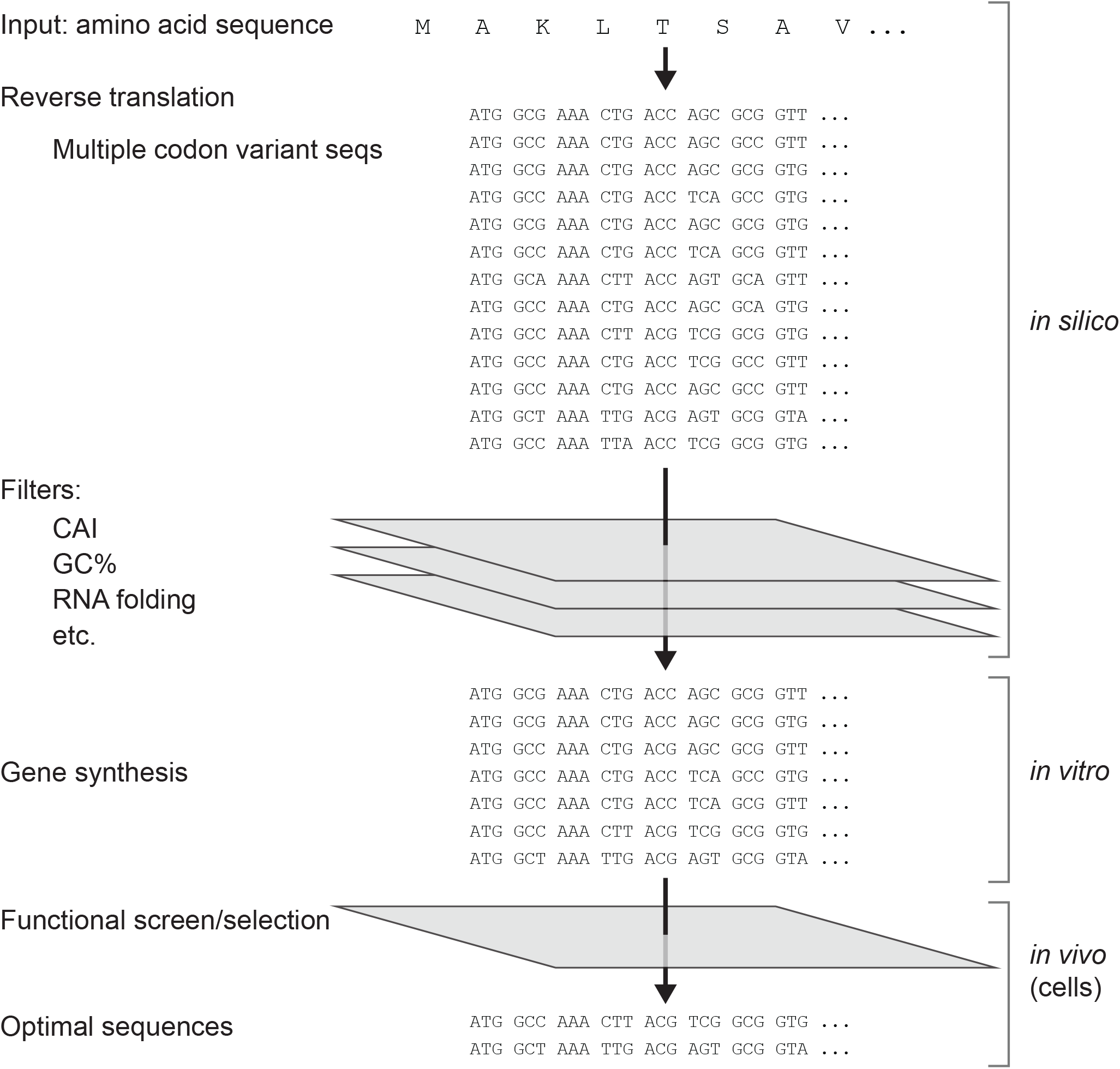
Gene optimization schematic. A gene optimization system comprising *in silico* reverse translation into a group of DNA sequences encoding an identical protein (top), where an amino acid sequence is used as input. Several filters could be employed to select sequences with different properties such as codon adaptation index (CAI), guanosine plus cytosine content (GC%) or RNA folding properties. Sequences fulfilling the filtered criteria are synthesized in an expression vector (middle), then selected for expression or function in cells (bottom), and identification of the optimal DNA sequences.

Using a random reverse translation algorithm to bias codons towards higher or lower CAI for a given organism (Supplemental Methods) we randomly produced 100 sequences with relatively higher CAI *in silico* and determined that they were unrelated to one another, with the percent identity at the nucleotide level ranging from 73%-90% (Supplemental Figures 1 and 2). We also produced ten sequences with low CAI (Seq101-Seq110), as well as a sequence with the optimal codon for *E.coli* at each position (Seq111). We additionally converted any poor codons in the wild-type sequence to the optimal codon, as a simple mechanism for codon optimization (Seq112). In order to establish the utility of the system at a small scale, nine total genes were chosen based on a range of homology with one another (ranging from 74% to 94%, Supplemental Figure 3). Seq111 and 112 were 94% identical as both of these sequences had a significant fraction of optimal codon usage. Seven genes had high CAI values, two had low values (Seqs 109 and 103), one was the best codon sequence with a CAI of 1.0 (Seq111), and one was the wild-type sequence from *Streptoalloteichus hindustanus*. The wild-type sequence notably has a much higher GC content than the other sequences, at nearly 70% compared to a range of 52-58% in the remaining eight sequences. The nine genes were synthesized and inserted into the pTWIST-CMV vector but under the control of the lac promoter for *E.coli* expression (Supplemental Figure 4). The only differences between the genes are the coding sequences; no differences in 5’ or 3’ non-coding regions were present, allowing for selection solely based on gene sequence related to expression.

**Figure 2.**
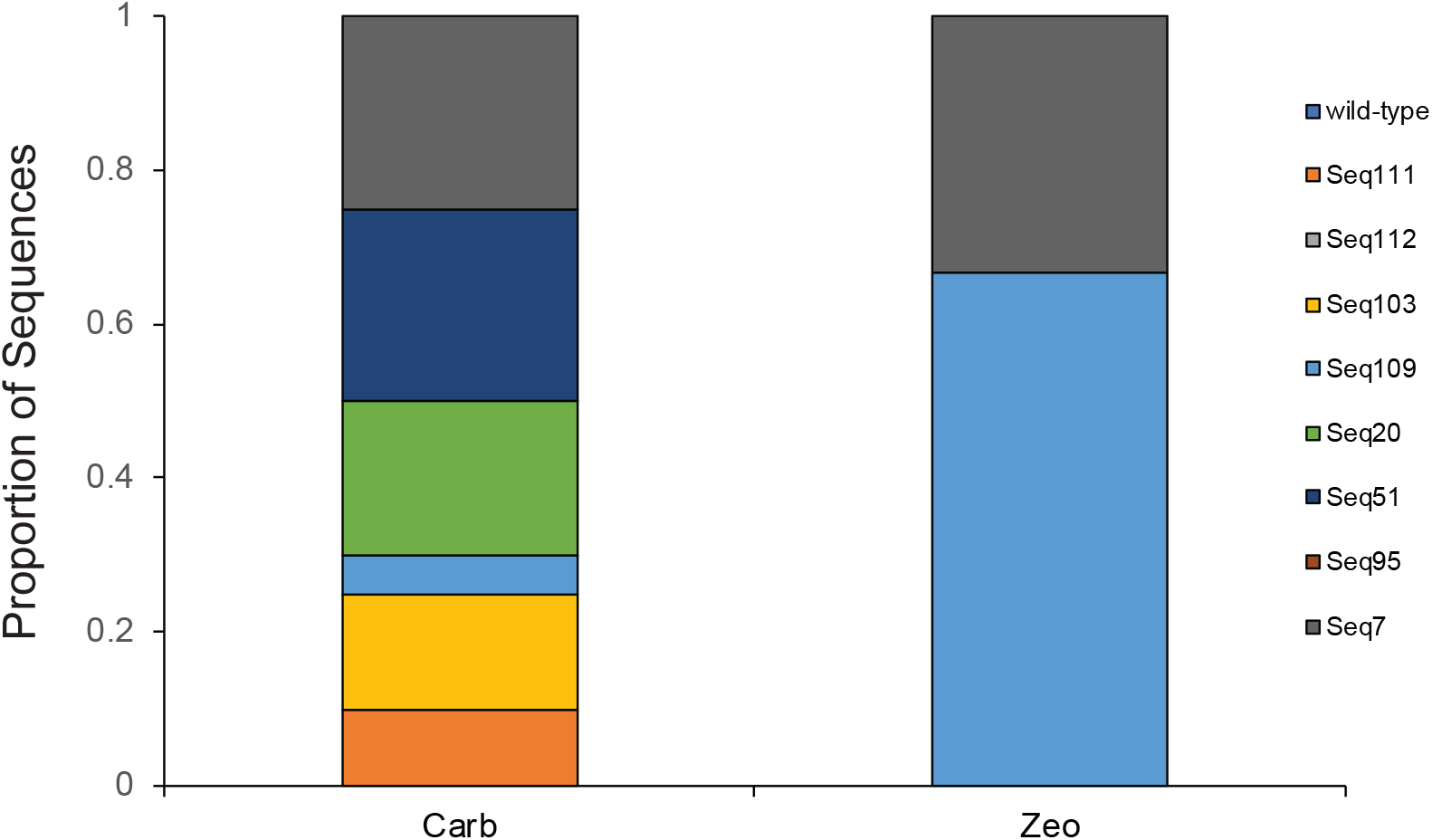
Selection of optimal *sh ble* genes in *E.coli*. Nine sequences all encoding *sh ble* were selected for growth overnight in either carbenicillin (Carb) or zeocin (Zeo) in liquid culture then plated on carbenicillin containing agar plates and colonies sequenced. The proportion of each sequence in each population is represented.

In order to identify expression differences, we plated the gene variants on different concentrations of zeocin, ranging from 5 to 50 μg/ml. Remarkably, the CAI had no predictable impact on zeocin resistance (Table 2, Supplemental Figure 5). The wild-type protein could form colonies at 10 μg/ml, but the sequence (111) with a CAI of 1.0 could also only form colonies at 10 μg/ml. Interestingly, Seq7 and Seq109 could both form colonies at the highest concentration, 50 μg/ml (Supplemental Figure 5). The resistance to zeocin was also reflected in colony size, as Seq7 and Seq109 formed larger colonies at lower zeocin concentrations than the wild-type or other sequences. Some sequences including Seq20, Seq51, and Seq103 could form small colonies at 25 μg/ml, and larger colonies than wild-type at 10 μg/ml zeocin.

**Table 2.** Features of *sh ble* codon variants

| Sequence | CAI | GC% | MFold $\Delta G$<br>Minimum<br>(kcal/mol) | MFold $\Delta G$<br>Maximum<br>(kcal/mol) | MFOLD $\Delta G$<br>-4 to 37<br>(kcal/mol) | Miniprep<br>Yield (ng/μl) |
| --- | --- | --- | --- | --- | --- | --- |
| Wild-type | 0.71 | 69.87% | -82.72 | -79.77 | -8.60 | 47.2 |
| Seq7 | 0.88 | 54.66% | -71.74 | -68.18 | -9.20 | 52.1 |
| Seq20 | 0.87 | 56.18% | -77.65 | -74.11 | -10.00 | 111.8 |
| Seq51 | 0.86 | 55.75% | -67.34 | -64.09 | -8.50 | 57.6 |
| Seq95 | 0.84 | 56.18% | -67.87 | -64.58 | -8.10 | 108.9 |
| Seq103 | 0.70 | 54.66% | -59.31 | -56.68 | -6.20 | 42.7 |
| Seq109 | 0.68 | 52.06% | -57.68 | -54.80 | -3.10 | 41.5 |
| Seq111 | 1.0 | 58.79% | -80.52 | -77.67 | -11.40 | 89.2 |
| Seq112 | 0.97 | 58.13% | -76.82 | -73.80 | -14.60 | 97.5 |

To verify these differences in zeocin resistance in the context of a competitive pool of sequences, we mixed *E.coli* harboring the nine sequences and evaluated which sequences would be selected in zeocin-containing cultures. Indeed, sequences 7 and 109 were clearly selected over all of the other sequences (Figure 2), with Seq7 representing 1/3 of the sequences and Seq109 representing 2/3 of the colonies grown after an overnight culture. Seq7 and Seq109 do not share substantial homology compared with the remaining seven sequences. Seq7 and Seq109 are 77.87% identical whereas Seq7 shares greater homology with six of the remaining seven sequences. Seq7 has greater homology with Seq103 (78.13%), Seq95 (84%), Seq51 (82.93%), Seq20 (82.70%), Seq111 (88.53%), and Seq112(88.27%). The only sequence with lower homology is the wild-type *sh ble* sequence. Seq109, on the other hand, has higher homology with the wild-type *sh ble* sequence (78.13%) and slightly lower homology with the remaining sequences other than Seq7 (between 74.13% and 77.33%). A pairwise alignment between Seq109 and Seq7 did not reveal any obvious homology that may mediate the expression effect. The polymorphic nucleotides are spread out over the length of the genes with the longest stretch of complete homology being 14 residues from position 103-116. We also utilized the Mfold algorithm to predict RNA folding. No obvious correlation was found between Mfold ΔG and Seq7 and Seq109, either over the entire gene or in the 5’ region[2]. While we cannot firmly identify features mediating high expression in this simple system, the *sh ble* gene can be used in a system used to select enhanced gene variants. This simple proof of concept suggests larger libraries may be used to identify many enhanced variants, and analyses of these ‘winner’ sequences may identify features more clearly associated with enhanced expression.

## Conclusions and Discussion

The ability to optimize genes for expression properties is important in many areas of biological research. While incorporation of cell-type specific promotors, enhancers, or other genetic elements are important in optimizing expression cassettes, the optimization of the target gene’s sequence itself is also important to establish high level expression. Previous studies have implicated CAI and GC content as important parameters in optimizing gene expression, but recent analysis of several GFP sequences show a low correlation of expression with CAI or GC content, but greater correlation with the RNA folding properties at the 5’ region of the sequence. We did not find an obvious difference n Mfold DG at the 5’ end of Seq7 and Seq109, however this experiment used a very small sample size with only two ‘winning’ sequences. Notably Seq109 had the highest ΔG at the 5’ end (Table 2) but this relevance is unclear given no particular difference was seen in the 5’ ΔG for Seq7 compared to the other sequences. Importantly, however, our study supports the notion that CAI is not critically important for gene expression, and that other factors must be involved in enhanced gene expression.

There are many programs that offer sequence-based algorithms to optimize gene expression [3]. We established a simple system based on zeocin resistance which can evaluate gene expression in parallel, or in a competitive culture. Here, libraries of genes with different codon content but encoding identical amino acid sequences can be evaluated to identify sequences that confer the highest resistance to zeocin, based on gene expression. Despite the importance of gene expression in many areas of biological research and biopharmaceutical manufacturing, relatively little is understood about the key parameters needed to optimize a gene. It is likely that many factors will impact expression, and many of these could be species or even cell-type specific. For example, RNA stability, transport, impact on cell growth, interaction with other RNAs through hairpin structures, etc., all could impact gene expression. In this regard, the ability to make libraries of genes and evaluate them in different cells could enable identification of unique cis-acting factors of gene sequences that positively or negatively impact their expression. Of note, the *sh ble* gene can mediate zeocin resistance in bacteria as well as eukaryotic cells, and in multiple cell types of different species, and therefore could be a useful model system to evaluate such sequence-specific expression properties.

## Methods

Bioinformatic analysis. The reverse translation and sequence filtering program were written in JAVA as detailed in Supplemental Methods. RNA folding analysis was done by Mfold[4, 5].

Zeocin resistance. Wild-type and codon variants of *sh ble* were synthesized by Twist Biosciences (South San Francisco, CA) in the TWIST-CMV vector (Supplemental Figure 4). These plasmids produce the β-lactamase protein conferring carbenicillin resistance in addition to the *sh ble* gene conferring zeocin resistance. Plasmids were transformed into DH5α *E.coli* (Thermo-Fisher, Carlsbad, CA) and selected on carbenicillin (50 μg/ml) LB-agar plates. For evaluation of resistance to zeocin on plates, LB-agar containing 5, 10, 25, and 50 μg/ml zeocin were prepared and codon variant plasmid-containing *E.coli* were streaked on each plate using a wire loop. Plates were incubated overnight at 37°C then visualized by imaging with a Chemi-Doc camera and QuantityOne imaging software (Bio-Rad, Hercules, CA). *E.coli* containing pUC19, that did not contain *sh ble* gene, were analyzed for zeocin resistance and were confirmed to not grow on LB-agar plates containing 5 μg/ml zeocin. For competition experiments, single colonies of *E.coli* containing plasmids encoding each codon variant were picked and grown overnight in terrific broth containing carbenicillin at 50 μg/ml. On the next day 10^6^ cells from each codon variant were pooled and grown overnight in either 50 μg/ml carbenicillin or zeocin in TB at 37°C. The mixed cultures were plated on 50 mg/ml carbenicillin LB-agar plates on the next day at approximately 100 cfu per plate, then plasmids from individual colonies analyzed by Sanger sequencing (Genewiz, San Diego, CA). Sequences were identified by homology analysis using the Bioconductor program in R Studio[6].

---

**Supplementary Figure 1.** Sequences of 110 randomly generated DNA sequences encoding *sh ble*.

**Supplementary Figure 2.** Percent identity matrix of 110 *sh ble* encoding sequences. The percent identity ranges between 73-90%.

**Supplemental Figure 3.** Nine sequence variants of *sh ble* evaluated in *E.coli*.

**Supplemental Figure 4.** Map of *sh ble* expression vector.

**Supplemental Figure 5. Zeocin resistance of *sh ble* codon variant genes.** *E.coli* containing the nine variant genes were streaked on zeocin containing agar plates with concentrations of 5, 10, 25, and 50 μg/ml, grown overnight and visualized.

## Acknowledgements

This work was funded by NIH grants R01 GM105826 and R01 HD088400 to V.V.S.

## Author Contributions

B.A.S. wrote the bioinformatic program, generated sequences, performed zeocin resistance experiments, and analyzed data. J.G. performed experiments, analyzed data, and assisted in algorithm design. V.V.S. supervised the project and wrote the manuscript. All authors read and approved the manuscript.

