## Supplemental Figure 1 for "A selectable system to evaluate synthetic gene optimization features"

>Sequence1

ATGGCGAAACTGACCAGCGCGGTTCCGGTGCTTACCGCACGCGATGTGGCGGGTGCCGTGGAGTTTTGGACCGATCGCCTGGGCTTTTCACGTGATTTTGTTGAAGATGACTTCGCGGGTGTAGTGCGCGATGATGTGACCCTGTTCATTAGCGCCGTTCAGGATCAGGTGGTCCCGGATAACACCCTGGCCTGGGTTTGGGTGCGCGGCTTAGATGAGCTGTATGCCGAATGGTCGGAAGTTGTGAGCACTAACTTTCGTGATGCGAGCGGTCCGGCGATGACCGAAATTGGTGAACAACCGTGGGGCCGTGAATTCGCGTTGCGTGATCCGGCGGGCAACTGCGTGCATTTTGTTGCCGAAGAACAGGATTAA

>Sequence2

ATGGCGAAACTGACCAGCGCGGTGCCGGTCCTGACCGCCCGCGATGTCGCGGGTGCCGTTGAATTCTGGACCGATCGCCTGGGCTTCAGCCGTGATTTTGTTGAAGATGATTTTGCCGGCGTAGTTCGTGATGACGTGACCCTGTTCATCAGCGCAGTGCAGGATCAGGTTGTACCGGATAACACCCTGGCGTGGGTGTGGGTGCGCGGCTTGGATGAGCTGTATGCGGAGTGGTCAGAAGTGGTGAGCACGAACTTTCGTGATGCATCGGGTCCGGCGATGACTGAAATTGGTGAACAGCCGTGGGGCCGCGAATTTGCGTTACGTGATCCGGCGGGCAACTGCGTGCATTTTGTAGCCGAAGAACAGGACTAA

>Sequence3

ATGGCCAAACTGACTAGCGCGGTACCGGTGCTGACGGCCCGCGACGTTGCGGGTGCCGTCGAGTTTTGGACAGATCGTCTGGGCTTTAGCCGCGACTTTGTGGAAGATGATTTCGCGGGCGTAGTCCGCGATGATGTTACCCTGTTTATTTCGGCCGTGCAGGATCAAGTAGTTCCGGATAACACGCTGGCCTGGGTGTGGGTGCGTGGACTGGATGAACTGTATGCGGAGTGGAGCGAAGTTGTGAGCACCAACTTCCGTGATGCCAGCGGCCCGGCCATGACCGAAATTGGTGAACAGCCGTGGGGCCGCGAATTTGCGCTGCGTGATCCGGCGGGCAACTGCGTGCATTTTGTTGCCGAAGAACAGGATTAA

>Sequence4

ATGGCGAAACTGACGTCGGCTGTGCCGGTTCTGACGGCACGCGATGTGGCGGGCGCGGTGGAATTTTGGACCGACCGCCTGGGTTTTAGCCGTGATTTCGTTGAAGATGATTTTGCCGGAGTGGTGCGCGATGACGTGACACTGTTTATTAGCGCAGTGCAGGATCAGGTTGTTCCGGATAACACCCTGGCGTGGGTGTGGGTCCGTGGTCTGGATGAACTGTATGCCGAATGGAGCGAGGTGGTGTCAACCAATTTCCGTGATGCCAGCGGCCCGGCCATGACCGAAATCGGCGAGCAGCCGTGGGGTCGCGAGTTTGCGCTGCGCGACCCGGCAGGCAACTGCGTGCATTTTGTGGCCGAAGAACAGGATTAA

>Sequence5

ATGGCCAAACTGACTAGCGCGGTGCCGGTGCTTACCGCCCGCGATGTGGCGGGCGCCGTGGAGTTTTGGACAGATCGTCTGGGCTTCAGCCGTGATTTTGTTGAAGATGATTTTGCGGGCGTCGTTCGTGATGACGTGACCCTGTTCATTTCGGCGGTCCAGGACCAGGTCGTGCCGGATAACACCTTGGCGTGGGTGTGGGTGCGCGGCCTGGATGAACTGTATGCCGAGTGGAGCGAAGTTGTTAGCACCAACTTTCGTGATGCCAGCGGTCCGGCGATGACCGAAATTGGTGAGCAGCCGTGGGGCCGCGAATTCGCGCTGCGCGACCCGGCTGGCAACTGCGTTCATTTTGTAGCGGAAGAACAGGATTAA

>Sequence6

ATGGCCAAACTGACTAGCGCGGTGCCTGTGCTGACGGCGCGTGATGTGGCCGGGGCGGTGGAATTTTGGACCGATCGCCTCGGCTTCAGTCGTGACTTTGTTGAAGATGATTTTGCGGGAGTGGTGCGCGACGATGTCACCCTGTTTATTAGCGCCGTGCAGGATCAGGTGGTTCCGGACAACACCCTGGCGTGGGTGTGGGTGCGCGGTCTGGATGAACTGTATGCGGAATGGTCAGAAGTTGTGAGCACCAACTTCCGTGACGCGAGCGGTCCGGCCATGACCGAAATTGGCGAACAGCCGTGGGGCCGCGAATTCGCGTTACGCGATCCGGCGGGCAACTGCGTTCATTTTGTGGCTGAGGAGCAGGATTAA

>Sequence7

ATGGCCAAACTGACGAGCGCGGTTCCGGTCCTGACCGCGCGTGATGTCGCAGGCGCGGTGGAATTTTGGACCGATCGCCTGGGCTTTAGTCGCGACTTTGTTGAGGACGATTTTGCCGGCGTGGTGCGCGATGATGTAACCTTATTTATTTCAGCGGTGCAGGATCAGGTGGTTCCGGATAACACCCTGGCCTGGGTTTGGGTTCGCGGCCTCGATGAACTGTATGCGGAATGGAGCGAAGTGGTATCGACCAATTTTCGTGATGCTAGCGGTCCGGCGATGACGGAAATCGGTGAACAGCCGTGGGGGCGCGAATTTGCCCTGCGTGATCCGGCAGGTAACTGCGTTCATTTTGTGGCCGAAGAACAAGATTAA

>Sequence8

ATGGCGAAACTGACCAGCGCAGTGCCGGTTCTCACGGCCCGTGATGTGGCTGGTGCGGTGGAATTTTGGACCGACCGTCTGGGCTTTAGCCGCGATTTTGTGGAAGATGATTTCGCCGGCGTGGTGCGTGATGATGTGACCCTGTTTATTAGCGCCGTTCAAGACCAGGTGGTTCCGGATAACACCCTGGCCTGGGTGTGGGTGCGCGGCCTGGACGAACTGTATGCGGAGTGGTCGGAAGTGGTCTCAACCAACTTTCGCGACGCCAGCGGCCCGGCGATGACTGAGATCGGGGAACAGCCGTGGGGTCGCGAATTTGCCCTGCGTGATCCGGCAGGCAACTGCGTTCATTTCGTGGCGGAAGAACAGGACTAA

>Sequence9

ATGGCCAAACTGACCAGCGCCGTTCCGGTTTTAACCGCCCGCGATGTGGCGGGTGCGGTAGAATTTTGGACCGATCGCCTGGGCTTTAGCCGCGATTTCGTTGAAGATGACTTTGCGGGTGTTGTGCGTGATGATGTTACCCTTTTTATTAGCGCGGTGCAGGACCAGGTTGTGCCGGATAACACGCTCGCATGGGTGTGGGTACGTGGCCTGGATGAGCTGTATGCCGAATGGAGCGAAGTTGTAAGTACCAACTTTCGCGATGCATCGGGCCCGGCAATGACTGAAATTGGCGAACAGCCGTGGGGCCGTGAATTCGCCCTGCGCGATCCGGCCGGTAATTGCGTGCATTTCGTGGCGGAAGAGCAGGATTGA

>Sequence10

ATGGCGAAACTGACCTCAGCGGTGCCGGTTCTTACCGCGCGTGACGTGGCCGGCGCCGTAGAATTTTGGACGGATCGCCTGGGCTTTAGCCGCGATTTTGTGGAAGATGATTTTGCTGGTGTGGTCCGTGATGATGTAACCCTGTTTATTAGCGCGGTGCAGGATCAGGTAGTTCCGGACAATACCCTGGCGTGGGTCTGGGTTCGCGGCCTGGATGAACTCTATGCAGAATGGAGCGAAGTTGTTAGCACGAACTTTCGCGATGCCTCGGGCCCGGCAATGACCGAAATTGGCGAACAACCGTGGGGCCGCGAGTTTGCCTTACGTGACCCGGCGGGTAACTGCGTGCATTTCGTTGCGGAAGAACAGGACTAA

>Sequence11

ATGGCCAAACTCACCAGCGCCGTGCCGGTGCTGACCGCCCGTGATGTCGCGGGCGCCGTTGAATTCTGGACCGATCGCTTGGGCTTTAGCCGCGATTTTGTGGAAGATGATTTTGCGGGCGTGGTTCGCGATGATGTGACCCTGTTTATCAGCGCCGTCCAAGATCAGGTTGTACCGGATAACACGCTGGCGTGGGTTTGGGTGCGTGGGTTAGACGAACTGTATGCTGAATGGAGCGAGGTTGTTAGCACCAACTTTCGCGATGCCAGTGGTCCGGCAATGACGGAAATTGGAGAGCAGCCGTGGGGCCGCGAATTTGCCCTGCGTGATCCGGCGGGTAACTGCGTGCATTTTGTCGCGGAAGAACAGGATTAA

>Sequence12

ATGGCGAAACTGACCAGCGCCGTGCCGGTGCTGACCGCGCGCGATGTCGCGGGTGCCGTGGAATTTTGGACCGATCGTCTCGGCTTTAGCCGCGATTTTGTTGAGGATGATTTTGCCGGCGTTGTTCGTGATGATGTTACCCTGTTCATTAGTGCTGTGCAGGACCAAGTGGTGCCGGATAATACCCTGGCGTGGGTTTGGGTGCGTGGACTGGACGAACTGTATGCGGAATGGAGCGAAGTGGTATCGACTAACTTCCGTGATGCAAGCGGCCCGGCTATGACGGAAATTGGCGAGCAGCCGTGGGGTCGCGAATTTGCGCTGCGCGATCCGGCGGGTAACTGCGTCCATTTCGTTGCCGAAGAACAGGATTGA

>Sequence13

ATGGCTAAACTGACCAGCGCAGTTCCGGTGTTAACCGCGCGTGATGTGGCGGGCGCGGTGGAATTCTGGACGGATCGCCTTGGGTTTAGCCGCGATTTTGTTGAAGATGACTTCGCCGGTGTGGTTCGCGACGACGTGACCCTGTTTATTAGCGCTGTTCAGGATCAGGTTGTGCCGGATAACACCCTGGCCTGGGTATGGGTGCGCGGTCTGGATGAACTGTATGCAGAGTGGAGCGAAGTTGTGAGCACGAACTTTCGTGATGCGAGCGGCCCTGCCATGACAGAAATTGGTGAACAGCCGTGGGGCCGCGAATTTGCCCTGCGTGATCCGGCCGGCAACTGCGTGCATTTTGTGGCGGAGGAACAGGATTGA

>Sequence14

ATGGCCAAACTGACCAGCGCGGTGCCGGTCCTGACGGCGCGCGATGTGGCAGGCGCCGTTGAATTCTGGACCGATCGCCTGGGCTTTAGCCGCGATTTCGTTGAGGATGATTTTGCTGGCGTAGTGCGCGACGACGTGACTCTGTTTATCAGCGCCGTGCAGGATCAGGTAGTTCCGGACAACACCTTAGCGTGGGTTTGGGTGCGTGGCCTGGATGAGCTGTATGCCGAATGGAGCGAAGTAGTTTCAACGAATTTTCGTGATGCGAGCGGCCCGGCGATGACCGAAATCGGAGAACAGCCGTGGGGTCGTGAATTTGCACTGCGTGATCCGGCGGGTAACTGCGTGCATTTTGTGGCCGAAGAACAGGACTAA

>Sequence15

ATGGCGAAACTGACTAGCGCCGTTCCGGTGCTGACCGCGCGCGACGTCGCGGGCGCGGTAGAGTTCTGGACCGATCGCCTGGGTTTTTCACGTGATTTTGTGGAAGACGACTTCGCCGGCGTGGTGCGCGATGATGTGACCCTGTTTATCAGCGCGGTCCAGGATCAAGTCGTGCCGGATAACACCCTGGCGTGGGTTTGGGTGCGTGGCCTCGATGAACTGTATGCAGAATGGAGCGAAGTGGTTAGCACGAATTTTCGTGATGCATCGGGCCCGGCGATGACCGAAATTGGTGAACAGCCGTGGGGCCGCGAATTCGCCCTGCGCGACCCGGCGGGGAACTGCGTTCATTTTGTGGCCGAAGAGCAGGACTAA

>Sequence16

ATGGCAAAATTAACCAGCGCGGTTCCGGTCCTGACGGCGCGCGATGTAGCAGGTGCGGTCGAATTTTGGACCGACCGTCTGGGCTTCAGCCGCGATTTTGTGGAAGATGATTTTGCGGGAGTGGTACGCGACGATGTTACCCTGTTCATTAGCGCCGTGCAGGATCAGGTCGTTCCGGATAACACGCTGGCCTGGGTGTGGGTGCGTGGCTTGGATGAACTGTATGCGGAATGGAGCGAAGTGGTGTCGACCAACTTCCGTGACGCGTCAGGCCCGGCCATGACCGAAATTGGGGAACAGCCTTGGGGTCGTGAATTTGCCCTGCGCGACCCGGCCGGCAACTGCGTTCATTTTGTTGCCGAAGAACAGGATTGA

>Sequence17

ATGGCCAAATTAACCTCGGCGGTGCCTGTTCTGACGGCGCGCGATGTGGCCGGCGCCGTGGAATTCTGGACCGATCGTCTGGGCTTCTCACGTGACTTTGTGGAAGATGATTTTGCCGGCGTGGTGCGCGATGACGTAACACTGTTCATTAGCGCGGTTCAGGATCAAGTGGTGCCGGATAACACCCTTGCGTGGGTATGGGTACGTGGCCTGGATGAATTGTATGCGGAATGGAGCGAAGTTGTTAGCACCAACTTTCGTGATGCCAGCGGTCCGGCTATGACTGAAATTGGTGAACAGCCGTGGGGCCGCGAATTTGCTCTGCGCGATCCGGCCGGAAACTGCGTCCATTTTGTGGCCGAAGAACAGGATTGA

>Sequence18

ATGGCCAAACTGACGAGCGCCGTGCCGGTTCTCACAGCCCGCGACGTGGCGGGTGCAGTAGAATTCTGGACCGATCGCCTGGGCTTTTCGCGCGATTTTGTTGAGGATGACTTCGCGGGCGTGGTGCGTGATGATGTAACCCTGTTTATTAGCGCCGTGCAGGACCAGGTGGTCCCGGATAACACCCTGGCTTGGGTTTGGGTGCGCGGTCTGGATGAGCTGTATGCAGAATGGAGCGAAGTTGTGAGCACTAACTTTCGCGACGCATCAGGCCCGGCGATGACCGAAATTGGTGAGCAGCCGTGGGGTCGTGAATTTGCGCTGCGTGACCCGGCGGGCAACTGCGTGCATTTTGTGGCGGAAGAACAGGATTAA

>Sequence19

ATGGCCAAACTTACGAGCGCGGTGCCGGTCCTGACCGCGCGTGATGTGGCTGGTGCCGTTGAATTTTGGACCGATCGTCTGGGTTTTAGCCGCGACTTTGTTGAGGATGATTTTGCTGGTGTGGTTCGCGATGACGTGACACTGTTTATTAGCGCGGTGCAGGATCAAGTTGTGCCGGACAACACCCTGGCCTGGGTGTGGGTGCGCGGCCTGGACGAATTGTATGCAGAATGGAGTGAAGTAGTGAGCACCAATTTCCGTGATGCATCGGGCCCGGCAATGACCGAAATTGGCGAACAGCCTTGGGGCCGTGAATTTGCCCTGCGCGATCCGGCGGGCAACTGCGTACATTTTGTAGCGGAAGAACAGGATTAA

>Sequence20

ATGGCCAAACTGACCTCAGCCGTGCCGGTGCTCACCGCCCGTGATGTGGCGGGCGCGGTTGAGTTTTGGACTGACCGTCTGGGCTTTAGCCGCGACTTTGTAGAAGACGATTTCGCGGGTGTTGTTCGTGATGATGTGACATTATTTATCAGCGCGGTCCAGGACCAGGTGGTGCCGGATAACACCCTGGCGTGGGTGTGGGTGCGCGGCCTGGATGAACTGTATGCGGAATGGAGCGAAGTTGTGAGCACCAACTTCCGTGATGCGAGCGGCCCGGCGATGACCGAAATTGGTGAGCAACCGTGGGGTCGCGAATTCGCCCTTCGCGATCCGGCAGGGAACTGCGTTCATTTTGTGGCAGAAGAACAGGATTAA

>Sequence21

ATGGCCAAACTCACCTCAGCGGTTCCGGTCCTGACCGCGCGTGATGTGGCCGGTGCAGTGGAATTTTGGACGGATCGCTTAGGCTTTAGCCGTGACTTTGTTGAAGATGATTTTGCCGGCGTGGTGCGCGACGACGTGACCCTGTTTATTTCGGCGGTGCAGGATCAGGTAGTGCCGGATAACACTCTGGCTTGGGTGTGGGTTCGCGGTCTGGACGAACTGTATGCCGAATGGAGCGAAGTAGTGAGCACCAACTTTCGTGATGCGAGCGGTCCGGCAATGACCGAGATTGGCGAACAGCCGTGGGGACGTGAATTCGCGCTGCGCGACCCGGCCGGCAACTGCGTGCATTTCGTTGCCGAAGAACAAGATTGA

>Sequence22

ATGGCGAAACTGACCAGCGCGGTGCCGGTTCTGACCGCACGTGATGTGGCCGGCGCAGTGGAATTTTGGACCGATCGTTTAGGTTTTAGCCGCGACTTTGTCGAAGATGATTTTGCCGGCGTAGTTCGTGACGATGTTACGCTGTTTATTAGCGCAGTCCAGGATCAGGTGGTACCGGACAACACTCTGGCCTGGGTCTGGGTTCGCGGGCTGGATGAACTGTATGCTGAATGGTCAGAAGTGGTTAGCACCAATTTTCGCGATGCCTCGGGTCCTGCCATGACCGAAATTGGCGAACAGCCGTGGGGCCGCGAATTCGCGCTGCGTGACCCGGCGGGTAACTGCGTACATTTCGTGGCGGAGGAACAGGATTAA

>Sequence23

ATGGCCAAACTCACCAGCGCAGTTCCGGTTCTGACCGCTCGCGATGTGGCGGGCGCCGTGGAATTTTGGACCGATCGTCTGGGCTTCAGCCGCGATTTCGTGGAAGATGACTTTGCGGGCGTGGTTCGTGATGATGTTACCTTATTTATTAGCGCGGTGCAGGACCAGGTCGTGCCGGACAACACGCTGGCCTGGGTGTGGGTTCGTGGCCTGGATGAACTGTATGCCGAATGGTCAGAAGTGGTTAGCACTAATTTTCGCGATGCCAGTGGCCCGGCGATGACGGAGATTGGAGAACAGCCGTGGGGTCGCGAATTTGCGTTGCGCGATCCGGCGGGTAACTGCGTCCATTTTGTCGCAGAAGAGCAGGATTGA

>Sequence24

ATGGCGAAACTGACCAGCGCGGTCCCGGTACTGACCGCCCGTGATGTTGCGGGTGCGGTTGAATTTTGGACCGATCGTCTGGGGTTTTCACGCGATTTCGTTGAAGATGATTTTGCGGGTGTAGTGCGCGATGACGTGACGCTGTTCATTAGCGCAGTGCAGGACCAGGTTGTCCCGGATAATACCCTGGCTTGGGTATGGGTGCGTGGCCTGGATGAACTGTATGCCGAGTGGTCGGAAGTGGTGAGCACCAACTTTCGCGATGCAAGCGGCCCGGCCATGACGGAGATCGGCGAACAGCCGTGGGGCCGCGAATTCGCCTTACGCGATCCGGCCGGAAACTGCGTCCATTTTGTTGCCGAAGAACAGGACTAA

>Sequence25

ATGGCCAAACTGACCTCGGCCGTTCCGGTACTTACCGCCCGTGATGTGGCCGGGGCGGTTGAATTTTGGACCGACCGCCTGGGTTTTTCACGCGATTTCGTGGAGGACGATTTCGCGGGCGTGGTGCGCGATGACGTGACGTTGTTTATTAGCGCAGTGCAGGATCAGGTTGTTCCGGATAACACCCTGGCGTGGGTGTGGGTACGTGGTCTGGATGAACTGTATGCGGAATGGAGCGAAGTGGTCAGCACCAACTTTCGTGATGCTAGTGGCCCGGCGATGACTGAGATTGGCGAACAGCCGTGGGGTCGCGAATTCGCTCTGCGCGATCCGGCGGGCAACTGCGTGCATTTTGTGGCGGAAGAGCAGGATTGA

>Sequence26

ATGGCGAAATTGACGTCAGCCGTGCCGGTGCTGACTGCGCGCGATGTGGCGGGTGCCGTGGAGTTCTGGACCGACCGCCTTGGTTTTAGCCGCGATTTCGTGGAAGATGATTTTGCCGGAGTAGTTCGTGATGATGTCACCCTCTTTATTAGCGCCGTTCAGGATCAGGTGGTTCCGGATAACACGCTGGCGTGGGTGTGGGTCCGTGGTCTGGACGAACTGTATGCCGAATGGAGCGAAGTTGTGAGCACCAACTTTCGCGACGCGAGCGGCCCTGCCATGACCGAAATCGGCGAACAGCCGTGGGGCCGCGAATTTGCTTTACGTGATCCGGCGGGCAATTGCGTGCATTTTGTGGCCGAAGAACAGGACTAA

>Sequence27

ATGGCCAAACTGACGAGCGCGGTACCGGTTCTTACAGCTCGCGATGTTGCGGGTGCCGTGGAATTTTGGACCGATCGTCTCGGATTCTCGCGTGATTTTGTGGAAGACGATTTTGCGGGCGTAGTGCGCGATGATGTCACCCTGTTTATTAGCGCGGTTCAGGATCAGGTGGTTCCGGATAACACCTTGGCGTGGGTGTGGGTGCGCGGCCTGGATGAGCTGTATGCCGAGTGGAGCGAAGTGGTTAGCACCAACTTTCGTGATGCAAGTGGCCCGGCCATGACTGAAATTGGCGAACAGCCGTGGGGCCGCGAATTTGCGTTACGCGATCCGGCCGGTAATTGCGTGCATTTCGTCGCAGAGGAACAGGATTAA

>Sequence28

ATGGCGAAACTGACGAGCGCGGTTCCGGTGCTGACCGCGCGTGATGTGGCCGGCGCCGTGGAATTTTGGACCGATCGCCTGGGCTTTAGCCGTGATTTTGTCGAAGACGATTTCGCCGGCGTTGTTCGCGATGACGTGACCCTGTTTATTAGCGCAGTCCAGGATCAGGTAGTGCCTGATAACACGTTAGCGTGGGTATGGGTGCGCGGTCTGGATGAACTGTATGCAGAATGGAGTGAAGTCGTATCGACCAACTTCCGTGACGCCAGCGGCCCGGCGATGACTGAAATTGGCGAGCAGCCGTGGGGTCGCGAATTTGCCCTGCGTGACCCGGCGGGGAATTGCGTGCATTTCGTTGCAGAAGAACAGGACTAA

>Sequence29

ATGGCGAAACTGACCAGCGCGGTACCGGTTCTGACGGCTCGTGACGTGGCCGGCGCGGTTGAATTTTGGACCGATCGCCTGGGCTTTAGCCGCGACTTTGTAGAAGACGATTTCGCCGGTGTGGTTCGCGATGATGTTACGCTTTTTATTTCGGCCGTACAGGATCAGGTGGTGCCGGACAACACCCTCGCGTGGGTGTGGGTTCGTGGTCTGGACGAACTGTATGCGGAATGGAGCGAGGTGGTCAGTACCAATTTTCGTGATGCCTCAGGCCCGGCGATGACCGAAATTGGCGAACAGCCTTGGGGGCGCGAATTCGCGCTGCGCGATCCGGCAGGTAACTGCGTGCATTTCGTGGCAGAAGAACAGGATTAA

>Sequence30

ATGGCAAAACTGACCAGCGCAGTCCCGGTTTTAACCGCGCGCGATGTGGCGGGCGCCGTGGAATTCTGGACTGATCGTCTGGGCTTTAGCCGCGACTTCGTTGAAGATGATTTCGCAGGCGTTGTGCGTGATGATGTCACCCTTTTTATTAGCGCGGTGCAGGATCAGGTCGTTCCGGATAACACCCTGGCGTGGGTGTGGGTTCGCGGTCTGGATGAACTGTATGCCGAGTGGAGCGAAGTAGTATCGACCAATTTTCGTGACGCCAGCGGTCCGGCCATGACCGAAATTGGCGAACAGCCGTGGGGACGCGAGTTTGCGCTGCGCGATCCGGCCGGTAACTGCGTGCATTTTGTGGCTGAGGAACAGGACTGA

>Sequence31

ATGGCCAAACTGACAAGCGCGGTTCCGGTGTTGACGGCCCGCGATGTGGCCGGTGCGGTTGAATTCTGGACCGATCGCCTGGGTTTTAGCCGCGATTTTGTTGAGGATGACTTCGCGGGAGTTGTGCGCGACGATGTTACCCTGTTTATCTCAGCAGTACAGGATCAGGTCGTGCCGGATAACACCCTTGCGTGGGTATGGGTTCGTGGCCTGGATGAACTGTATGCTGAATGGAGTGAAGTGGTGTCGACCAACTTTCGTGATGCGAGCGGCCCGGCTATGACCGAAATCGGTGAACAGCCTTGGGGCCGCGAATTTGCGCTGCGTGACCCGGCGGGCAACTGCGTCCATTTTGTGGCCGAAGAACAGGATTAA

>Sequence32

ATGGCCAAACTGACAAGCGCGGTGCCGGTGCTTACCGCGCGCGATGTAGCGGGCGCTGTGGAATTCTGGACCGACCGCCTGGGCTTTAGCCGCGATTTTGTTGAAGATGATTTTGCGGGCGTGGTTCGTGACGATGTGACCCTGTTTATCAGTGCCGTGCAGGATCAAGTGGTTCCGGATAACACCCTGGCCTGGGTTTGGGTGCGTGGCTTAGATGAGCTGTATGCGGAATGGTCGGAAGTGGTCTCAACCAACTTCCGCGACGCCAGCGGCCCGGCGATGACCGAAATCGGCGAACAGCCTTGGGGTCGTGAGTTTGCTCTGCGTGATCCGGCGGGTAACTGCGTGCATTTTGTAGCGGAAGAACAGGACTGA

>Sequence33

ATGGCTAAACTGACGAGCGCCGTGCCGGTCCTCACAGCACGTGATGTTGCAGGCGCGGTGGAGTTTTGGACCGATCGCCTGGGTTTTAGCCGTGACTTCGTTGAAGATGATTTCGCGGGCGTGGTGCGTGATGATGTGACCCTTTTTATTTCAGCCGTTCAGGATCAAGTGGTTCCGGACAATACGCTGGCATGGGTGTGGGTTCGCGGCCTGGATGAACTGTATGCCGAATGGAGCGAGGTTGTTAGTACCAACTTTCGCGATGCGTCGGGTCCTGCGATGACTGAAATTGGCGAACAGCCGTGGGGCCGCGAATTTGCTCTGCGTGATCCGGCCGGCAACTGCGTGCATTTTGTGGCGGAAGAACAGGATTAA

>Sequence34

ATGGCAAAACTCACCAGCGCGGTTCCGGTGCTGACCGCCCGTGATGTAGCCGGCGCCGTGGAATTTTGGACTGATCGCCTGGGGTTCAGCCGCGACTTTGTTGAAGATGATTTTGCGGGCGTTGTTCGCGATGATGTGACATTGTTCATTAGTGCGGTGCAGGATCAAGTGGTCCCGGATAACACCCTTGCGTGGGTGTGGGTACGTGGTTTAGATGAACTGTATGCGGAATGGTCAGAAGTGGTTAGCACGAACTTTCGCGATGCATCGGGCCCGGCAATGACCGAGATCGGCGAACAGCCTTGGGGTCGCGAATTTGCGCTGCGTGATCCGGCGGGCAATTGCGTCCATTTTGTGGCCGAAGAACAGGATTAA

>Sequence35

ATGGCCAAACTGACCTCGGCCGTTCCGGTTTTAACTGCCCGTGATGTGGCCGGAGCCGTAGAATTCTGGACCGATCGCCTGGGCTTTAGCCGCGATTTCGTGGAAGACGACTTCGCAGGGGTAGTGCGTGATGATGTAACACTGTTTATTAGCGCGGTTCAAGATCAGGTGGTGCCGGATAACACCCTGGCGTGGGTTTGGGTTCGTGGTCTTGATGAATTGTATGCCGAATGGAGCGAGGTTGTGTCAACCAACTTTCGCGATGCGAGCGGTCCGGCCATGACCGAAATTGGCGAGCAGCCGTGGGGCCGCGAGTTTGCGCTGCGTGATCCGGCGGGCAACTGCGTGCATTTTGTGGCGGAAGAACAGGATTAA

>Sequence36

ATGGCGAAACTTACGAGTGCGGTCCCGGTTCTGACCGCCCGCGATGTGGCGGGCGCGGTGGAGTTTTGGACCGACCGCCTGGGGTTCAGCCGCGATTTCGTGGAAGATGACTTTGCCGGCGTGGTTCGTGATGATGTGACCTTATTTATCAGCGCTGTCCAAGACCAGGTGGTTCCGGATAACACCCTGGCCTGGGTGTGGGTACGCGGTCTGGATGAACTGTATGCGGAGTGGAGCGAAGTCGTGTCAACCAACTTTCGCGATGCCAGCGGCCCGGCCATGACGGAAATTGGCGAACAGCCGTGGGGTCGTGAGTTCGCGCTGCGTGATCCGGCGGGCAACTGCGTTCATTTTGTTGCCGAAGAACAGGATTGA

>Sequence37

ATGGCCAAACTGACCAGCGCGGTGCCGGTGCTGACTGCGCGCGACGTGGCGGGCGCTGTTGAATTTTGGACCGATCGTCTGGGTTTCAGCCGCGATTTTGTGGAAGATGACTTTGCCGGCGTCGTACGTGATGATGTTACGCTGTTTATTAGCGCCGTTCAGGACCAGGTTGTACCGGACAACACCTTGGCGTGGGTCTGGGTGCGCGGACTGGATGAGTTATATGCAGAATGGAGCGAAGTGGTGTCGACCAACTTTCGCGATGCCAGTGGCCCGGCGATGACCGAAATCGGCGAACAGCCGTGGGGTCGTGAATTTGCGCTCCGTGATCCGGCTGGCAACTGCGTACATTTCGTGGCAGAAGAACAGGACTAA

>Sequence38

ATGGCCAAACTGACCAGCGCCGTACCTGTGCTGACGGCCCGCGATGTGGCGGGCGCCGTTGAATTTTGGACCGACCGTCTGGGCTTTTCGCGTGATTTCGTGGAAGATGATTTCGCGGGCGTGGTGCGCGATGATGTGACTCTGTTTATTAGCGCCGTTCAAGATCAGGTTGTGCCGGACAACACCCTGGCGTGGGTGTGGGTTCGCGGGCTGGACGAACTGTATGCGGAGTGGAGCGAAGTAGTTAGCACCAATTTCCGCGATGCGTCAGGCCCGGCCATGACGGAAATTGGCGAACAGCCGTGGGGTCGTGAATTTGCGCTCCGCGATCCGGCCGGTAACTGCGTCCATTTTGTAGCAGAAGAGCAGGACTGA

>Sequence39

ATGGCCAAACTGACGAGCGCTGTGCCGGTACTCACCGCCCGTGACGTGGCGGGCGCGGTTGAGTTTTGGACCGATCGCCTTGGCTTTTCGCGTGATTTTGTGGAAGATGATTTCGCGGGGGTTGTGCGTGACGACGTCACCCTGTTTATCAGCGCGGTGCAGGATCAGGTGGTTCCGGATAATACCCTGGCGTGGGTGTGGGTGCGCGGCCTGGATGAACTGTATGCCGAGTGGAGCGAAGTTGTGAGCACCAACTTTCGTGACGCCAGCGGTCCGGCTATGACTGAAATTGGTGAACAACCGTGGGGTCGCGAATTTGCCCTGCGCGATCCTGCCGGCAACTGCGTGCATTTCGTGGCGGAAGAACAGGATTGA

>Sequence40

ATGGCGAAACTGACTTCGGCCGTACCGGTGCTGACCGCGCGCGATGTGGCCGGTGCGGTTGAATTTTGGACCGATCGTCTGGGGTTTAGTCGCGACTTTGTGGAAGATGATTTTGCGGGCGTGGTGCGCGACGATGTTACCCTTTTTATCAGCGCTGTACAGGACCAGGTTGTCCCGGATAACACCTTGGCATGGGTGTGGGTGCGTGGTCTCGATGAACTGTATGCGGAATGGAGCGAAGTGGTTAGCACGAATTTCCGTGATGCAAGCGGCCCGGCGATGACGGAGATCGGCGAACAACCGTGGGGCCGTGAATTTGCCCTGCGCGATCCGGCCGGAAACTGCGTGCATTTCGTGGCCGAAGAACAGGATTAA

>Sequence41

ATGGCCAAACTGACCAGCGCAGTGCCGGTCCTCACCGCGCGCGACGTTGCCGGCGCCGTGGAGTTTTGGACCGATCGTCTGGGCTTTAGCCGCGATTTTGTGGAAGATGATTTTGCCGGCGTGGTTCGCGATGATGTGACGCTTTTTATTTCGGCGGTTCAGGATCAGGTGGTACCGGATAACACGCTGGCCTGGGTTTGGGTGCGTGGCTTAGACGAACTGTATGCCGAATGGAGCGAAGTGGTTAGCACCAATTTTCGCGATGCGTCAGGTCCGGCAATGACTGAAATTGGTGAACAACCGTGGGGCCGTGAATTCGCGCTGCGTGATCCTGCGGGAAACTGCGTACATTTCGTTGCGGAAGAGCAGGATTGA

>Sequence42

ATGGCGAAATTGACGTCGGCCGTTCCGGTGCTGACCGCACGCGATGTTGCGGGCGCCGTCGAGTTTTGGACCGATCGTCTGGGCTTCAGCCGCGATTTTGTTGAAGACGATTTTGCGGGCGTCGTGCGTGATGATGTTACCCTGTTCATTAGTGCAGTGCAAGATCAGGTTGTGCCGGATAACACACTGGCGTGGGTGTGGGTGCGTGGTCTGGATGAACTGTATGCTGAATGGAGCGAAGTTGTGAGCACCAACTTCCGCGACGCCAGCGGTCCGGCGATGACGGAAATCGGCGAACAGCCGTGGGGCCGCGAATTTGCATTACGTGACCCGGCTGGCAACTGCGTTCATTTTGTCGCCGAAGAGCAGGATTAA

>Sequence43

ATGGCCAAACTGACCAGTGCCGTCCCTGTATTGACAGCACGCGACGTGGCGGGCGCGGTTGAATTTTGGACCGATCGCCTGGGCTTCAGCCGCGATTTTGTCGAGGACGATTTTGCCGGTGTTGTTCGCGATGATGTGACCCTGTTTATTAGCGCCGTGCAGGATCAAGTGGTGCCGGACAATACCCTCGCCTGGGTTTGGGTGCGTGGCCTGGATGAACTGTATGCGGAATGGAGCGAAGTTGTGAGCACGAACTTCCGTGATGCGTCAGGGCCGGCTATGACCGAAATCGGCGAACAGCCGTGGGGCCGTGAATTCGCGCTGCGTGATCCGGCGGGTAACTGCGTGCATTTTGTGGCTGAGGAACAGGATTAA

>Sequence44

ATGGCGAAACTCACCAGCGCCGTGCCGGTTCTGACCGCCCGCGATGTCGCGGGCGCGGTTGAATTTTGGACGGATCGTTTGGGCTTTAGCCGTGATTTTGTGGAGGATGATTTCGCGGGTGTAGTGCGCGATGACGTTACTCTGTTTATCAGCGCAGTGCAGGACCAGGTAGTGCCGGACAATACGTTAGCCTGGGTGTGGGTGCGCGGTCTTGATGAACTGTATGCTGAATGGAGCGAAGTGGTTAGCACCAACTTTCGTGATGCCTCAGGCCCGGCGATGACCGAAATCGGCGAACAGCCGTGGGGTCGCGAATTTGCCCTGCGCGATCCGGCGGGGAACTGCGTGCATTTCGTGGCCGAAGAGCAGGACTAA

>Sequence45

ATGGCCAAACTGACGAGTGCGGTACCGGTGCTGACCGCGCGTGACGTAGCGGGCGCGGTGGAATTTTGGACCGATCGTCTGGGTTTTTCGCGCGACTTTGTTGAAGATGATTTTGCCGGTGTGGTGCGCGATGACGTGACCCTGTTTATCAGCGCGGTTCAGGATCAGGTCGTGCCGGATAACACGCTGGCGTGGGTTTGGGTGCGTGGTTTAGATGAACTGTATGCCGAATGGTCAGAAGTTGTTAGCACTAACTTTCGTGATGCAAGCGGTCCTGCAATGACCGAGATTGGCGAGCAACCGTGGGGCCGCGAATTCGCCCTGCGCGACCCGGCTGGCAATTGCGTCCATTTCGTGGCGGAAGAACAGGATTAA

>Sequence46

ATGGCCAAACTGACCAGTGCCGTTCCGGTGCTTACCGCGCGCGATGTTGCCGGCGCGGTGGAATTTTGGACTGATCGTCTGGGCTTTAGCCGTGATTTTGTGGAGGATGACTTCGCTGGCGTTGTGCGCGATGATGTGACCCTGTTTATCAGCGCGGTGCAGGATCAAGTGGTGCCGGACAACACGCTGGCCTGGGTTTGGGTGCGTGGTCTGGATGAGTTGTATGCGGAATGGAGCGAAGTTGTTAGCACGAACTTTCGCGATGCCTCGGGTCCGGCCATGACCGAAATTGGAGAACAGCCTTGGGGCCGCGAATTTGCGTTACGCGACCCGGCCGGTAATTGCGTACATTTTGTAGCGGAAGAACAGGACTAA

>Sequence47

ATGGCGAAATTAACGAGCGCGGTGCCGGTGCTGACCGCGCGCGATGTGGCCGGCGCCGTGGAATTCTGGACAGATCGCCTGGGCTTTAGCCGCGATTTTGTTGAAGATGATTTTGCCGGGGTGGTACGCGATGACGTTACTCTGTTTATCAGCGCAGTTCAGGATCAGGTGGTCCCGGATAATACGCTGGCGTGGGTTTGGGTTCGTGGTCTGGACGAACTGTATGCAGAATGGAGCGAAGTGGTTTCGACCAACTTTCGTGATGCGTCAGGTCCGGCTATGACCGAGATTGGCGAACAGCCGTGGGGCCGTGAGTTCGCCCTGCGTGACCCTGCGGGTAACTGCGTACATTTTGTGGCAGAAGAACAGGACTAA

>Sequence48

ATGGCTAAATTGACCAGTGCGGTTCCGGTTCTGACCGCGCGTGATGTAGCCGGCGCAGTCGAATTCTGGACCGATCGCTTAGGCTTTTCGCGCGATTTTGTGGAAGATGATTTTGCGGGCGTTGTCCGTGACGACGTGACCCTGTTTATTAGCGCCGTGCAGGATCAGGTGGTTCCGGATAATACGCTGGCGTGGGTGTGGGTACGCGGTCTGGATGAACTCTATGCTGAATGGAGCGAAGTGGTATCAACTAACTTTCGTGACGCAAGCGGCCCGGCGATGACCGAAATTGGTGAACAACCGTGGGGCCGTGAGTTCGCCCTGCGCGATCCGGCCGGCAACTGCGTGCATTTTGTGGCAGAAGAGCAGGACTAA

>Sequence49

ATGGCGAAACTGACCTCGGCAGTCCCGGTGCTCACCGCGCGCGATGTGGCCGGCGCTGTTGAGTTTTGGACTGATCGTCTGGGTTTTAGCCGCGATTTTGTGGAAGATGATTTTGCTGGCGTGGTACGTGATGACGTTACCCTTTTTATTAGTGCGGTACAAGATCAGGTGGTGCCGGATAACACCCTGGCGTGGGTTTGGGTTCGTGGTTTGGACGAATTATATGCCGAATGGAGCGAGGTGGTGAGCACCAATTTTCGCGATGCATCAGGCCCGGCGATGACAGAAATTGGTGAGCAGCCGTGGGGACGCGAATTTGCCCTGCGCGATCCGGCCGGCAACTGCGTGCATTTTGTCGCGGAAGAACAGGATTAA

>Sequence50

ATGGCAAAACTGACCAGTGCGGTTCCGGTTTTGACGGCGCGCGATGTCGCCGGTGCCGTGGAATTCTGGACGGATCGCCTGGGCTTTAGCCGCGATTTCGTTGAGGATGATTTTGCGGGTGTGGTGCGTGATGATGTGACCCTGTTTATCTCAGCAGTTCAGGACCAGGTTGTTCCGGACAATACCCTGGCCTGGGTTTGGGTGCGCGGCTTAGACGAACTTTATGCCGAATGGAGCGAAGTGGTAAGCACCAACTTTCGTGATGCGTCGGGTCCGGCTATGACAGAAATTGGCGAACAGCCTTGGGGCCGTGAATTTGCGCTGCGCGACCCGGCGGGCAACTGCGTGCATTTTGTTGCGGAAGAGCAGGATTAA

>Sequence51

ATGGCCAAACTGACCTCAGCGGTTCCGGTGCTGACCGCCCGCGACGTGGCGGGAGCGGTTGAATTTTGGACCGACCGCCTGGGTTTCAGCCGCGACTTTGTGGAAGATGACTTTGCGGGCGTGGTTCGTGATGATGTGACACTGTTTATTAGTGCTGTTCAAGATCAGGTCGTACCGGATAACACCCTGGCGTGGGTCTGGGTACGTGGCCTGGATGAACTGTATGCCGAGTGGAGCGAGGTGGTGAGCACCAATTTTCGTGATGCAAGCGGCCCGGCTATGACCGAAATTGGCGAACAGCCGTGGGGTCGCGAATTCGCCCTCCGTGATCCGGCGGGGAACTGCGTACATTTCGTTGCAGAAGAACAGGATTGA

>Sequence52

ATGGCTAAACTGACGAGCGCGGTTCCGGTGTTAACCGCGCGCGATGTTGCAGGCGCAGTGGAATTTTGGACCGACCGCCTGGGATTTAGTCGTGACTTCGTCGAAGATGATTTCGCGGGTGTAGTTCGTGATGATGTAACCCTGTTTATCTCAGCAGTGCAGGATCAAGTGGTTCCGGATAACACCCTTGCCTGGGTGTGGGTTCGTGGCCTGGATGAACTGTATGCCGAATGGAGCGAAGTGGTGAGCACTAACTTTCGCGATGCCTCGGGTCCGGCGATGACCGAGATTGGCGAGCAGCCGTGGGGCCGCGAATTTGCCCTGCGCGATCCGGCGGGGAACTGCGTACATTTTGTCGCCGAAGAACAGGACTAA

>Sequence53

ATGGCCAAACTGACCAGCGCAGTTCCGGTGCTGACCGCCCGTGATGTCGCGGGCGCCGTTGAATTTTGGACGGATCGTCTGGGCTTTAGCCGCGATTTCGTTGAAGATGACTTCGCGGGTGTCGTGCGCGATGACGTCACGCTGTTTATCTCGGCAGTTCAGGATCAGGTGGTGCCGGACAATACTCTGGCATGGGTATGGGTTCGCGGCCTGGACGAACTGTATGCCGAATGGTCAGAAGTGGTGAGTACCAACTTTCGCGATGCGAGCGGCCCGGCGATGACCGAAATTGGCGAACAGCCGTGGGGCCGTGAGTTCGCTCTCCGCGACCCGGCGGGTAACTGCGTACATTTTGTTGCGGAAGAGCAGGATTGA

>Sequence54

ATGGCCAAACTGACGAGCGCCGTGCCTGTTTTAACCGCGCGTGATGTTGCCGGAGCGGTGGAATTCTGGACCGATCGTCTGGGCTTTAGTCGCGACTTTGTAGAGGATGATTTTGCAGGCGTGGTACGCGATGATGTTACCTTGTTTATTAGCGCAGTGCAGGACCAGGTTGTGCCGGACAACACCCTTGCGTGGGTCTGGGTTCGCGGCCTGGACGAACTGTATGCGGAATGGAGCGAGGTAGTGAGCACGAACTTTCGCGATGCGAGCGGTCCGGCTATGACCGAAATCGGCGAACAGCCGTGGGGTCGTGAATTTGCTCTGCGTGATCCGGCCGGTAACTGCGTTCATTTTGTGGCCGAAGAACAGGATTAA

>Sequence55

ATGGCAAAACTGACCTCAGCCGTGCCGGTGCTGACAGCGCGCGACGTGGCCGGCGCTGTTGAATTCTGGACCGATCGCCTGGGCTTTAGCCGTGATTTTGTAGAAGATGATTTTGCCGGCGTGGTGCGTGATGATGTGACGTTATTCATCAGCGCCGTGCAGGATCAGGTGGTCCCGGATAATACCCTGGCCTGGGTTTGGGTACGCGGTTTGGATGAACTGTATGCGGAGTGGAGTGAAGTGGTTAGCACCAACTTTCGTGATGCGTCGGGTCCGGCGATGACCGAAATCGGCGAACAACCGTGGGGCCGCGAGTTTGCTCTGCGTGACCCGGCCGGAAACTGCGTGCATTTCGTTGCGGAAGAGCAGGACTAA

>Sequence56

ATGGCGAAACTCACCAGCGCCGTTCCGGTTCTTACCGCGCGCGATGTGGCCGGCGCGGTGGAGTTTTGGACCGACCGCCTGGGTTTCAGTCGCGATTTTGTTGAAGATGACTTCGCGGGTGTCGTGCGTGACGATGTTACCCTGTTTATCTCGGCTGTGCAGGACCAGGTTGTTCCGGATAACACCCTGGCATGGGTTTGGGTGCGTGGACTGGATGAACTGTATGCCGAATGGAGCGAGGTCGTGTCAACCAACTTTCGCGATGCAAGCGGCCCGGCGATGACGGAAATTGGCGAGCAGCCGTGGGGGCGCGAATTTGCGCTGCGTGATCCGGCCGGCAACTGCGTACATTTTGTGGCAGAAGAACAGGATTGA

>Sequence57

ATGGCTAAACTTACGAGCGCCGTACCGGTGCTCACCGCGCGCGACGTGGCGGGCGCCGTTGAGTTTTGGACCGACCGTCTGGGTTTTAGCCGCGACTTTGTTGAAGATGATTTCGCGGGTGTGGTGCGTGATGATGTTACCTTATTTATTAGCGCTGTCCAAGATCAGGTCGTGCCTGATAATACCCTGGCCTGGGTCTGGGTACGTGGTCTGGACGAATTGTATGCGGAATGGTCGGAAGTGGTGAGTACCAACTTTCGCGATGCGAGCGGCCCGGCGATGACCGAAATCGGCGAACAGCCGTGGGGTCGCGAATTTGCCCTGCGCGACCCGGCCGGCAACTGCGTGCATTTCGTTGCGGAAGAACAGGATTAA

>Sequence58

ATGGCTAAATTAACGAGCGCGGTGCCTGTTCTGACCGCCCGCGATGTAGCCGGCGCGGTGGAGTTTTGGACCGACCGCTTGGGTTTTAGCCGCGATTTTGTGGAAGATGATTTTGCGGGCGTTGTGCGTGATGATGTGACTCTCTTCATCAGCGCGGTACAGGATCAGGTAGTCCCGGACAACACGCTGGCGTGGGTTTGGGTGCGTGGTCTGGATGAACTGTATGCCGAGTGGAGCGAAGTGGTTTCAACCAATTTTCGTGATGCGAGTGGGCCGGCGATGACCGAAATTGGCGAGCAGCCGTGGGGTCGCGAATTCGCCCTGCGCGACCCGGCCGGCAACTGCGTTCATTTTGTCGCGGAAGAACAGGATTAA

>Sequence59

ATGGCCAAACTTACCAGCGCAGTTCCGGTGCTGACGGCCCGCGATGTGGCGGGCGCGGTGGAGTTCTGGACCGATCGTCTGGGCTTTTCGCGCGACTTTGTTGAAGATGACTTTGCGGGCGTGGTGCGTGATGACGTTACCCTGTTCATTTCAGCGGTGCAGGATCAAGTTGTTCCGGACAACACCTTAGCGTGGGTGTGGGTGCGCGGGCTGGATGAGCTCTATGCGGAATGGAGCGAAGTGGTGAGCACGAATTTTCGCGATGCAAGCGGCCCGGCCATGACAGAAATCGGAGAACAGCCTTGGGGTCGTGAATTTGCGCTGCGTGATCCGGCCGGTAACTGCGTCCATTTTGTGGCCGAAGAACAGGATTGA

>Sequence60

ATGGCCAAACTGACCAGCGCAGTGCCGGTACTTACCGCTCGCGATGTGGCAGGTGCCGTGGAATTTTGGACCGATCGTCTCGGCTTTAGCCGTGACTTCGTTGAAGACGATTTTGCCGGCGTTGTGCGTGACGATGTTACGCTGTTTATTAGCGCGGTGCAAGACCAGGTTGTGCCGGATAACACGCTGGCCTGGGTATGGGTCCGCGGCCTGGACGAACTGTATGCCGAATGGTCGGAGGTCGTGAGCACTAACTTTCGTGATGCGAGCGGCCCGGCGATGACCGAGATCGGCGAACAGCCGTGGGGCCGCGAATTCGCGCTGCGCGATCCGGCAGGTAACTGCGTCCATTTTGTTGCGGAAGAACAGGATTGA

>Sequence61

ATGGCGAAACTTACCAGTGCCGTGCCGGTCCTGACGGCTCGCGATGTTGCCGGTGCGGTGGAATTTTGGACTGATCGCTTAGGCTTTAGCCGCGATTTCGTGGAAGATGACTTTGCAGGAGTGGTCCGCGACGATGTGACCCTGTTCATTAGCGCCGTGCAGGATCAGGTGGTGCCGGATAACACGCTGGCATGGGTTTGGGTACGTGGCCTGGACGAACTGTATGCGGAATGGTCGGAAGTTGTGAGCACCAACTTTCGTGACGCGAGCGGTCCTGCGATGACCGAAATCGGCGAGCAGCCGTGGGGCCGTGAATTTGCACTCCGCGATCCGGCGGGCAACTGCGTTCATTTTGTCGCCGAAGAGCAGGATTGA

>Sequence62

ATGGCCAAACTGACCAGCGCGGTTCCGGTCCTGACCGCGCGCGACGTTGCAGGCGCCGTGGAATTCTGGACCGACCGTTTAGGTTTTAGCCGTGATTTCGTTGAAGATGACTTCGCGGGTGTGGTTCGTGATGATGTCACCTTGTTTATTAGTGCGGTGCAGGATCAGGTTGTACCGGACAACACCCTGGCCTGGGTGTGGGTGCGCGGCCTGGATGAGCTCTATGCCGAATGGAGCGAGGTAGTAAGCACCAACTTTCGTGATGCCAGCGGGCCTGCGATGACAGAAATTGGCGAACAGCCGTGGGGCCGCGAATTTGCGCTGCGCGATCCGGCTGGCAACTGCGTCCATTTTGTGGCCGAAGAACAGGACTGA

>Sequence63

ATGGCGAAACTGACCAGCGCCGTTCCGGTCCTGACTGCCCGCGATGTGGCGGGCGCGGTCGAATTTTGGACCGATCGTCTGGGCTTTAGCCGCGATTTTGTGGAAGACGACTTCGCCGGTGTGGTTCGCGACGATGTCACCTTGTTTATCAGCGCAGTTCAGGATCAGGTTGTGCCGGATAACACCCTGGCGTGGGTTTGGGTGCGTGGCCTGGATGAGCTGTATGCGGAATGGAGCGAAGTGGTGTCGACCAACTTTCGCGATGCGAGCGGTCCGGCCATGACGGAAATCGGGGAGCAGCCGTGGGGTCGCGAGTTCGCCCTCCGTGATCCTGCAGGCAACTGCGTTCATTTTGTAGCGGAAGAACAAGATTGA

>Sequence64

ATGGCCAAACTGACGTCGGCCGTGCCGGTTCTGACCGCACGTGATGTTGCGGGTGCGGTGGAATTCTGGACAGATCGCTTGGGCTTTAGCCGCGACTTTGTAGAAGATGACTTCGCAGGTGTGGTGCGCGATGATGTGACCCTCTTTATTTCAGCGGTGCAGGATCAGGTTGTGCCGGATAACACGCTGGCGTGGGTCTGGGTGCGTGGCCTGGACGAACTGTATGCAGAATGGAGCGAAGTAGTTAGTACTAATTTTCGCGATGCCAGCGGTCCTGCCATGACCGAGATTGGCGAACAGCCGTGGGGCCGTGAGTTTGCGCTGCGCGATCCGGCCGGTAACTGCGTGCATTTTGTCGCCGAAGAGCAAGATTAA

>Sequence65

ATGGCGAAACTGACCAGCGCCGTACCGGTGCTGACCGCGCGTGACGTTGCGGGCGCCGTGGAATTTTGGACCGATCGTCTGGGCTTTAGCCGCGATTTTGTTGAGGATGACTTTGCGGGCGTTGTCCGCGATGATGTGACGCTGTTTATCAGCGCCGTGCAGGATCAAGTGGTACCTGATAACACCCTGGCTTGGGTGTGGGTTCGTGGTTTGGATGAACTGTATGCGGAATGGAGTGAAGTCGTGAGCACGAATTTCCGCGATGCCTCAGGGCCGGCAATGACCGAAATCGGCGAACAGCCGTGGGGTCGCGAATTTGCGCTCCGTGACCCGGCGGGCAACTGCGTTCATTTTGTTGCGGAGGAGCAGGACTAA

>Sequence66

ATGGCAAAACTGACCAGCGCCGTTCCGGTCCTGACCGCGCGCGATGTGGCCGGCGCGGTTGAATTTTGGACCGATCGCTTAGGCTTTTCACGTGATTTTGTGGAAGATGACTTTGCAGGTGTTGTACGTGACGATGTGACTCTGTTTATCAGCGCGGTTCAGGATCAGGTCGTGCCGGACAACACGTTGGCGTGGGTGTGGGTCCGCGGTCTTGACGAGCTGTATGCGGAATGGAGCGAGGTTGTGTCGACGAACTTTCGTGATGCTAGCGGCCCGGCCATGACCGAGATTGGCGAACAGCCGTGGGGCCGCGAATTCGCCCTGCGCGATCCGGCCGGCAACTGCGTGCATTTTGTTGCGGAAGAACAAGATTGA

>Sequence67

ATGGCTAAACTGACCAGTGCCGTGCCGGTTTTAACGGCCCGCGATGTTGCGGGCGCGGTTGAATTTTGGACCGATCGCCTGGGCTTTTCACGCGATTTTGTGGAAGATGATTTCGCCGGTGTAGTTCGCGATGACGTGACCCTGTTTATTAGCGCGGTCCAGGATCAGGTCGTACCGGACAATACCCTGGCATGGGTGTGGGTGCGTGGTCTGGACGAACTTTATGCCGAGTGGAGCGAAGTTGTGAGCACTAACTTTCGTGACGCGAGCGGTCCTGCCATGACCGAAATTGGCGAGCAGCCGTGGGGTCGCGAATTTGCGTTGCGTGATCCGGCCGGCAACTGCGTGCATTTTGTAGCGGAAGAGCAAGATTAA

>Sequence68

ATGGCGAAACTGACCAGCGCCGTTCCGGTGCTGACCGCACGCGATGTCGCCGGCGCCGTTGAATTTTGGACGGATCGCCTGGGCTTCAGCCGCGACTTTGTTGAAGATGATTTTGCTGGAGTGGTGCGTGACGATGTGACTCTGTTTATTAGCGCAGTGCAGGATCAGGTTGTGCCGGACAACACCCTGGCTTGGGTGTGGGTTCGCGGCTTAGACGAACTCTATGCGGAATGGAGTGAGGTCGTGAGCACGAACTTTCGTGATGCGTCGGGTCCGGCGATGACCGAAATTGGCGAACAACCGTGGGGTCGTGAGTTTGCGCTGCGCGATCCGGCAGGCAACTGCGTACATTTTGTCGCCGAGGAACAGGACTAA

>Sequence69

ATGGCGAAACTCACGAGCGCCGTACCTGTTTTGACCGCCCGCGATGTTGCGGGCGCCGTCGAGTTCTGGACCGACCGTCTGGGCTTTAGTCGTGATTTTGTGGAAGACGATTTTGCGGGTGTTGTACGCGATGATGTTACGCTGTTTATTAGCGCCGTTCAGGATCAGGTGGTGCCGGATAACACCCTGGCGTGGGTGTGGGTACGCGGTCTGGACGAACTGTATGCGGAATGGAGCGAAGTGGTGAGCACTAATTTCCGTGATGCGTCAGGTCCGGCCATGACCGAGATTGGCGAACAGCCGTGGGGCCGTGAATTCGCACTGCGCGATCCGGCGGGTAACTGCGTGCATTTTGTGGCGGAAGAGCAAGACTGA

>Sequence70

ATGGCGAAACTTACCAGTGCCGTGCCGGTCCTGACCGCGCGTGATGTGGCGGGTGCTGTTGAATTTTGGACAGATCGCCTGGGCTTCAGCCGCGACTTTGTCGAAGACGATTTTGCAGGCGTGGTTCGTGATGATGTGACTCTGTTCATTTCAGCCGTTCAGGATCAGGTGGTGCCGGACAACACCCTCGCGTGGGTGTGGGTCCGTGGCCTGGATGAACTGTATGCGGAGTGGAGCGAAGTGGTAAGCACGAACTTCCGCGACGCAAGCGGTCCGGCAATGACCGAAATTGGCGAACAACCGTGGGGACGTGAATTTGCGCTGCGCGATCCGGCCGGCAATTGCGTGCATTTTGTTGCCGAAGAGCAGGACTGA

>Sequence71

ATGGCGAAACTGACGAGCGCCGTACCGGTTCTGACTGCGCGCGATGTTGCGGGCGCCGTAGAATTTTGGACCGATCGTCTGGGCTTCTCGCGCGATTTCGTCGAAGATGATTTTGCTGGAGTTGTCCGCGACGACGTAACCTTGTTCATTTCAGCGGTTCAGGATCAGGTGGTGCCTGATAACACGCTGGCATGGGTGTGGGTTCGTGGCCTTGATGAACTCTATGCCGAATGGAGCGAAGTGGTGAGCACCAACTTTCGTGACGCCAGCGGCCCGGCCATGACCGAAATTGGTGAGCAGCCGTGGGGTCGCGAATTTGCGCTGCGCGATCCGGCTGGCAATTGCGTGCATTTTGTGGCGGAGGAACAAGATTAA

>Sequence72

ATGGCGAAACTCACATCAGCCGTGCCGGTACTTACTGCCCGTGACGTTGCCGGCGCGGTGGAGTTCTGGACCGATCGCCTGGGCTTTAGCCGTGATTTTGTAGAAGATGACTTTGCGGGCGTGGTCCGCGATGACGTGACCCTGTTTATCAGCGCCGTGCAAGATCAGGTCGTTCCGGATAATACGCTGGCGTGGGTTTGGGTGCGCGGTCTGGATGAATTGTATGCGGAATGGAGCGAAGTGGTAAGCACCAACTTTCGCGACGCAAGCGGTCCGGCGATGACGGAAATTGGTGAACAGCCGTGGGGCCGTGAATTCGCACTGCGCGATCCTGCGGGCAACTGCGTCCATTTTGTTGCGGAGGAACAGGATTAA

>Sequence73

ATGGCCAAACTGACGAGCGCCGTCCCGGTTCTGACCGCGCGTGATGTGGCCGGCGCAGTTGAATTCTGGACCGATCGCCTGGGATTTAGTCGCGATTTTGTGGAAGATGATTTTGCGGGTGTTGTGCGTGATGATGTGACGCTGTTTATCTCGGCCGTGCAGGATCAGGTTGTGCCTGATAATACTCTGGCCTGGGTATGGGTACGCGGTTTAGATGAGCTCTATGCGGAGTGGAGCGAAGTGGTGAGCACCAACTTCCGTGACGCTAGCGGCCCGGCCATGACAGAAATTGGCGAACAGCCGTGGGGGCGCGAATTTGCGCTTCGTGATCCGGCGGGCAACTGCGTTCATTTTGTGGCCGAGGAACAAGATTAA

>Sequence74

ATGGCCAAACTGACCTCGGCGGTGCCGGTTCTGACGGCGCGCGACGTCGCCGGCGCAGTAGAATTCTGGACCGACCGTTTGGGCTTTAGCCGTGATTTTGTTGAAGATGATTTTGCGGGCGTTGTTCGCGATGATGTTACCCTTTTCATTAGCGCGGTGCAGGACCAGGTTGTGCCGGACAATACCCTGGCCTGGGTGTGGGTCCGTGGACTGGATGAACTCTATGCCGAGTGGTCAGAGGTAGTTAGCACCAACTTTCGCGACGCCAGCGGTCCGGCAATGACCGAAATTGGTGAACAGCCGTGGGGCCGTGAATTTGCACTGCGCGATCCGGCGGGTAACTGCGTGCATTTCGTGGCGGAAGAACAAGATTGA

>Sequence75

ATGGCCAAACTGACTAGCGCGGTTCCGGTTCTGACAGCCCGTGACGTGGCGGGTGCGGTGGAGTTTTGGACCGATCGCCTGGGCTTTAGTCGTGACTTTGTCGAAGACGACTTTGCAGGTGTCGTTCGCGATGATGTGACCCTGTTTATCTCGGCGGTGCAAGATCAGGTTGTTCCGGATAACACGCTGGCCTGGGTTTGGGTTCGTGGGCTGGACGAACTGTATGCAGAATGGAGCGAGGTGGTTAGCACGAACTTTCGTGATGCTAGCGGTCCGGCCATGACCGAAATTGGCGAGCAGCCGTGGGGCCGCGAATTCGCGCTGCGCGATCCGGCTGGCAACTGCGTACATTTTGTGGCGGAAGAACAGGATTAA

>Sequence76

ATGGCAAAACTGACTAGCGCAGTTCCGGTTCTGACCGCTCGTGATGTTGCCGGCGCCGTTGAATTCTGGACCGATCGCCTTGGCTTCAGCCGCGATTTTGTCGAAGATGACTTTGCGGGCGTTGTGCGTGATGATGTTACGCTGTTTATTAGCGCTGTGCAGGACCAGGTTGTGCCGGATAACACGCTGGCGTGGGTCTGGGTGCGTGGTCTGGATGAACTCTATGCCGAATGGAGCGAAGTGGTATCGACCAACTTTCGTGATGCGAGTGGCCCTGCCATGACCGAAATTGGTGAACAGCCGTGGGGCCGCGAGTTTGCGTTGCGCGACCCGGCAGGCAACTGCGTACATTTTGTGGCGGAGGAACAAGATTGA

>Sequence77

ATGGCCAAACTGACTTCGGCAGTGCCGGTGCTGACGGCGCGTGATGTCGCAGGCGCTGTTGAATTTTGGACCGATCGCCTGGGTTTTAGCCGCGATTTTGTGGAAGATGACTTTGCGGGTGTGGTGCGCGATGATGTTACATTGTTTATCTCAGCCGTACAGGATCAGGTTGTTCCGGATAATACGCTGGCGTGGGTGTGGGTTCGTGGCCTTGATGAGCTCTATGCCGAATGGAGCGAAGTGGTTAGTACCAACTTCCGCGACGCAAGCGGGCCGGCCATGACCGAAATTGGCGAACAGCCGTGGGGCCGTGAATTTGCGTTACGTGATCCGGCGGGTAACTGCGTTCATTTTGTGGCGGAGGAACAAGACTAA

>Sequence78

ATGGCCAAACTCACCTCAGCAGTTCCGGTTCTGACCGCCCGCGATGTCGCCGGTGCGGTGGAATTCTGGACCGATCGTCTGGGCTTTAGCCGCGACTTTGTGGAAGACGACTTCGCCGGCGTCGTCCGTGATGATGTTACCCTGTTTATCAGCGCGGTGCAGGATCAGGTGGTGCCGGACAACACCCTGGCATGGGTTTGGGTGCGTGGCTTAGATGAGCTTTATGCCGAATGGAGCGAAGTTGTGAGTACCAACTTTCGCGATGCGTCGGGCCCGGCGATGACCGAAATCGGAGAACAGCCGTGGGGTCGCGAATTTGCCCTGCGTGACCCGGCTGGTAACTGCGTACATTTTGTTGCGGAAGAGCAAGATTGA

>Sequence79

ATGGCCAAACTGACGAGCGCGGTGCCGGTTTTGACCGCGCGCGATGTGGCGGGTGCAGTCGAGTTTTGGACCGACCGTCTGGGCTTTAGTCGCGATTTTGTTGAGGACGATTTCGCTGGTGTGGTTCGCGATGATGTTACCTTATTCATTAGCGCGGTACAGGATCAGGTTGTGCCGGACAACACCCTCGCGTGGGTATGGGTGCGTGGGCTGGATGAGCTGTATGCCGAATGGTCAGAAGTCGTGAGCACAAACTTTCGTGATGCATCGGGCCCGGCCATGACGGAAATTGGCGAACAGCCGTGGGGCCGCGAATTTGCGCTGCGTGACCCGGCCGGCAATTGCGTTCATTTCGTGGCCGAAGAACAGGACTAA

>Sequence80

ATGGCCAAACTGACCAGCGCTGTACCGGTGTTGACCGCACGCGACGTCGCGGGCGCGGTGGAATTTTGGACGGATCGCCTGGGTTTTAGCCGCGATTTCGTGGAGGATGATTTCGCTGGCGTAGTGCGTGACGATGTTACCCTGTTTATCAGCGCCGTTCAGGATCAAGTGGTACCGGACAATACCCTGGCGTGGGTGTGGGTTCGCGGCTTAGATGAACTGTATGCCGAGTGGTCGGAAGTGGTGTCAACGAACTTTCGTGATGCCAGCGGACCGGCGATGACCGAAATCGGCGAACAGCCGTGGGGTCGTGAATTCGCCCTGCGCGATCCTGCAGGTAACTGCGTTCATTTTGTCGCGGAGGAACAGGATTAA

>Sequence81

ATGGCGAAACTGACCAGCGCAGTGCCGGTTCTGACTGCTCGTGATGTAGCCGGCGCGGTGGAGTTTTGGACCGACCGTCTGGGCTTTTCGCGCGATTTTGTCGAAGATGATTTTGCGGGTGTTGTGCGCGACGACGTTACCCTGTTTATTAGCGCAGTCCAGGATCAGGTCGTGCCGGACAACACACTGGCCTGGGTGTGGGTTCGCGGGCTGGATGAATTATATGCGGAATGGAGCGAAGTTGTGAGCACCAACTTTCGTGATGCGAGCGGCCCGGCTATGACGGAAATCGGCGAACAGCCGTGGGGCCGCGAATTTGCCCTGCGCGATCCTGCCGGTAATTGCGTTCATTTCGTAGCCGAGGAGCAGGACTGA

>Sequence82

ATGGCGAAACTGACCAGTGCGGTTCCTGTCCTCACGGCGCGTGACGTTGCAGGTGCGGTCGAGTTTTGGACCGATCGTCTGGGCTTTAGCCGCGATTTTGTTGAAGATGATTTCGCGGGCGTCGTACGCGATGATGTGACCCTGTTTATCAGCGCCGTGCAGGATCAGGTGGTTCCGGACAACACATTGGCCTGGGTGTGGGTGCGCGGCCTGGACGAATTATATGCCGAATGGTCAGAAGTTGTGAGCACCAACTTTCGTGATGCATCGGGTCCGGCGATGACTGAAATCGGAGAACAGCCGTGGGGCCGCGAATTCGCGCTGCGCGACCCGGCCGGTAATTGCGTGCATTTTGTGGCCGAGGAACAAGATTAA

>Sequence83

ATGGCGAAACTGACCAGCGCCGTGCCGGTGCTGACTGCGCGCGACGTTGCCGGGGCCGTGGAATTCTGGACCGATCGTCTGGGTTTTAGCCGTGACTTTGTAGAAGATGACTTTGCGGGCGTCGTTCGCGATGATGTCACACTCTTTATCAGTGCCGTGCAAGACCAGGTTGTCCCGGATAATACGTTAGCGTGGGTTTGGGTGCGTGGCCTGGATGAACTGTATGCGGAATGGTCGGAGGTTGTGTCAACCAACTTTCGCGATGCAAGCGGCCCTGCGATGACCGAAATTGGCGAACAGCCGTGGGGTCGTGAATTTGCACTGCGCGATCCGGCCGGCAACTGCGTACATTTTGTTGCAGAAGAGCAGGATTGA

>Sequence84

ATGGCCAAACTGACCAGTGCGGTTCCTGTGCTGACCGCCCGTGACGTAGCGGGCGCCGTCGAATTTTGGACTGATCGCCTGGGCTTTAGCCGCGATTTTGTGGAGGATGATTTCGCGGGCGTGGTTCGTGATGATGTGACATTATTCATCTCAGCCGTTCAGGATCAGGTGGTACCGGATAACACCCTTGCCTGGGTGTGGGTCCGCGGACTCGATGAGCTGTATGCTGAGTGGAGCGAAGTAGTGAGCACGAATTTCCGCGATGCGAGCGGCCCGGCGATGACGGAAATCGGTGAACAGCCGTGGGGCCGCGAATTTGCACTGCGTGATCCGGCCGGTAACTGCGTGCATTTTGTTGCCGAAGAACAGGACTGA

>Sequence85

ATGGCGAAACTGACCAGTGCAGTCCCGGTGCTGACAGCGCGCGACGTGGCCGGCGCAGTTGAATTTTGGACCGATCGCCTGGGCTTCTCGCGTGATTTCGTTGAGGATGATTTTGCTGGTGTAGTTCGCGATGACGTGACCCTCTTTATTAGCGCAGTTCAGGATCAGGTGGTCCCGGATAACACGCTGGCGTGGGTTTGGGTTCGTGGCTTGGATGAACTGTATGCCGAATGGTCAGAGGTGGTAAGCACCAACTTTCGCGACGCGAGCGGCCCTGCCATGACCGAAATCGGTGAACAACCGTGGGGCCGTGAGTTTGCTCTGCGCGACCCGGCCGGCAACTGCGTGCATTTTGTGGCGGAAGAACAGGATTGA

>Sequence86

ATGGCAAAACTTACCAGTGCAGTTCCTGTTCTCACGGCCCGTGATGTGGCCGGTGCCGTGGAATTTTGGACCGATCGCCTGGGATTTTCACGCGATTTCGTCGAAGATGATTTTGCGGGCGTCGTGCGCGACGATGTAACCCTGTTTATTAGCGCGGTTCAGGATCAGGTGGTTCCGGACAACACGCTGGCGTGGGTATGGGTGCGCGGGTTGGATGAGCTGTATGCAGAATGGAGCGAAGTGGTGTCGACAAACTTTCGTGACGCCAGCGGCCCGGCGATGACCGAAATCGGTGAACAGCCGTGGGGCCGCGAATTTGCGTTACGTGATCCGGCCGGCAATTGCGTGCATTTTGTCGCCGAGGAGCAGGATTAA

>Sequence87

ATGGCTAAATTAACGTCGGCCGTTCCGGTACTGACCGCGCGCGACGTAGCGGGTGCCGTGGAGTTTTGGACCGATCGCCTGGGTTTTTCACGCGACTTCGTTGAGGATGATTTTGCAGGCGTTGTCCGTGACGACGTCACCCTCTTCATCAGCGCGGTACAAGATCAGGTGGTGCCGGATAACACTCTGGCCTGGGTTTGGGTGCGTGGCCTGGATGAACTGTATGCAGAATGGAGCGAAGTGGTTAGCACGAACTTCCGCGATGCTAGCGGCCCGGCAATGACCGAAATTGGCGAACAGCCGTGGGGCCGCGAGTTTGCGCTGCGTGATCCGGCGGGTAATTGCGTGCATTTTGTGGCCGAAGAACAGGATTGA

>Sequence88

ATGGCCAAACTGACCTCGGCCGTGCCGGTATTAACCGCCCGTGACGTTGCTGGCGCGGTAGAATTTTGGACGGATCGTCTCGGTTTTAGCCGTGACTTTGTCGAAGATGACTTTGCCGGCGTGGTTCGCGATGACGTTACCCTGTTTATCAGCGCGGTGCAGGATCAAGTTGTGCCGGATAACACCCTGGCATGGGTGTGGGTCCGCGGATTGGATGAACTGTATGCGGAATGGTCAGAAGTTGTGAGCACGAATTTCCGCGATGCGAGTGGCCCGGCCATGACTGAAATTGGTGAACAGCCGTGGGGTCGCGAATTTGCACTGCGTGATCCTGCGGGCAACTGCGTGCATTTCGTCGCGGAAGAGCAGGACTAA

>Sequence89

ATGGCCAAACTGACTAGCGCCGTGCCGGTTTTAACGGCGCGTGACGTGGCCGGTGCGGTTGAATTTTGGACCGACCGTCTGGGTTTTAGCCGCGATTTTGTGGAAGATGATTTCGCCGGCGTGGTACGCGACGATGTAACCTTGTTCATCAGCGCGGTCCAGGATCAGGTTGTTCCTGATAATACACTGGCATGGGTCTGGGTGCGCGGTCTGGACGAACTGTATGCCGAATGGAGCGAAGTCGTTAGTACGAACTTTCGTGATGCCAGCGGCCCGGCGATGACCGAAATTGGCGAGCAGCCGTGGGGCCGCGAATTCGCCCTTCGTGATCCGGCGGGTAACTGCGTACATTTTGTGGCTGAAGAACAGGACTGA

>Sequence90

ATGGCAAAATTGACCTCAGCGGTGCCGGTTCTGACCGCGCGTGACGTAGCCGGCGCGGTGGAATTTTGGACCGACCGCCTGGGTTTTAGCCGCGATTTTGTTGAAGATGACTTCGCGGGCGTGGTGCGTGATGACGTTACCCTGTTTATCAGCGCCGTTCAGGATCAGGTTGTGCCGGATAACACGCTCGCATGGGTGTGGGTTCGTGGGCTTGATGAACTGTATGCCGAATGGAGCGAAGTGGTGTCGACTAACTTCCGCGATGCCAGTGGCCCGGCGATGACAGAAATTGGCGAGCAACCGTGGGGTCGCGAGTTCGCACTGCGTGATCCGGCTGGTAATTGCGTCCATTTTGTGGCCGAGGAACAGGATTAA

>Sequence91

ATGGCGAAACTGACCAGCGCGGTGCCGGTGCTGACCGCGCGTGACGTCGCGGGAGCGGTGGAATTTTGGACCGATCGCCTGGGCTTCTCACGTGACTTTGTAGAAGACGATTTTGCGGGCGTAGTGCGCGATGACGTTACGCTGTTTATTAGCGCCGTGCAGGATCAGGTTGTGCCGGACAATACGCTCGCGTGGGTCTGGGTGCGCGGTTTGGATGAACTGTATGCAGAATGGAGTGAAGTTGTTAGCACCAACTTTCGTGATGCCTCGGGTCCGGCAATGACAGAAATCGGCGAACAACCTTGGGGCCGCGAATTTGCCCTTCGTGATCCGGCGGGGAACTGCGTCCATTTTGTAGCCGAGGAACAGGATTGA

>Sequence92

ATGGCAAAACTGACCAGCGCAGTACCGGTACTGACGGCGCGTGATGTGGCGGGTGCCGTGGAGTTCTGGACCGACCGCCTGGGCTTTAGCCGTGACTTCGTGGAAGACGATTTTGCGGGTGTGGTTCGCGACGATGTTACTTTATTTATTTCGGCAGTGCAGGATCAGGTTGTGCCGGATAACACCCTCGCTTGGGTTTGGGTTCGCGGCCTGGATGAACTGTATGCGGAGTGGAGCGAAGTGGTGAGCACGAACTTTCGTGATGCCTCAGGCCCGGCCATGACAGAAATCGGCGAACAGCCGTGGGGCCGCGAATTTGCTCTTCGCGACCCGGCCGGCAATTGCGTGCATTTCGTGGCGGAAGAGCAAGATTGA

>Sequence93

ATGGCGAAACTGACTAGTGCGGTGCCGGTCCTGACAGCGCGCGACGTGGCGGGTGCCGTCGAATTTTGGACGGACCGCCTGGGCTTTAGCCGCGATTTTGTGGAAGATGATTTTGCCGGTGTGGTTCGTGACGATGTTACCCTGTTTATCTCGGCCGTACAGGATCAGGTGGTTCCGGACAATACCCTGGCGTGGGTATGGGTGCGTGGATTAGATGAGTTGTATGCGGAATGGAGCGAAGTGGTATCAACGAACTTTCGCGATGCGAGCGGCCCGGCCATGACCGAAATTGGCGAGCAGCCTTGGGGTCGTGAATTCGCGCTGCGTGATCCGGCGGGCAACTGCGTCCATTTCGTTGCCGAAGAACAAGACTGA

>Sequence94

ATGGCGAAACTTACAAGCGCAGTGCCGGTACTGACGGCCCGTGATGTTGCGGGTGCAGTGGAATTCTGGACGGATCGCCTGGGTTTTTCACGCGATTTTGTGGAGGACGATTTTGCGGGCGTTGTTCGCGACGACGTGACTCTGTTTATTAGCGCCGTTCAGGATCAGGTGGTCCCGGATAACACCTTAGCGTGGGTGTGGGTACGCGGCCTGGATGAACTGTATGCCGAGTGGTCGGAAGTCGTAAGCACCAACTTTCGTGATGCCAGTGGCCCGGCAATGACCGAAATCGGCGAACAGCCGTGGGGTCGTGAATTTGCGCTCCGCGATCCGGCCGGCAATTGCGTTCATTTTGTGGCCGAAGAGCAAGACTGA

>Sequence95

ATGGCCAAACTTACGTCGGCGGTGCCGGTCCTGACTGCCCGCGACGTGGCGGGCGCAGTGGAGTTTTGGACGGATCGCTTAGGTTTCAGCCGTGACTTTGTGGAGGATGATTTTGCGGGCGTTGTGCGCGACGATGTAACCCTGTTTATCTCAGCCGTTCAAGATCAGGTTGTTCCGGATAACACCCTGGCGTGGGTCTGGGTCCGCGGTCTCGATGAATTGTATGCCGAATGGAGCGAAGTTGTTAGCACCAATTTCCGCGACGCTAGCGGCCCGGCGATGACCGAAATCGGCGAACAGCCGTGGGGTCGTGAATTTGCGCTGCGTGATCCTGCGGGCAACTGCGTGCATTTCGTGGCGGAAGAACAGGATTGA

>Sequence96

ATGGCCAAACTGACCAGCGCGGTTCCTGTTCTGACTGCCCGTGATGTGGCGGGCGCCGTTGAATTCTGGACGGACCGCCTTGGCTTTTCGCGCGACTTCGTCGAAGATGATTTTGCAGGCGTGGTGCGTGACGATGTGACCCTGTTTATCAGTGCGGTACAGGATCAGGTAGTTCCGGACAACACACTGGCATGGGTCTGGGTACGCGGCCTGGACGAACTCTATGCCGAATGGTCAGAAGTTGTGAGCACCAACTTTCGCGATGCTAGCGGCCCGGCGATGACCGAGATTGGAGAACAGCCGTGGGGTCGTGAGTTTGCCCTGCGTGATCCGGCAGGTAACTGCGTGCATTTTGTCGCGGAAGAACAGGATTGA

>Sequence97

ATGGCAAAACTGACCAGCGCGGTGCCGGTTCTGACCGCACGTGACGTTGCGGGCGCGGTAGAGTTTTGGACCGACCGCCTGGGTTTTAGCCGCGATTTTGTCGAAGACGATTTTGCAGGGGTGGTGCGTGATGACGTGACGCTTTTTATTAGCGCGGTTCAGGATCAGGTTGTGCCGGATAACACATTGGCGTGGGTGTGGGTACGCGGTTTAGACGAGCTGTATGCTGAATGGAGCGAAGTGGTGAGCACGAACTTCCGTGATGCGTCGGGTCCGGCCATGACCGAAATTGGCGAACAACCGTGGGGCCGTGAATTTGCCCTCCGCGATCCTGCCGGCAATTGCGTTCATTTCGTTGCCGAAGAACAGGATTGA

>Sequence98

ATGGCCAAATTAACCAGCGCGGTTCCGGTTTTGACCGCGCGCGATGTCGCGGGTGCCGTGGAATTTTGGACCGACCGCCTGGGTTTTAGCCGTGACTTTGTTGAGGATGATTTCGCCGGAGTGGTGCGTGATGACGTTACGCTGTTCATCAGCGCAGTGCAGGATCAGGTGGTCCCGGACAACACACTGGCGTGGGTGTGGGTGCGCGGCCTGGATGAACTCTATGCGGAATGGTCGGAAGTAGTGTCAACTAACTTTCGCGATGCGAGCGGCCCGGCCATGACGGAGATTGGCGAACAGCCGTGGGGCCGTGAGTTTGCCCTGCGCGATCCTGCCGGCAATTGCGTGCATTTTGTAGCCGAAGAACAAGACTGA

>Sequence99

ATGGCTAAACTCACCAGCGCGGTGCCGGTGCTGACCGCGCGCGATGTGGCTGGTGCGGTTGAATTTTGGACTGACCGTTTAGGCTTCTCACGTGATTTTGTGGAAGATGATTTCGCGGGCGTGGTTCGTGACGATGTAACACTGTTTATCAGCGCGGTCCAGGATCAAGTAGTGCCGGACAACACCCTGGCCTGGGTGTGGGTGCGCGGACTGGATGAACTTTATGCCGAGTGGAGCGAGGTAGTTTCGACCAACTTTCGCGATGCCAGCGGCCCGGCCATGACGGAAATTGGGGAGCAGCCGTGGGGCCGCGAATTCGCCTTGCGTGACCCGGCGGGTAACTGCGTCCATTTTGTTGCGGAAGAACAGGATTAA

>Sequence100

ATGGCCAAACTGACCAGTGCAGTACCGGTTCTGACCGCGCGCGATGTGGCGGGGGCCGTGGAATTTTGGACGGATCGCTTGGGTTTCAGCCGTGACTTTGTGGAAGATGACTTTGCCGGCGTGGTCCGTGATGACGTCACCCTGTTTATCAGCGCAGTTCAGGATCAAGTCGTTCCGGATAATACCTTAGCGTGGGTGTGGGTGCGCGGCCTGGATGAGCTTTATGCGGAATGGTCAGAAGTTGTTAGCACCAACTTCCGCGATGCGAGCGGCCCTGCCATGACTGAAATTGGTGAACAGCCGTGGGGCCGTGAATTCGCCCTCCGTGACCCGGCTGGAAACTGCGTGCATTTTGTGGCAGAGGAGCAGGACTGA

>Sequence101

ATGGCGAAACTCACGAGTGCGGTCCCTGTACTGACAGCCCGGGATGTAGCGGGCGCAGTTGAATTCTGGACTGACCGTCTTGGCTTTTCTCGAGATTTTGTCGAGGACGACTTCGCTGGCGTTGTTCGCGATGATGTTACCTTGTTCATCTCGGCGGTACAAGATCAGGTGGTACCCGACAATACGCTAGCCTGGGTGTGGGTACGTGGGCTGGATGAACTGTATGCAGAGTGGTCAGAGGTGGTATCCACGAATTTTAGGGATGCCAGCGGTCCGGCCATGACCGAAATTGGCGAACAACCGTGGGGACGCGAGTTCGCATTAAGGGATCCAGCGGGTAATTGCGTCCATTTCGTGGCCGAAGAACAAGATTAA

>Sequence102

ATGGCTAAACTGACTTCGGCGGTCCCGGTGCTGACGGCGCGCGATGTAGCTGGGGCAGTCGAATTCTGGACAGATCGACTCGGATTCAGTCGCGACTTTGTAGAAGACGACTTCGCCGGCGTGGTACGTGACGACGTTACCCTGTTCATTTCAGCTGTGCAAGACCAGGTTGTCCCAGACAATACGCTAGCATGGGTGTGGGTCCGTGGCTTGGACGAACTTTATGCAGAATGGTCTGAAGTGGTGAGCACCAATTTCCGCGATGCCTCCGGGCCTGCAATGACCGAGATTGGCGAACAGCCTTGGGGACGGGAATTCGCCTTACGTGATCCCGCCGGCAACTGCGTTCATTTCGTAGCGGAGGAGCAAGATTGA

>Sequence103

ATGGCGAAACTTACTTCCGCGGTTCCGGTCCTAACCGCTCGCGATGTCGCGGGTGCTGTGGAGTTCTGGACCGACCGTCTCGGCTTCTCGCGTGATTTCGTGGAGGATGACTTCGCCGGAGTGGTCCGTGATGACGTAACGCTGTTCATTTCTGCAGTACAAGACCAGGTCGTGCCCGACAACACATTGGCCTGGGTTTGGGTACGGGGCCTCGATGAGTTATATGCCGAATGGAGTGAAGTCGTAAGCACCAATTTTCGCGATGCCTCAGGACCAGCAATGACGGAAATTGGCGAGCAACCGTGGGGGCGAGAATTCGCTCTGCGCGACCCTGCAGGCAATTGCGTACATTTTGTCGCCGAAGAGCAGGATTAA

>Sequence104

ATGGCCAAACTAACATCGGCAGTCCCTGTATTAACTGCGCGTGATGTAGCGGGAGCAGTGGAATTTTGGACCGACCGTCTCGGCTTTTCTCGGGATTTCGTGGAGGATGATTTCGCAGGAGTTGTCCGCGATGACGTTACGCTTTTCATTTCGGCCGTCCAAGATCAGGTTGTACCGGATAACACACTAGCTTGGGTATGGGTCCGCGGGTTGGACGAGCTGTATGCGGAATGGTCAGAGGTTGTGTCCACAAATTTCCGCGACGCGAGTGGCCCAGCGATGACGGAAATTGGGGAACAGCCCTGGGGCCGAGAATTCGCATTACGTGATCCGGCTGGCAATTGCGTGCATTTCGTGGCCGAGGAACAAGACTAA

>Sequence105

ATGGCAAAACTGACCTCGGCAGTGCCTGTCTTAACTGCGAGAGACGTTGCAGGTGCCGTAGAATTCTGGACGGATCGTTTGGGCTTCTCTCGCGACTTTGTCGAAGATGACTTCGCGGGGGTAGTACGTGACGATGTCACCCTCTTTATCTCAGCAGTGCAGGATCAAGTGGTTCCAGACAACACCCTTGCTTGGGTGTGGGTTCGGGGACTAGATGAGCTGTATGCAGAATGGTCAGAGGTCGTTAGTACAAATTTTCGCGATGCCTCCGGCCCGGCGATGACGGAGATTGGCGAGCAGCCATGGGGTCGAGAGTTCGCGCTACGCGACCCCGCAGGCAATTGCGTTCATTTTGTGGCGGAAGAACAAGATTAA

>Sequence106

ATGGCTAAATTAACCAGTGCAGTTCCTGTGCTGACCGCACGCGATGTGGCTGGTGCCGTAGAATTCTGGACGGATCGACTCGGGTTTAGTCGGGACTTCGTGGAGGATGATTTTGCAGGAGTAGTCCGAGACGATGTCACACTATTTATTTCGGCCGTTCAGGATCAAGTAGTGCCAGATAACACGTTGGCCTGGGTGTGGGTTCGTGGCCTTGATGAGCTTTATGCAGAATGGTCCGAAGTCGTTTCTACCAATTTTCGGGATGCCTCAGGACCCGCGATGACTGAAATTGGCGAACAACCTTGGGGTCGCGAATTCGCCCTGCGTGACCCGGCTGGCAATTGCGTGCATTTCGTCGCAGAGGAACAAGACTGA

>Sequence107

ATGGCGAAATTAACCAGCGCCGTCCCCGTTCTAACTGCACGTGATGTGGCAGGCGCGGTTGAGTTCTGGACAGATCGTTTGGGATTTTCTCGAGATTTCGTAGAAGATGACTTTGCCGGCGTCGTACGTGATGATGTAACGCTGTTCATCTCAGCCGTCCAAGACCAAGTAGTGCCAGACAACACGCTCGCATGGGTGTGGGTCAGGGGGCTCGATGAGCTTTATGCGGAATGGTCCGAAGTGGTGTCGACCAATTTCCGGGATGCAAGTGGGCCGGCCATGACTGAGATTGGCGAACAGCCTTGGGGTCGCGAATTCGCGCTCCGCGACCCGGCGGGAAATTGCGTTCATTTCGTCGCTGAAGAGCAGGATTGA

>Sequence108

ATGGCAAAACTTACCTCTGCTGTTCCTGTACTGACAGCCAGGGACGTCGCGGGTGCCGTAGAGTTCTGGACGGATCGTCTGGGGTTTTCCCGGGATTTCGTTGAAGACGATTTTGCCGGCGTTGTACGTGACGACGTGACTCTCTTCATTAGTGCGGTGCAAGACCAAGTGGTACCGGATAATACGTTGGCGTGGGTCTGGGTCCGTGGTTTAGATGAATTATATGCCGAATGGTCGGAAGTGGTCTCGACAAATTTCCGAGATGCGTCAGGACCCGCCATGACCGAGATTGGCGAACAGCCGTGGGGACGCGAGTTCGCACTAAGGGATCCAGCGGGAAACTGCGTGCATTTTGTAGCAGAAGAACAAGATTGA

>Sequence109

ATGGCTAAATTGACGAGTGCGGTACCAGTTTTAACGGCACGTGACGTTGCAGGGGCTGTAGAATTCTGGACCGATCGGCTCGGCTTTTCACGGGATTTTGTAGAGGACGATTTTGCAGGGGTGGTCAGGGATGACGTTACCCTTTTCATCAGCGCGGTACAAGATCAAGTCGTGCCGGACAATACGTTAGCCTGGGTGTGGGTTCGCGGTCTAGATGAGCTGTATGCAGAATGGTCCGAGGTCGTATCTACTAACTTTCGCGACGCTTCGGGTCCCGCGATGACAGAAATCGGCGAACAACCGTGGGGCCGAGAATTCGCCTTACGTGATCCTGCGGGAAATTGCGTTCATTTCGTAGCGGAAGAACAGGACTAA

>Sequence110

ATGGCAAAACTCACATCCGCGGTTCCAGTTCTAACCGCCCGTGACGTCGCCGGCGCAGTAGAATTCTGGACCGACCGTTTAGGATTCTCACGCGACTTTGTAGAAGATGACTTTGCGGGAGTAGTCCGGGATGATGTTACACTCTTCATCTCGGCTGTACAGGATCAAGTAGTGCCCGACAATACTCTTGCATGGGTATGGGTTCGAGGGTTGGATGAATTGTATGCAGAATGGTCGGAAGTCGTCAGTACGAACTTTAGAGACGCATCTGGTCCTGCGATGACGGAAATTGGCGAGCAACCGTGGGGTCGCGAGTTTGCACTTCGGGACCCGGCTGGCAATTGCGTTCATTTCGTGGCGGAAGAGCAAGATTAA

>Sequence111

ATGGCGAAACTGACCAGCGCGGTGCCGGTGCTGACCGCGCGCGATGTGGCGGGCGCGGTGGAATTTTGGACCGATCGCCTGGGCTTTAGCCGCGATTTTGTGGAAGATGATTTTGCGGGCGTGGTGCGCGATGATGTGACCCTGTTTATTAGCGCGGTGCAGGATCAGGTGGTGCCGGATAACACCCTGGCGTGGGTGTGGGTGCGCGGCCTGGATGAACTGTATGCGGAATGGAGCGAAGTGGTGAGCACCAACTTTCGCGATGCGAGCGGCCCGGCGATGACCGAAATTGGCGAACAGCCGTGGGGCCGCGAATTTGCGCTGCGCGATCCGGCGGGCAACTGCGTGCATTTTGTGGCGGAAGAACAGGATTAA

>Sequence112

ATGGCCAAACTGACCAGCGCCGTTCCGGTGCTGACCGCGCGCGATGTGGCCGGCGCGGTTGAATTTTGGACCGATCGCCTGGGCTTTAGCCGCGATTTTGTGGAAGATGATTTTGCCGGTGTGGTTCGTGATGATGTGACCCTGTTTATCAGCGCGGTGCAGGATCAGGTGGTGCCGGATAACACCCTGGCCTGGGTGTGGGTGCGCGGCCTGGATGAACTGTATGCCGAATGGAGCGAAGTGGTGAGCACCAACTTTCGTGATGCCAGCGGCCCGGCCATGACCGAAATCGGCGAACAGCCGTGGGGCCGCGAATTTGCCCTGCGCGATCCGGCCGGCAACTGCGTGCATTTTGTGGCCGAAGAACAGGATTGA
