## Supplemental Figure 2 for "A selectable system to evaluate synthetic gene optimization features"

```
Supplemental Figure 2.
#
#
#  Percent Identity  Matrix - created by Clustal2.1
#
#

     1: Sequence103  100.00   78.13   78.67   76.53   78.13   77.87   76.53   77.87   76.53   77.87   79.73   79.20   78.13   79.73   78.13   76.00   77.60   75.47   76.53   76.80   78.67   77.60   80.00   78.93   77.60   78.13   76.53   80.27   80.00   78.13   80.00   76.27   75.73   78.67   78.13   80.00   79.47   77.87   78.13   78.93   76.27   78.93   77.07   77.87   77.87   78.67   75.73   79.47   80.80   77.33   77.87   78.13   77.33   79.47   78.40   77.33   76.53   76.53   75.73   76.53   79.20   77.07   76.27   79.47   79.20   76.53   79.20   78.13   76.80   78.40   77.87   75.73   80.00   78.40   78.93   75.73   78.13   80.27   78.93   78.13   77.60   77.60   78.40   80.53   76.80   78.40   78.13   78.13   77.60   78.40   78.67   75.73   78.93   79.20   80.00   77.87   77.07   80.27   81.07   78.40   78.40   79.47   80.00   77.33   78.93   77.60   77.60   76.00   81.60   76.53   78.67   78.40
     2: Sequence105   78.13  100.00   81.33   77.33   78.93   78.13   77.07   78.93   76.80   77.33   77.60   77.60   76.53   76.80   79.20   78.40   76.53   77.33   76.27   78.13   78.93   77.60   81.33   79.73   79.20   76.80   78.67   78.67   78.67   77.60   79.47   76.27   78.40   78.13   75.20   76.27   76.80   76.53   77.60   76.80   77.33   77.07   76.53   76.27   75.73   76.00   78.40   78.67   77.60   77.87   75.20   76.80   76.53   76.53   77.07   78.93   77.07   77.33   77.87   77.87   76.53   77.87   77.60   76.00   78.93   77.60   78.40   78.40   78.67   78.40   80.53   79.20   79.20   80.00   78.40   79.20   79.20   78.67   78.93   79.47   79.20   76.00   78.13   77.07   76.53   78.67   79.47   79.20   77.87   77.60   76.27   76.53   77.07   77.07   78.13   78.40   77.60   80.53   78.13   76.00   78.13   77.07   77.33   76.53   77.33   77.07   78.40   78.93   77.60   76.27   78.40   77.07
     3: Sequence110   78.67   81.33  100.00   78.40   77.33   77.33   75.47   76.80   78.93   76.80   78.13   76.80   74.93   77.33   77.07   74.40   77.60   76.80   77.33   75.47   76.53   76.80   78.13   79.20   77.33   76.27   79.20   76.27   77.07   73.87   78.13   78.67   77.07   77.87   76.27   76.80   75.20   75.73   76.53   76.27   74.93   75.47   75.20   78.13   78.40   77.33   75.20   76.53   76.27   76.27   76.27   76.27   78.13   77.33   76.00   76.00   73.60   75.47   76.00   74.67   77.87   78.93   77.33   74.93   78.13   76.53   78.67   76.27   76.80   77.60   77.60   78.67   79.47   76.53   76.00   79.47   78.13   76.53   77.33   80.00   78.93   78.13   79.47   75.73   77.07   77.87   77.87   78.13   77.07   75.73   76.53   74.40   75.47   76.53   77.33   74.93   76.80   78.13   77.87   75.73   77.07   76.27   76.27   77.07   74.67   76.27   76.27   76.53   76.80   75.73   76.27   75.47
     4: Sequence109   76.53   77.33   78.40  100.00   79.47   76.27   76.27   76.27   79.20   78.67   76.53   78.13   76.80   77.87   78.13   76.80   75.20   78.40   80.00   78.40   77.60   77.07   77.87   75.20   76.53   77.87   78.13   76.53   76.00   77.33   75.73   76.00   77.33   75.20   80.80   78.13   76.00   77.07   77.60   75.73   76.00   77.87   79.47   76.80   80.00   76.27   78.67   76.53   76.00   76.27   79.20   76.53   76.80   78.67   77.07   76.53   77.87   78.13   77.87   77.33   78.93   77.87   76.80   76.27   73.33   77.87   79.73   74.67   79.20   73.87   74.93   77.87   76.53   78.13   78.93   76.00   77.33   77.33   76.00   77.87   80.00   78.13   80.00   77.07   78.93   77.33   79.20   78.40   76.00   77.87   74.13   78.40   78.40   75.73   79.20   76.53   74.40   78.67   78.67   76.27   78.93   77.60   78.40   76.80   74.67   77.87   75.47   77.87   75.73   76.53   77.33   76.80
     5: Sequence106   78.13   78.93   77.33   79.47  100.00   80.00   77.87   78.40   76.00   78.13   78.13   78.67   78.93   76.53   77.87   79.47   77.87   78.67   78.67   78.93   78.67   77.60   79.47   78.40   78.40   75.73   75.73   78.13   79.73   78.40   79.73   77.87   77.87   80.27   77.07   76.27   78.13   78.13   78.40   78.13   78.93   78.40   80.00   78.93   77.07   78.13   79.73   77.33   77.33   78.40   80.00   80.27   77.87   79.47   78.67   79.47   79.73   78.67   78.93   80.80   73.60   77.87   78.40   77.60   77.87   75.73   78.13   77.07   75.73   79.47   77.87   77.07   79.47   78.40   78.67   78.67   78.93   78.93   79.73   80.27   78.93   76.80   77.87   79.73   75.20   77.60   79.73   79.73   78.93   79.20   78.13   77.87   78.67   76.53   80.00   78.13   76.00   77.07   78.13   77.87   77.87   78.93   78.13   77.33   77.33   76.27   79.73   77.60   78.13   78.93   78.67   79.20
     6: Sequence107   77.87   78.13   77.33   76.27   80.00  100.00   77.07   78.93   78.40   77.07   78.13   76.80   76.27   77.87   77.33   77.87   78.40   78.67   77.07   80.53   78.93   74.93   77.60   79.73   78.67   78.93   76.00   76.80   80.00   78.13   78.93   80.00   77.33   76.80   76.80   78.93   79.47   78.67   76.80   79.47   76.80   77.07   77.33   80.53   75.73   78.13   77.07   81.07   76.27   77.60   77.87   78.93   76.80   78.13   80.00   77.33   77.07   78.93   77.33   78.13   77.07   78.67   79.20   80.00   79.20   77.07   77.07   76.80   77.07   77.07   77.33   78.93   77.07   77.60   77.33   77.07   78.67   77.33   74.93   79.47   74.93   75.73   77.07   77.33   76.53   77.87   77.87   79.20   77.87   77.60   77.07   77.07   77.60   78.40   79.73   78.67   77.60   77.07   79.20   79.47   77.07   76.27   77.07   77.87   77.87   76.27   77.87   77.87   78.40   75.47   78.40   78.67
     7: Sequence108   76.53   77.07   75.47   76.27   77.87   77.07  100.00   77.87   78.93   75.73   77.60   80.27   77.60   77.33   79.20   78.40   78.40   77.87   77.87   78.40   74.40   77.87   74.13   77.07   78.13   79.73   78.40   74.93   77.60   79.47   76.27   74.93   74.67   76.00   78.13   74.67   77.60   79.47   78.13   75.73   76.00   77.87   79.20   77.60   76.80   76.53   78.40   76.80   79.47   76.27   76.27   77.87   76.53   76.53   77.07   77.87   77.33   76.00   78.67   76.27   77.33   76.27   78.40   77.60   74.40   75.47   74.67   76.80   73.33   77.60   76.53   77.87   77.87   77.60   75.73   79.47   77.33   77.07   76.00   76.80   77.07   77.87   77.60   78.13   76.27   76.00   76.00   75.73   74.67   75.47   78.13   78.13   76.00   78.13   76.80   77.07   76.53   76.53   77.87   79.47   78.93   76.80   77.33   76.53   78.67   76.27   76.53   75.47   76.53   75.73   77.07   76.80
     8: Sequence102   77.87   78.93   76.80   76.27   78.40   78.93   77.87  100.00   78.67   78.13   77.33   77.60   77.87   80.53   75.73   76.27   78.40   75.73   77.07   76.53   76.00   76.80   78.93   78.67   78.13   78.67   80.27   77.07   79.20   79.47   78.13   78.67   80.80   77.87   77.07   76.53   77.33   77.87   77.60   79.73   76.53   77.87   78.40   76.00   80.00   78.67   77.33   79.47   80.00   80.00   76.53   80.00   78.40   76.80   77.87   78.40   75.47   78.40   78.13   79.47   77.07   77.60   79.73   77.07   78.13   77.33   77.87   78.40   78.13   80.80   81.07   79.73   80.27   77.07   76.80   78.67   78.67   79.47   78.13   80.00   78.67   78.13   77.87   78.67   78.93   78.13   78.40   79.47   77.87   77.33   79.73   77.60   76.27   77.60   78.67   78.13   76.53   76.80   77.33   77.87   79.20   80.00   76.53   76.80   76.80   76.00   79.47   81.07   78.40   79.20   78.67   78.67
     9: Sequence101   76.53   76.80   78.93   79.20   76.00   78.40   78.93   78.67  100.00   80.53   77.33   78.93   77.07   77.33   77.33   78.93   77.87   78.13   78.67   77.60   79.73   78.93   77.87   79.20   79.20   79.47   80.80   78.13   79.47   77.07   79.73   78.67   77.87   77.33   75.47   77.60   77.87   79.47   79.73   79.47   79.73   77.33   78.13   78.67   76.80   76.80   78.40   78.93   75.73   76.53   78.67   75.73   77.33   78.67   76.27   80.00   78.13   77.87   77.33   74.13   77.60   78.93   77.33   80.27   77.33   77.33   80.53   76.00   78.13   76.80   77.87   77.07   75.47   77.60   79.47   79.20   76.80   75.20   74.93   79.73   77.33   76.80   79.20   76.80   78.67   81.60   76.00   79.47   79.20   80.00   78.40   78.93   78.93   77.33   76.00   78.93   77.60   76.53   78.13   78.93   78.93   76.00   78.67   78.67   77.33   78.93   79.20   78.13   77.60   76.53   78.67   78.40
    10: Sequence104   77.87   77.33   76.80   78.67   78.13   77.07   75.73   78.13   80.53  100.00   77.60   78.93   80.27   78.67   78.67   77.87   76.00   80.00   77.33   77.07   76.00   78.13   76.27   78.13   77.33   79.47   78.67   76.53   76.53   78.67   78.67   79.47   78.67   77.60   76.53   79.20   78.93   78.67   78.40   76.27   79.73   79.73   74.67   75.47   76.27   77.60   76.00   77.87   77.60   77.07   78.13   77.87   76.27   75.20   77.87   77.60   77.60   79.47   77.60   77.60   77.87   78.93   76.53   76.27   76.80   77.33   80.53   78.40   78.93   78.13   76.80   80.80   75.47   77.33   76.27   75.73   77.33   74.67   74.93   77.33   75.73   74.67   77.60   77.33   77.87   77.33   76.53   77.33   77.60   77.60   76.27   77.07   77.87   78.93   77.07   76.27   76.80   77.33   74.67   77.07   78.13   76.27   79.20   78.13   78.40   76.27   78.13   79.73   76.27   78.13   78.13   76.00
    11: Sequence17    79.73   77.60   78.13   76.53   78.13   78.13   77.60   77.33   77.33   77.60  100.00   85.33   83.73   84.27   85.07   84.00   82.67   84.00   83.47   82.93   83.73   84.00   82.40   80.80   82.93   83.47   85.60   83.73   83.73   83.47   84.53   81.60   81.60   82.40   81.33   84.80   84.80   83.47   83.47   83.73   82.40   82.93   84.27   83.20   85.07   83.47   79.47   85.33   84.00   84.53   82.67   83.73   81.33   82.13   82.40   80.53   84.00   82.40   81.33   84.27   84.00   82.67   82.40   82.13   81.33   82.93   82.13   83.20   85.60   84.80   83.47   82.67   83.20   81.60   84.27   84.00   80.53   81.33   81.60   82.13   83.20   84.27   84.53   83.73   83.20   82.40   82.67   82.67   83.47   84.00   84.53   83.47   85.60   85.87   82.40   82.13   83.73   82.40   84.80   84.53   84.00   84.27   84.00   84.00   85.07   85.33   84.53   85.07   85.87   84.27   88.00   87.47
    12: Sequence87    79.20   77.60   76.80   78.13   78.67   76.80   80.27   77.60   78.93   78.93   85.33  100.00   86.13   85.07   83.73   83.20   81.07   83.20   82.93   82.40   81.33   83.47   82.40   81.60   83.47   82.40   82.40   82.40   82.93   83.47   84.27   81.87   81.60   81.33   81.60   82.40   81.07   81.07   82.67   83.73   82.93   84.00   82.93   83.73   82.13   82.13   80.53   83.20   82.93   84.80   81.87   82.13   82.93   80.80   80.27   81.60   83.73   79.73   83.47   81.60   82.67   84.53   81.33   83.73   82.93   82.93   82.13   84.53   82.40   82.93   84.00   81.60   82.67   80.53   82.67   83.20   82.67   81.07   80.80   81.87   83.47   81.87   84.27   82.13   81.33   84.53   82.13   79.73   82.67   82.93   83.47   83.47   83.20   84.53   83.47   81.33   82.40   83.73   84.80   85.07   82.40   83.47   83.73   83.47   83.73   83.73   84.27   81.07   82.13   83.47   86.40   86.40
    13: Sequence26    78.13   76.53   74.93   76.80   78.93   76.27   77.60   77.87   77.07   80.27   83.73   86.13  100.00   87.47   86.40   84.53   83.47   84.27   83.20   82.40   81.07   85.07   83.73   81.33   82.67   83.20   82.67   84.27   82.67   83.20   84.27   83.20   85.07   83.20   82.93   83.20   82.67   84.27   83.20   84.00   82.67   82.93   84.80   83.47   81.33   85.87   81.07   81.33   84.27   83.73   83.73   83.20   84.27   81.87   84.27   83.73   84.53   84.80   84.00   82.93   84.00   84.00   83.20   85.33   82.93   83.73   83.47   84.80   83.47   82.93   84.00   81.87   82.67   81.87   82.67   82.13   82.93   81.87   81.60   82.93   85.07   84.00   82.67   82.40   85.07   82.67   85.33   81.60   83.47   81.33   84.80   82.93   84.80   83.73   83.47   83.20   84.27   84.00   82.67   84.00   84.53   82.40   84.27   85.07   87.47   83.47   85.87   85.60   82.67   85.33   88.53   88.00
    14: Sequence42    79.73   76.80   77.33   77.87   76.53   77.87   77.33   80.53   77.33   78.67   84.27   85.07   87.47  100.00   84.53   82.93   84.53   83.47   82.40   80.27   81.33   81.60   82.93   82.67   80.80   84.53   83.47   83.73   84.80   83.20   83.47   83.47   84.00   81.87   83.47   84.00   82.93   84.27   80.53   84.27   83.73   81.60   82.40   83.73   83.73   84.53   80.80   83.47   85.07   85.33   84.53   84.53   84.80   83.47   84.00   80.00   81.87   82.93   82.93   81.60   84.80   84.00   83.20   84.00   84.53   81.33   84.00   84.00   84.00   82.67   84.27   82.93   84.53   83.20   84.00   83.20   85.60   81.60   84.00   82.40   83.73   83.20   82.40   81.60   83.20   82.93   83.20   80.80   83.47   83.47   84.27   83.47   84.00   84.00   83.47   84.80   84.80   83.47   83.47   85.33   85.33   82.13   84.27   85.87   86.13   82.93   86.67   85.33   87.20   82.67   88.00   86.93
    15: Sequence35    78.13   79.20   77.07   78.13   77.87   77.33   79.20   75.73   77.33   78.67   85.07   83.73   86.40   84.53  100.00   86.93   85.87   83.73   82.93   84.80   82.40   82.93   83.20   83.47   83.20   80.80   82.93   82.93   82.40   83.47   83.73   82.93   84.00   81.07   82.13   81.07   82.40   83.20   79.73   82.13   81.87   82.13   84.00   83.47   80.27   83.20   82.67   80.53   85.60   81.60   83.47   80.53   82.67   81.07   83.73   82.67   84.53   84.27   82.40   80.00   84.27   85.07   82.93   84.80   85.60   83.47   81.60   83.20   84.80   82.93   84.80   82.93   85.33   80.27   83.73   81.87   81.87   83.20   82.67   82.40   85.07   82.93   81.87   81.87   85.33   85.60   84.00   83.20   85.87   83.47   85.33   83.47   82.40   85.60   85.33   86.67   84.80   85.87   82.40   82.93   81.60   81.87   84.27   86.40   87.20   83.20   83.73   82.93   82.93   85.60   87.73   86.93
    16: Sequence30    76.00   78.40   74.40   76.80   79.47   77.87   78.40   76.27   78.93   77.87   84.00   83.20   84.53   82.93   86.93  100.00   86.93   84.53   85.33   84.27   83.20   83.20   84.27   83.20   83.73   82.40   84.53   86.67   85.60   85.07   84.00   83.73   81.87   83.20   82.40   83.73   85.07   84.53   82.67   84.53   83.20   84.53   84.53   84.27   81.60   81.60   85.07   84.27   84.27   82.93   85.07   83.73   83.47   82.93   83.73   81.60   86.13   85.33   84.80   82.93   82.13   83.73   82.40   85.60   84.53   84.53   82.67   81.87   85.33   84.53   85.07   81.33   82.40   82.13   84.00   82.13   83.73   83.47   84.27   84.00   84.00   83.20   81.87   82.13   84.80   82.93   84.00   84.00   84.00   85.87   83.20   82.67   85.33   82.93   86.13   83.73   83.73   83.47   82.40   85.87   83.47   82.93   82.40   82.93   86.67   85.87   85.07   85.07   85.60   86.93   88.27   88.00
    17: Sequence76    77.60   76.53   77.60   75.20   77.87   78.40   78.40   78.40   77.87   76.00   82.67   81.07   83.47   84.53   85.87   86.93  100.00   84.80   86.13   85.33   82.67   84.27   81.07   84.00   83.47   82.93   83.47   85.07   86.13   85.07   84.80   85.60   84.80   82.67   82.93   83.47   81.07   83.73   82.13   81.87   80.27   82.93   84.80   85.60   82.93   82.40   81.07   82.40   82.40   82.67   84.53   85.07   82.67   82.40   85.87   81.87   83.73   82.93   82.40   82.13   83.73   84.80   84.00   84.00   85.87   82.40   81.07   80.27   83.20   82.93   84.27   80.80   84.27   81.60   84.00   82.67   83.20   83.73   84.80   84.80   83.47   82.40   82.13   82.67   83.20   84.00   82.13   83.20   86.13   84.00   83.73   82.40   84.53   84.00   85.33   84.80   82.67   84.00   83.20   85.87   83.47   82.40   82.13   84.53   85.33   83.47   84.27   84.53   84.80   86.93   87.73   87.47
    18: Sequence86    75.47   77.33   76.80   78.40   78.67   78.67   77.87   75.73   78.13   80.00   84.00   83.20   84.27   83.47   83.73   84.53   84.80  100.00   85.87   84.53   82.40   83.20   81.07   81.87   83.73   83.47   79.20   81.60   83.20   82.67   84.53   83.20   81.87   82.40   81.60   85.33   82.67   80.27   84.27   82.67   82.13   81.07   81.07   83.20   83.20   83.73   82.93   84.00   81.33   81.87   84.53   84.00   80.27   81.33   84.00   82.13   84.80   84.80   82.40   80.53   82.13   82.93   80.53   82.67   83.20   82.67   81.60   80.80   82.40   81.60   81.33   79.73   82.67   83.73   82.93   82.67   81.60   82.93   81.60   82.13   82.93   80.53   81.60   81.07   84.53   82.67   82.13   80.80   84.80   83.73   80.53   82.13   86.93   83.73   82.67   82.67   80.80   84.80   81.87   81.87   83.47   81.33   85.87   83.20   83.73   83.73   83.73   85.87   81.87   84.53   86.67   85.87
    19: Sequence48    76.53   76.27   77.33   80.00   78.67   77.07   77.87   77.07   78.67   77.33   83.47   82.93   83.20   82.40   82.93   85.33   86.13   85.87  100.00   86.40   82.93   85.07   81.87   81.87   84.27   82.93   82.93   83.47   81.87   81.33   84.00   84.27   82.13   80.00   82.40   84.53   81.87   83.73   84.27   83.73   82.67   83.73   85.33   85.33   82.13   83.73   82.67   82.67   82.13   83.20   85.07   85.07   80.53   83.47   82.93   82.93   85.07   83.73   84.53   83.20   85.33   84.53   82.40   83.73   82.93   82.67   79.47   81.07   81.60   84.27   82.13   79.47   80.53   82.13   81.33   81.60   81.33   82.67   83.20   82.67   84.53   83.47   84.00   80.80   84.27   81.87   81.87   81.87   82.67   83.47   81.87   81.60   83.20   85.07   83.73   81.60   82.67   84.80   84.00   83.47   82.93   81.60   83.73   83.47   83.47   86.40   82.93   84.27   82.40   84.53   87.47   86.93
    20: Sequence66    76.80   78.13   75.47   78.40   78.93   80.53   78.40   76.53   77.60   77.07   82.93   82.40   82.40   80.27   84.80   84.27   85.33   84.53   86.40  100.00   86.67   83.20   81.07   82.67   84.53   82.40   81.60   84.53   82.40   85.33   83.47   84.27   81.33   82.13   84.53   84.27   83.73   84.80   81.87   83.20   81.87   83.47   83.73   85.07   81.87   84.00   82.93   82.93   82.40   84.00   84.00   84.53   82.13   84.27   84.00   83.73   84.53   82.13   81.60   82.67   82.13   82.40   83.20   83.73   84.27   83.47   80.27   81.60   82.93   85.87   82.67   80.80   82.67   81.87   84.53   80.80   81.33   81.60   84.27   81.87   82.40   81.60   84.00   81.07   85.33   85.07   82.13   82.93   83.47   82.67   83.73   82.40   83.20   84.80   84.80   86.40   83.20   82.13   84.53   83.73   82.40   82.93   84.53   84.27   83.47   85.33   81.87   83.73   83.20   85.87   87.47   89.60
    21: Sequence99    78.67   78.93   76.53   77.60   78.67   78.93   74.40   76.00   79.73   76.00   83.73   81.33   81.07   81.33   82.40   83.20   82.67   82.40   82.93   86.67  100.00   80.80   82.93   81.60   82.93   83.47   83.73   84.80   84.00   82.93   81.33   80.53   80.53   80.00   78.40   82.67   84.27   80.27   79.73   80.80   82.67   81.87   82.13   85.33   82.40   80.27   82.67   82.13   81.87   81.87   84.00   84.00   81.87   83.47   83.47   83.20   83.20   84.53   82.93   82.40   80.80   82.93   80.27   85.07   84.80   84.80   81.60   80.53   81.07   82.67   79.73   80.80   82.93   80.80   84.00   80.80   80.53   80.80   80.53   84.27   83.73   80.80   81.87   80.80   84.27   81.60   81.87   80.80   82.93   81.87   84.00   80.80   84.00   81.87   81.07   84.53   84.00   82.67   83.73   84.53   82.40   83.73   83.20   85.87   82.67   84.00   84.53   83.20   85.60   83.73   86.67   86.67
    22: Sequence85    77.60   77.60   76.80   77.07   77.60   74.93   77.87   76.80   78.93   78.13   84.00   83.47   85.07   81.60   82.93   83.20   84.27   83.20   85.07   83.20   80.80  100.00   81.60   85.33   83.47   85.33   84.80   82.67   83.47   83.20   85.33   80.80   81.33   82.40   81.33   82.93   82.93   80.80   84.27   83.47   81.33   82.40   83.47   81.07   82.93   84.80   82.67   81.60   84.53   82.40   82.67   82.93   83.20   81.33   81.60   81.33   84.27   82.13   84.00   83.73   82.93   80.53   84.27   82.13   81.87   79.20   82.40   81.60   82.40   82.40   81.07   80.80   80.53   81.87   82.40   82.67   82.13   83.20   83.47   83.20   82.67   84.53   86.67   83.73   84.00   83.20   83.20   81.87   83.20   81.60   82.40   84.27   82.93   83.73   82.93   82.93   81.07   84.53   82.67   81.60   84.00   82.13   86.40   83.20   85.07   84.53   81.87   83.73   82.93   84.53   87.20   87.73
    23: Sequence52    80.00   81.33   78.13   77.87   79.47   77.60   74.13   78.93   77.87   76.27   82.40   82.40   83.73   82.93   83.20   84.27   81.07   81.07   81.87   81.07   82.93   81.60  100.00   83.47   82.40   83.47   81.33   86.13   84.27   82.93   83.73   83.20   82.93   83.73   81.87   81.87   82.13   84.80   80.00   83.20   83.73   83.73   84.00   82.93   82.67   81.60   83.47   82.13   83.20   82.67   83.73   82.93   83.20   84.53   81.07   82.40   82.93   82.40   84.27   80.53   81.07   83.20   81.33   82.93   85.60   83.47   83.73   81.33   84.27   82.93   83.73   81.87   82.67   82.93   85.33   84.80   84.00   85.33   84.53   84.53   84.00   82.13   82.67   83.20   83.20   86.13   87.73   84.00   84.80   85.60   83.47   83.47   83.20   82.93   84.53   81.87   83.47   83.73   85.33   83.20   84.80   84.53   84.00   83.47   83.73   83.20   85.87   82.93   84.80   83.47   87.47   88.00
    24: Sequence53    78.93   79.73   79.20   75.20   78.40   79.73   77.07   78.67   79.20   78.13   80.80   81.60   81.33   82.67   83.47   83.20   84.00   81.87   81.87   82.67   81.60   85.33   83.47  100.00   86.40   86.13   82.67   83.47   85.87   85.87   82.67   83.47   84.80   83.20   81.87   83.47   85.07   83.73   82.40   82.67   83.73   84.00   84.80   81.33   81.33   84.27   85.33   82.93   81.60   80.27   82.40   81.07   85.07   84.80   83.20   82.67   83.20   82.67   83.47   84.53   82.40   82.67   84.00   82.40   83.73   83.73   81.33   80.80   82.67   85.33   84.00   82.40   85.07   83.73   82.93   81.60   84.00   86.13   84.27   84.53   83.20   81.87   83.47   85.07   82.40   84.27   84.53   84.27   85.33   82.40   83.73   82.13   81.33   84.00   85.33   84.80   83.73   81.87   82.40   83.20   82.93   84.27   85.07   83.73   81.87   82.93   84.00   84.00   85.33   84.80   87.20   86.40
    25: Sequence96    77.60   79.20   77.33   76.53   78.40   78.67   78.13   78.13   79.20   77.33   82.93   83.47   82.67   80.80   83.20   83.73   83.47   83.73   84.27   84.53   82.93   83.47   82.40   86.40  100.00   83.20   82.93   83.47   85.60   85.33   83.47   82.13   80.80   80.53   81.87   84.00   84.80   81.87   82.67   81.87   82.67   84.27   81.33   82.67   81.07   83.47   83.73   85.07   81.87   81.60   81.33   81.07   83.20   80.80   82.13   81.60   85.33   84.80   85.60   82.93   80.80   81.07   81.87   83.73   82.40   83.73   80.80   83.20   84.27   84.27   83.20   82.67   83.73   83.20   80.27   82.67   79.73   83.20   84.80   83.73   84.53   80.80   83.47   80.53   83.20   83.20   82.67   81.60   81.87   82.93   84.27   81.60   82.40   84.27   83.73   82.67   83.20   84.27   81.07   80.80   82.93   82.93   83.73   85.33   81.87   85.33   82.93   83.73   82.13   84.27   86.40   86.40
    26: Sequence24    78.13   76.80   76.27   77.87   75.73   78.93   79.73   78.67   79.47   79.47   83.47   82.40   83.20   84.53   80.80   82.40   82.93   83.47   82.93   82.40   83.47   85.33   83.47   86.13   83.20  100.00   82.67   83.20   84.53   86.40   82.93   83.47   82.40   82.40   82.13   85.07   84.27   82.13   85.87   82.67   82.40   84.53   82.13   82.67   84.80   83.73   82.93   83.73   81.60   85.60   81.87   82.67   84.00   84.80   82.67   80.80   83.47   85.60   82.13   81.60   83.20   81.33   83.47   83.47   82.93   82.67   84.27   77.87   83.20   82.40   82.13   81.60   80.53   83.73   81.87   84.53   83.47   82.13   81.87   82.93   82.67   82.13   85.60   81.07   85.07   82.67   81.33   81.07   83.20   82.13   84.00   82.67   85.87   86.13   84.80   83.73   82.93   82.67   83.47   84.53   87.20   84.27   87.20   83.20   84.27   84.53   84.27   84.27   84.00   84.53   87.47   87.20
    27: Sequence74    76.53   78.67   79.20   78.13   75.73   76.00   78.40   80.27   80.80   78.67   85.60   82.40   82.67   83.47   82.93   84.53   83.47   79.20   82.93   81.60   83.73   84.80   81.33   82.67   82.93   82.67  100.00   86.40   85.60   84.53   82.93   82.40   84.27   81.87   81.07   82.67   82.67   83.20   81.87   82.40   82.13   82.67   81.07   82.40   82.13   83.47   81.60   82.13   82.93   83.73   81.07   82.40   82.93   83.73   82.93   81.33   83.47   83.47   84.00   81.60   84.00   82.13   84.27   81.60   81.87   83.20   83.73   83.73   84.27   84.80   83.73   84.80   83.20   83.20   82.67   82.13   81.07   82.93   81.33   84.80   85.07   84.00   82.67   83.73   83.20   82.40   81.87   82.93   81.33   81.60   83.47   82.67   82.13   82.13   85.07   80.53   82.40   83.20   81.07   84.27   84.00   83.20   82.40   85.07   84.00   84.00   84.27   84.00   84.80   82.40   86.67   85.87
    28: Sequence62    80.27   78.67   76.27   76.53   78.13   76.80   74.93   77.07   78.13   76.53   83.73   82.40   84.27   83.73   82.93   86.67   85.07   81.60   83.47   84.53   84.80   82.67   86.13   83.47   83.47   83.20   86.40  100.00   86.40   85.60   83.20   83.73   83.47   83.73   82.67   83.47   84.27   82.67   81.87   82.13   84.53   84.53   80.80   82.93   82.40   83.73   82.40   85.07   82.40   81.60   82.93   83.20   83.47   84.00   85.60   82.93   85.07   82.93   84.27   83.73   82.67   83.73   84.80   84.00   85.60   84.53   84.80   81.87   84.27   84.53   83.73   81.87   83.47   84.00   84.27   82.40   83.20   85.07   84.53   84.00   83.47   81.87   83.20   83.20   82.93   84.80   85.60   82.93   83.47   84.53   85.07   82.40   84.53   84.00   84.80   84.00   83.47   83.47   84.00   84.53   85.33   85.33   84.80   84.27   84.53   84.53   85.07   84.00   85.87   86.67   88.27   89.33
    29: Sequence23    80.00   78.67   77.07   76.00   79.73   80.00   77.60   79.20   79.47   76.53   83.73   82.93   82.67   84.80   82.40   85.60   86.13   83.20   81.87   82.40   84.00   83.47   84.27   85.87   85.60   84.53   85.60   86.40  100.00   88.00   84.00   81.07   82.67   83.20   80.27   84.00   85.87   83.73   82.40   83.20   83.73   83.20   81.87   84.00   81.87   84.53   83.73   85.33   84.00   83.73   82.13   84.00   84.80   85.07   84.80   81.07   84.00   85.07   84.00   84.00   82.13   83.73   84.00   86.40   84.80   82.40   84.27   80.80   84.00   82.40   84.27   84.00   85.60   82.40   84.80   85.87   84.00   82.67   83.47   85.60   83.73   81.07   83.20   85.07   82.93   83.20   82.40   83.73   82.40   84.00   85.33   82.67   85.33   84.80   86.93   83.73   83.73   83.47   83.47   85.87   84.53   83.73   84.00   86.13   82.93   82.13   84.27   84.00   85.60   83.20   88.53   88.00
    30: Sequence60    78.13   77.60   73.87   77.33   78.40   78.13   79.47   79.47   77.07   78.67   83.47   83.47   83.20   83.20   83.47   85.07   85.07   82.67   81.33   85.33   82.93   83.20   82.93   85.87   85.33   86.40   84.53   85.60   88.00  100.00   81.87   81.60   80.00   84.00   82.13   84.00   83.47   84.27   83.20   81.87   81.33   84.27   82.13   81.87   83.20   85.60   84.27   82.67   82.67   83.47   82.93   82.40   85.33   82.67   82.93   80.80   84.00   86.40   83.73   81.87   82.13   81.33   84.80   82.93   83.73   84.27   83.20   82.13   85.33   86.13   82.67   83.20   84.27   83.47   82.67   82.67   82.13   82.67   85.07   85.33   81.87   82.13   84.00   84.00   81.33   85.60   83.20   83.47   83.20   83.73   83.73   83.47   84.53   84.00   85.87   82.93   84.53   82.67   81.33   86.67   84.00   84.80   85.87   83.47   84.27   83.47   84.00   85.33   85.87   84.53   88.00   87.73
    31: Sequence71    80.00   79.47   78.13   75.73   79.73   78.93   76.27   78.13   79.73   78.67   84.53   84.27   84.27   83.47   83.73   84.00   84.80   84.53   84.00   83.47   81.33   85.33   83.73   82.67   83.47   82.93   82.93   83.20   84.00   81.87  100.00   85.07   84.27   83.47   81.07   85.07   82.40   83.20   83.20   84.00   81.07   80.53   80.80   81.60   80.53   83.73   81.60   83.47   83.20   81.87   83.73   83.73   82.40   82.40   81.87   81.60   82.40   81.33   80.80   80.27   81.60   82.67   80.00   81.87   83.20   82.93   82.40   82.13   82.13   82.67   82.40   80.27   81.07   81.33   81.87   82.13   83.47   81.87   83.47   81.60   81.60   78.93   84.53   82.40   85.87   84.27   82.93   82.67   83.47   84.27   80.80   82.13   86.13   82.67   83.20   84.00   82.40   84.27   82.40   83.47   81.87   82.13   82.93   82.40   84.80   85.07   85.87   82.93   86.13   83.47   87.47   86.67
    32: Sequence73    76.27   76.27   78.67   76.00   77.87   80.00   74.93   78.67   78.67   79.47   81.60   81.87   83.20   83.47   82.93   83.73   85.60   83.20   84.27   84.27   80.53   80.80   83.20   83.47   82.13   83.47   82.40   83.73   81.07   81.60   85.07  100.00   87.47   84.00   84.00   82.93   82.40   84.00   82.13   83.47   83.73   80.53   81.60   82.67   83.20   81.60   81.87   83.47   81.87   83.47   83.73   82.40   82.93   83.20   83.20   84.27   82.93   82.67   84.00   81.33   83.20   82.93   80.53   84.27   85.07   83.73   82.40   83.20   83.73   83.73   82.67   80.80   81.87   80.80   81.87   82.93   84.80   84.80   84.00   82.93   82.40   84.00   81.87   82.40   83.47   84.80   82.67   83.47   84.80   85.60   82.93   84.00   83.20   84.00   82.40   82.67   83.73   84.27   81.87   82.93   84.27   83.47   82.93   84.80   83.20   84.27   85.87   85.33   83.20   85.60   87.47   88.53
    33: Sequence77    75.73   78.40   77.07   77.33   77.87   77.33   74.67   80.80   77.87   78.67   81.60   81.60   85.07   84.00   84.00   81.87   84.80   81.87   82.13   81.33   80.53   81.33   82.93   84.80   80.80   82.40   84.27   83.47   82.67   80.00   84.27   87.47  100.00   81.60   81.07   81.33   82.40   81.60   81.07   83.73   83.47   80.27   83.20   81.60   81.07   81.33   82.13   83.47   82.67   84.53   80.80   82.40   81.07   82.13   84.27   84.53   84.27   81.87   84.00   82.93   84.53   84.80   81.60   81.60   82.93   84.53   81.33   84.80   82.40   82.13   85.60   81.60   82.40   80.80   82.93   81.33   83.47   83.20   81.60   81.87   85.07   82.40   81.60   82.67   82.67   84.00   84.53   82.93   84.80   84.53   83.73   82.67   83.20   83.20   83.20   83.20   82.93   81.87   82.67   81.60   82.13   82.67   81.60   85.07   84.27   81.60   84.80   85.07   82.13   82.13   86.67   85.33
    34: Sequence92    78.67   78.13   77.87   75.20   80.27   76.80   76.00   77.87   77.33   77.60   82.40   81.33   83.20   81.87   81.07   83.20   82.67   82.40   80.00   82.13   80.00   82.40   83.73   83.20   80.53   82.40   81.87   83.73   83.20   84.00   83.47   84.00   81.60  100.00   83.73   84.53   84.80   85.60   84.80   84.80   82.67   81.60   81.33   80.53   83.47   82.13   80.00   83.47   82.13   82.93   82.13   84.00   83.73   83.47   80.53   82.13   82.13   82.13   81.33   80.80   80.27   80.27   81.33   82.13   82.67   81.87   85.33   81.07   83.73   84.80   82.67   80.53   82.67   81.87   81.60   82.67   83.47   85.07   82.67   82.67   82.93   84.53   80.53   83.47   80.80   85.33   83.73   86.13   83.47   84.27   82.13   83.47   83.47   84.27   85.07   83.73   82.93   82.13   82.93   81.87   83.47   83.20   80.80   81.87   83.47   83.20   85.07   82.93   83.47   85.60   86.67   86.93
    35: Sequence97    78.13   75.20   76.27   80.80   77.07   76.80   78.13   77.07   75.47   76.53   81.33   81.60   82.93   83.47   82.13   82.40   82.93   81.60   82.40   84.53   78.40   81.33   81.87   81.87   81.87   82.13   81.07   82.67   80.27   82.13   81.07   84.00   81.07   83.73  100.00   85.33   83.73   85.87   83.20   84.80   82.93   83.20   83.20   83.47   85.07   82.67   82.67   82.13   83.47   83.47   82.93   84.53   83.20   83.47   83.73   82.13   81.60   81.60   81.07   81.07   82.40   84.00   80.00   83.20   80.53   82.67   84.53   80.80   83.20   83.73   82.67   82.93   81.60   82.67   82.40   83.20   83.47   83.47   85.87   83.47   84.27   84.00   84.80   86.13   82.13   84.00   83.73   84.00   80.00   81.60   84.00   83.20   83.73   83.20   85.60   82.93   84.27   82.13   84.27   82.13   84.53   85.33   83.73   83.47   84.00   84.80   82.40   82.40   82.93   84.27   86.93   86.67
    36: Sequence81    80.00   76.27   76.80   78.13   76.27   78.93   74.67   76.53   77.60   79.20   84.80   82.40   83.20   84.00   81.07   83.73   83.47   85.33   84.53   84.27   82.67   82.93   81.87   83.47   84.00   85.07   82.67   83.47   84.00   84.00   85.07   82.93   81.33   84.53   85.33  100.00   85.87   84.27   84.80   82.13   81.87   84.00   82.40   82.93   85.07   84.00   82.13   85.07   82.67   84.00   84.00   84.27   84.00   85.87   82.93   81.33   82.67   84.27   81.60   81.33   84.00   83.47   80.80   82.67   83.47   83.47   84.00   81.33   85.33   83.20   80.80   82.40   82.40   84.80   83.47   80.80   82.93   81.60   83.73   84.80   83.20   81.33   84.00   82.13   84.80   82.40   82.13   83.20   83.20   83.20   81.60   83.47   85.07   84.00   84.53   84.27   84.00   81.07   83.20   83.73   81.60   83.73   82.93   82.93   83.20   86.40   83.47   85.07   83.47   85.60   87.47   87.47
    37: Sequence38    79.47   76.80   75.20   76.00   78.13   79.47   77.60   77.33   77.87   78.93   84.80   81.07   82.67   82.93   82.40   85.07   81.07   82.67   81.87   83.73   84.27   82.93   82.13   85.07   84.80   84.27   82.67   84.27   85.87   83.47   82.40   82.40   82.40   84.80   83.73   85.87  100.00   88.53   84.53   84.53   85.07   82.67   82.93   82.93   83.73   82.40   82.13   85.87   83.47   83.73   82.13   84.27   83.47   85.33   84.53   81.33   84.80   84.53   82.13   82.93   81.60   81.33   82.40   84.00   81.60   82.93   83.20   82.40   84.53   82.13   82.67   82.13   81.87   81.87   84.80   81.60   85.60   82.93   85.33   82.40   82.40   83.20   81.07   84.00   85.07   82.13   83.73   85.07   81.33   80.80   84.80   84.27   86.13   85.87   84.53   84.00   84.00   84.80   84.80   83.73   83.47   85.33   82.93   85.60   84.00   84.00   84.80   84.53   83.47   83.20   87.73   86.13
    38: Sequence69    77.87   76.53   75.73   77.07   78.13   78.67   79.47   77.87   79.47   78.67   83.47   81.07   84.27   84.27   83.20   84.53   83.73   80.27   83.73   84.80   80.27   80.80   84.80   83.73   81.87   82.13   83.20   82.67   83.73   84.27   83.20   84.00   81.60   85.60   85.87   84.27   88.53  100.00   84.00   83.47   82.93   83.20   86.40   83.47   82.40   83.20   81.33   82.13   83.20   84.00   82.13   84.53   83.47   85.87   81.87   82.13   82.40   82.13   81.87   81.60   83.73   80.53   81.07   83.47   81.33   82.93   84.27   81.87   85.60   82.93   83.73   82.67   82.93   83.47   82.93   82.40   85.87   83.20   83.47   83.47   82.13   83.20   81.87   85.07   84.27   83.73   82.40   85.60   81.33   82.13   82.40   86.40   84.27   84.80   85.33   84.00   84.27   84.00   85.60   84.00   83.73   84.80   84.00   84.27   83.47   82.93   84.80   84.80   83.20   83.47   87.73   86.13
    39: Sequence94    78.13   77.60   76.53   77.60   78.40   76.80   78.13   77.60   79.73   78.40   83.47   82.67   83.20   80.53   79.73   82.67   82.13   84.27   84.27   81.87   79.73   84.27   80.00   82.40   82.67   85.87   81.87   81.87   82.40   83.20   83.20   82.13   81.07   84.80   83.20   84.80   84.53   84.00  100.00   85.87   81.87   82.67   81.07   82.13   83.20   85.60   83.73   83.47   79.20   81.60   83.47   81.60   80.27   84.27   83.20   81.07   84.80   84.80   83.20   82.67   81.60   80.00   83.73   82.93   80.00   81.33   82.93   82.67   82.93   82.93   81.07   81.07   80.80   82.93   80.00   84.27   84.00   81.07   81.87   81.60   80.00   80.53   82.13   80.27   83.73   80.53   80.53   80.80   82.93   81.87   82.40   81.60   85.60   85.33   82.40   81.87   81.60   83.20   82.93   81.87   83.73   83.20   84.00   82.40   82.93   84.53   82.67   83.73   83.20   83.73   86.13   85.60
    40: Sequence70    78.93   76.80   76.27   75.73   78.13   79.47   75.73   79.73   79.47   76.27   83.73   83.73   84.00   84.27   82.13   84.53   81.87   82.67   83.73   83.20   80.80   83.47   83.20   82.67   81.87   82.67   82.40   82.13   83.20   81.87   84.00   83.47   83.73   84.80   84.80   82.13   84.53   83.47   85.87  100.00   86.40   82.93   81.60   81.60   82.93   85.33   80.80   84.27   82.40   84.00   83.47   84.00   81.87   82.13   83.20   85.60   83.47   83.20   83.47   82.67   82.40   81.87   82.40   84.00   82.40   80.80   82.93   83.47   84.27   85.60   85.33   81.60   81.33   81.33   82.40   84.53   83.73   83.73   85.60   81.87   83.47   83.73   81.87   84.00   83.47   84.00   83.73   85.07   81.60   83.47   84.53   85.33   82.67   82.93   84.27   82.40   84.00   82.13   82.13   85.60   85.33   84.80   82.93   83.47   82.40   82.93   85.07   84.27   85.07   83.20   88.00   87.20
    41: Sequence84    76.27   77.33   74.93   76.00   78.93   76.80   76.00   76.53   79.73   79.73   82.40   82.93   82.67   83.73   81.87   83.20   80.27   82.13   82.67   81.87   82.67   81.33   83.73   83.73   82.67   82.40   82.13   84.53   83.73   81.33   81.07   83.73   83.47   82.67   82.93   81.87   85.07   82.93   81.87   86.40  100.00   82.67   80.00   82.13   80.27   81.33   83.20   84.80   81.60   83.20   82.40   81.87   82.40   82.67   83.20   84.27   85.07   83.20   84.53   82.40   81.33   83.20   80.53   84.80   82.40   82.67   82.67   84.27   83.73   82.40   82.13   83.20   82.40   82.93   83.20   84.53   84.80   82.67   81.60   81.07   82.67   84.27   80.00   84.00   81.60   82.13   83.20   82.93   84.27   86.40   85.07   84.00   81.60   82.40   81.87   83.47   85.07   83.20   82.67   83.47   84.80   82.40   84.27   85.33   81.33   83.20   87.47   81.07   84.27   85.33   86.93   87.73
    42: Sequence98    78.93   77.07   75.47   77.87   78.40   77.07   77.87   77.87   77.33   79.73   82.93   84.00   82.93   81.60   82.13   84.53   82.93   81.07   83.73   83.47   81.87   82.40   83.73   84.00   84.27   84.53   82.67   84.53   83.20   84.27   80.53   80.53   80.27   81.60   83.20   84.00   82.67   83.20   82.67   82.93   82.67  100.00   84.80   83.73   81.33   81.87   85.07   82.93   79.47   81.87   82.93   84.80   83.20   82.67   82.67   83.47   84.27   84.53   84.80   84.53   80.27   83.47   83.73   84.27   83.73   82.13   84.53   79.47   83.20   84.00   82.93   82.40   82.40   82.13   82.13   82.40   80.27   82.40   84.00   81.60   81.33   80.80   82.67   79.47   81.60   80.27   81.33   80.00   83.47   83.20   80.80   80.53   81.60   82.40   83.20   84.00   80.80   81.60   80.80   84.53   85.33   84.27   84.27   81.07   84.53   82.40   82.13   83.47   81.33   83.47   86.40   86.67
    43: Sequence47    77.07   76.53   75.20   79.47   80.00   77.33   79.20   78.40   78.13   74.67   84.27   82.93   84.80   82.40   84.00   84.53   84.80   81.07   85.33   83.73   82.13   83.47   84.00   84.80   81.33   82.13   81.07   80.80   81.87   82.13   80.80   81.60   83.20   81.33   83.20   82.40   82.93   86.40   81.07   81.60   80.00   84.80  100.00   86.40   83.47   82.40   82.93   80.00   83.73   82.93   86.40   85.33   82.40   85.07   84.53   83.73   86.40   82.67   84.27   82.67   84.27   84.53   82.93   83.73   82.67   84.27   82.13   80.27   84.27   84.00   83.20   79.47   82.40   81.60   83.73   83.47   83.73   85.07   84.80   85.33   85.60   85.60   84.53   85.87   81.33   83.47   82.93   84.80   85.33   84.27   82.40   84.27   81.07   81.87   82.67   84.80   81.07   83.73   85.07   85.33   84.27   84.80   82.93   85.60   85.07   82.13   82.40   82.93   83.47   84.00   87.20   85.87
    44: Sequence58    77.87   76.27   78.13   76.80   78.93   80.53   77.60   76.00   78.67   75.47   83.20   83.73   83.47   83.73   83.47   84.27   85.60   83.20   85.33   85.07   85.33   81.07   82.93   81.33   82.67   82.67   82.40   82.93   84.00   81.87   81.60   82.67   81.60   80.53   83.47   82.93   82.93   83.47   82.13   81.60   82.13   83.73   86.40  100.00   80.80   83.20   84.80   81.33   84.53   83.47   84.53   85.87   84.27   84.53   83.47   82.93   85.60   84.27   83.47   80.00   83.47   85.60   83.20   86.13   85.60   84.00   82.93   82.67   82.40   82.67   82.67   79.73   83.20   81.60   83.47   82.93   82.40   82.40   81.87   84.00   82.13   83.47   84.80   83.20   84.00   84.27   84.27   82.93   84.00   83.73   84.80   81.87   83.47   84.00   83.20   84.00   84.80   82.93   84.80   84.00   82.40   82.93   82.40   85.87   86.67   84.27   83.20   83.20   86.67   82.40   88.00   86.67
    45: Sequence91    77.87   75.73   78.40   80.00   77.07   75.73   76.80   80.00   76.80   76.27   85.07   82.13   81.33   83.73   80.27   81.60   82.93   83.20   82.13   81.87   82.40   82.93   82.67   81.33   81.07   84.80   82.13   82.40   81.87   83.20   80.53   83.20   81.07   83.47   85.07   85.07   83.73   82.40   83.20   82.93   80.27   81.33   83.47   80.80  100.00   81.60   82.13   83.73   80.80   82.40   84.80   82.93   80.53   86.40   83.47   80.80   82.67   80.53   81.60   81.60   83.73   83.20   82.13   80.27   80.53   82.67   82.40   81.33   84.27   82.13   81.60   80.80   82.67   83.20   83.73   84.80   85.07   82.67   84.80   84.80   85.33   84.80   85.33   82.93   82.67   82.93   83.73   84.53   81.87   83.20   83.20   85.07   84.00   83.47   83.20   81.33   82.40   82.93   85.33   82.93   85.87   85.60   84.80   85.33   81.87   84.80   84.00   85.60   83.20   84.27   86.93   85.33
    46: Sequence61    78.67   76.00   77.33   76.27   78.13   78.13   76.53   78.67   76.80   77.60   83.47   82.13   85.87   84.53   83.20   81.60   82.40   83.73   83.73   84.00   80.27   84.80   81.60   84.27   83.47   83.73   83.47   83.73   84.53   85.60   83.73   81.60   81.33   82.13   82.67   84.00   82.40   83.20   85.60   85.33   81.33   81.87   82.40   83.20   81.60  100.00   82.93   80.80   81.87   82.93   81.33   85.87   82.67   82.67   81.07   82.13   84.00   84.53   84.00   82.93   84.80   80.80   85.33   83.73   82.67   83.47   80.80   82.13   84.27   85.07   85.33   82.40   83.20   83.73   83.47   82.67   83.47   83.20   84.53   82.93   81.07   82.67   82.93   83.20   85.07   83.47   82.93   81.60   83.47   82.13   82.40   84.00   81.07   82.93   84.00   85.07   82.40   82.93   82.13   84.53   83.73   83.20   85.60   85.07   83.73   81.87   84.27   86.93   82.67   84.00   88.27   87.47
    47: Sequence80    75.73   78.40   75.20   78.67   79.73   77.07   78.40   77.33   78.40   76.00   79.47   80.53   81.07   80.80   82.67   85.07   81.07   82.93   82.67   82.93   82.67   82.67   83.47   85.33   83.73   82.93   81.60   82.40   83.73   84.27   81.60   81.87   82.13   80.00   82.67   82.13   82.13   81.33   83.73   80.80   83.20   85.07   82.93   84.80   82.13   82.93  100.00   81.33   81.60   82.40   81.07   82.13   83.73   84.00   84.80   84.00   85.87   85.87   86.40   82.67   81.60   84.27   82.67   84.00   82.13   83.20   80.80   82.40   83.47   80.53   80.00   82.67   82.93   82.40   82.40   82.13   84.53   83.47   85.60   84.00   82.13   81.60   83.47   81.33   82.40   80.80   83.20   82.40   84.27   84.00   83.47   85.07   82.67   81.87   82.40   84.27   82.67   81.60   83.47   84.00   83.47   81.87   84.80   83.47   84.00   85.07   83.47   83.47   82.93   84.00   86.67   86.67
    48: Sequence59    79.47   78.67   76.53   76.53   77.33   81.07   76.80   79.47   78.93   77.87   85.33   83.20   81.33   83.47   80.53   84.27   82.40   84.00   82.67   82.93   82.13   81.60   82.13   82.93   85.07   83.73   82.13   85.07   85.33   82.67   83.47   83.47   83.47   83.47   82.13   85.07   85.87   82.13   83.47   84.27   84.80   82.93   80.00   81.33   83.73   80.80   81.33  100.00   82.67   84.27   80.53   83.47   82.13   82.67   85.07   83.20   86.40   82.67   84.00   83.73   82.67   82.13   82.40   83.73   85.07   81.07   83.20   83.47   85.60   82.93   83.47   82.40   84.53   82.13   83.47   84.80   82.40   82.93   82.67   81.60   80.80   79.73   81.60   82.13   82.67   84.00   83.20   82.93   85.33   85.07   83.47   82.40   84.27   82.93   84.27   83.47   82.13   85.60   83.73   84.53   83.73   84.00   84.00   82.93   83.47   84.27   86.13   82.93   85.07   85.07   87.47   86.93
    49: Sequence49    80.80   77.60   76.27   76.00   77.33   76.27   79.47   80.00   75.73   77.60   84.00   82.93   84.27   85.07   85.60   84.27   82.40   81.33   82.13   82.40   81.87   84.53   83.20   81.60   81.87   81.60   82.93   82.40   84.00   82.67   83.20   81.87   82.67   82.13   83.47   82.67   83.47   83.20   79.20   82.40   81.60   79.47   83.73   84.53   80.80   81.87   81.60   82.67  100.00   85.87   81.87   82.93   86.13   82.67   84.00   81.33   81.87   83.20   81.60   80.27   84.27   83.20   82.13   82.40   84.27   82.13   82.67   82.40   82.67   83.47   81.87   83.73   84.80   83.20   84.27   82.93   83.20   82.67   84.53   82.93   85.07   84.00   82.67   86.13   82.67   82.40   84.00   85.60   81.87   82.13   85.60   81.60   81.87   84.27   83.73   82.40   85.07   84.00   84.27   84.27   83.20   85.33   81.60   85.87   85.07   83.20   82.93   80.00   84.53   84.27   88.27   87.47
    50: Sequence40    77.33   77.87   76.27   76.27   78.40   77.60   76.27   80.00   76.53   77.07   84.53   84.80   83.73   85.33   81.60   82.93   82.67   81.87   83.20   84.00   81.87   82.40   82.67   80.27   81.60   85.60   83.73   81.60   83.73   83.47   81.87   83.47   84.53   82.93   83.47   84.00   83.73   84.00   81.60   84.00   83.20   81.87   82.93   83.47   82.40   82.93   82.40   84.27   85.87  100.00   82.93   84.00   85.07   85.87   84.00   82.67   85.07   84.00   82.67   82.40   83.47   84.00   83.73   83.47   85.07   85.07   83.20   84.80   85.33   84.53   84.53   83.73   82.93   82.40   82.67   83.47   83.20   81.87   84.00   82.40   83.20   85.07   85.07   82.93   81.60   83.20   81.87   84.53   84.00   84.80   85.07   82.93   84.53   85.87   85.07   83.20   85.60   82.93   85.60   85.60   82.67   84.53   82.93   84.27   85.60   83.47   85.33   84.53   82.93   84.53   88.53   88.80
    51: Sequence28    77.87   75.20   76.27   79.20   80.00   77.87   76.27   76.53   78.67   78.13   82.67   81.87   83.73   84.53   83.47   85.07   84.53   84.53   85.07   84.00   84.00   82.67   83.73   82.40   81.33   81.87   81.07   82.93   82.13   82.93   83.73   83.73   80.80   82.13   82.93   84.00   82.13   82.13   83.47   83.47   82.40   82.93   86.40   84.53   84.80   81.33   81.07   80.53   81.87   82.93  100.00   83.47   83.73   85.87   85.07   82.93   85.87   85.60   84.27   83.47   83.47   84.80   83.47   84.27   83.47   84.80   84.00   81.60   82.40   84.27   81.07   82.40   81.60   82.13   83.47   82.93   84.80   81.07   82.40   83.73   84.53   82.93   84.27   81.60   82.93   83.20   82.13   84.27   85.07   85.07   83.20   82.67   84.27   83.73   82.67   84.53   84.53   83.20   84.80   86.67   84.27   84.27   83.47   85.07   86.13   84.27   84.53   83.73   85.60   85.87   88.00   86.93
    52: Sequence16    78.13   76.80   76.27   76.53   80.27   78.93   77.87   80.00   75.73   77.87   83.73   82.13   83.20   84.53   80.53   83.73   85.07   84.00   85.07   84.53   84.00   82.93   82.93   81.07   81.07   82.67   82.40   83.20   84.00   82.40   83.73   82.40   82.40   84.00   84.53   84.27   84.27   84.53   81.60   84.00   81.87   84.80   85.33   85.87   82.93   85.87   82.13   83.47   82.93   84.00   83.47  100.00   83.47   83.73   84.00   85.60   84.80   85.33   86.13   81.07   82.13   83.73   83.20   84.80   84.27   83.20   83.73   80.80   82.93   83.73   80.53   80.27   84.27   84.53   83.47   84.00   84.00   84.53   84.53   81.60   81.87   82.40   81.33   83.73   82.13   81.87   81.87   83.20   85.87   84.27   83.73   84.00   82.67   82.67   84.27   86.13   82.40   83.47   82.93   85.60   84.00   85.60   83.20   82.93   85.33   82.67   86.93   85.33   82.93   83.47   87.73   88.00
    53: Sequence57    77.33   76.53   78.13   76.80   77.87   76.80   76.53   78.40   77.33   76.27   81.33   82.93   84.27   84.80   82.67   83.47   82.67   80.27   80.53   82.13   81.87   83.20   83.20   85.07   83.20   84.00   82.93   83.47   84.80   85.33   82.40   82.93   81.07   83.73   83.20   84.00   83.47   83.47   80.27   81.87   82.40   83.20   82.40   84.27   80.53   82.67   83.73   82.13   86.13   85.07   83.73   83.47  100.00   84.00   84.53   82.67   81.33   86.40   84.27   81.60   82.93   81.87   84.53   85.33   84.53   84.80   85.07   83.20   84.27   82.40   81.87   82.40   82.67   83.20   82.13   83.20   84.27   81.60   82.93   83.47   83.20   85.07   85.07   82.13   83.47   82.40   84.53   84.27   83.73   82.93   85.60   82.13   81.60   83.73   84.00   83.73   85.87   84.00   83.20   84.80   81.33   85.07   84.00   83.73   84.00   84.27   85.07   82.67   83.73   84.53   88.00   87.47
    54: Sequence44    79.47   76.53   77.33   78.67   79.47   78.13   76.53   76.80   78.67   75.20   82.13   80.80   81.87   83.47   81.07   82.93   82.40   81.33   83.47   84.27   83.47   81.33   84.53   84.80   80.80   84.80   83.73   84.00   85.07   82.67   82.40   83.20   82.13   83.47   83.47   85.87   85.33   85.87   84.27   82.13   82.67   82.67   85.07   84.53   86.40   82.67   84.00   82.67   82.67   85.87   85.87   83.73   84.00  100.00   84.80   81.60   86.67   84.53   81.60   82.93   83.20   82.67   82.93   82.13   82.67   83.73   83.47   82.93   84.00   82.93   82.40   82.93   84.53   83.47   85.60   83.20   84.80   84.80   85.07   84.80   85.07   84.80   82.13   82.40   84.53   82.13   83.20   85.33   83.47   82.40   81.87   85.07   83.47   84.80   85.60   82.93   84.80   84.53   86.13   84.80   85.33   85.07   82.40   85.87   84.53   85.07   85.07   82.40   84.27   83.47   88.27   88.53
    55: Sequence37    78.40   77.07   76.00   77.07   78.67   80.00   77.07   77.87   76.27   77.87   82.40   80.27   84.27   84.00   83.73   83.73   85.87   84.00   82.93   84.00   83.47   81.60   81.07   83.20   82.13   82.67   82.93   85.60   84.80   82.93   81.87   83.20   84.27   80.53   83.73   82.93   84.53   81.87   83.20   83.20   83.20   82.67   84.53   83.47   83.47   81.07   84.80   85.07   84.00   84.00   85.07   84.00   84.53   84.80  100.00   86.13   86.93   86.40   85.07   84.27   81.87   84.80   85.87   85.33   85.87   84.53   83.20   84.80   84.80   80.53   80.53   82.67   84.00   82.67   83.20   81.87   84.80   81.33   83.73   85.07   84.53   83.47   81.60   83.20   82.13   82.40   81.60   84.00   84.53   82.93   86.93   84.00   84.80   82.13   83.47   86.13   83.20   85.60   82.67   85.60   84.53   84.53   82.67   84.00   84.53   82.40   84.53   82.93   85.07   83.47   88.53   87.47
    56: Sequence93    77.33   78.93   76.00   76.53   79.47   77.33   77.87   78.40   80.00   77.60   80.53   81.60   83.73   80.00   82.67   81.60   81.87   82.13   82.93   83.73   83.20   81.33   82.40   82.67   81.60   80.80   81.33   82.93   81.07   80.80   81.60   84.27   84.53   82.13   82.13   81.33   81.33   82.13   81.07   85.60   84.27   83.47   83.73   82.93   80.80   82.13   84.00   83.20   81.33   82.67   82.93   85.60   82.67   81.60   86.13  100.00   85.87   83.73   87.20   82.40   82.40   82.67   82.40   86.13   85.33   84.27   83.20   83.73   81.07   83.73   81.33   81.07   82.40   80.53   81.60   81.87   81.87   85.07   82.67   81.60   84.27   81.87   81.87   84.53   82.67   84.27   81.07   84.53   84.80   82.93   83.47   83.73   80.80   80.27   81.60   86.40   81.33   82.93   81.60   87.20   83.47   81.87   83.20   82.93   83.47   81.60   85.87   82.40   82.67   82.40   86.40   85.60
    57: Sequence14    76.53   77.07   73.60   77.87   79.73   77.07   77.33   75.47   78.13   77.60   84.00   83.73   84.53   81.87   84.53   86.13   83.73   84.80   85.07   84.53   83.20   84.27   82.93   83.20   85.33   83.47   83.47   85.07   84.00   84.00   82.40   82.93   84.27   82.13   81.60   82.67   84.80   82.40   84.80   83.47   85.07   84.27   86.40   85.60   82.67   84.00   85.87   86.40   81.87   85.07   85.87   84.80   81.33   86.67   86.93   85.87  100.00   84.53   88.00   83.20   82.93   83.73   84.80   83.47   84.27   84.53   83.47   84.00   85.87   83.47   82.67   81.60   83.73   84.00   84.27   82.93   84.27   84.27   84.80   82.40   84.27   83.73   81.87   82.93   83.47   82.40   83.20   83.47   87.20   86.67   83.20   86.13   83.47   84.00   83.47   84.80   82.40   85.07   84.53   85.07   86.40   85.87   84.80   85.87   87.20   83.47   84.80   84.00   84.00   84.53   88.53   87.73
    58: Sequence8     76.53   77.33   75.47   78.13   78.67   78.93   76.00   78.40   77.87   79.47   82.40   79.73   84.80   82.93   84.27   85.33   82.93   84.80   83.73   82.13   84.53   82.13   82.40   82.67   84.80   85.60   83.47   82.93   85.07   86.40   81.33   82.67   81.87   82.13   81.60   84.27   84.53   82.13   84.80   83.20   83.20   84.53   82.67   84.27   80.53   84.53   85.87   82.67   83.20   84.00   85.60   85.33   86.40   84.53   86.40   83.73   84.53  100.00   87.73   82.93   83.47   83.73   83.73   87.47   84.80   85.33   82.93   81.87   84.53   81.60   81.60   83.47   83.47   84.80   81.87   82.93   83.20   82.13   82.13   84.53   83.47   83.20   82.40   83.47   83.73   82.67   84.53   84.27   83.47   85.60   84.00   81.33   83.73   84.80   83.47   86.40   85.33   85.33   81.87   85.33   85.07   83.20   84.27   84.53   85.60   85.87   84.80   84.53   84.27   85.07   89.07   87.73
    59: Sequence32    75.73   77.87   76.00   77.87   78.93   77.33   78.67   78.13   77.33   77.60   81.33   83.47   84.00   82.93   82.40   84.80   82.40   82.40   84.53   81.60   82.93   84.00   84.27   83.47   85.60   82.13   84.00   84.27   84.00   83.73   80.80   84.00   84.00   81.33   81.07   81.60   82.13   81.87   83.20   83.47   84.53   84.80   84.27   83.47   81.60   84.00   86.40   84.00   81.60   82.67   84.27   86.13   84.27   81.60   85.07   87.20   88.00   87.73  100.00   83.20   83.47   84.27   85.33   85.87   85.07   82.67   81.60   83.20   83.20   80.53   81.60   82.40   84.00   81.33   84.00   85.60   85.07   83.47   83.20   82.67   82.93   84.53   82.40   83.47   80.80   81.33   82.40   81.33   84.53   84.80   83.47   83.73   81.60   82.67   82.67   85.07   82.40   84.80   82.40   86.40   85.60   83.73   85.60   86.13   84.80   82.93   84.53   84.27   83.47   83.47   88.53   87.73
    60: Sequence100   76.53   77.87   74.67   77.33   80.80   78.13   76.27   79.47   74.13   77.60   84.27   81.60   82.93   81.60   80.00   82.93   82.13   80.53   83.20   82.67   82.40   83.73   80.53   84.53   82.93   81.60   81.60   83.73   84.00   81.87   80.27   81.33   82.93   80.80   81.07   81.33   82.93   81.60   82.67   82.67   82.40   84.53   82.67   80.00   81.60   82.93   82.67   83.73   80.27   82.40   83.47   81.07   81.60   82.93   84.27   82.40   83.20   82.93   83.20  100.00   82.40   81.33   84.53   83.47   81.07   82.13   80.80   82.40   81.07   82.40   81.07   85.07   83.47   81.60   82.40   81.07   82.67   82.40   81.33   84.00   83.47   81.33   84.00   81.87   81.60   80.80   82.40   79.20   81.07   80.27   83.73   83.20   84.53   81.87   83.20   83.20   82.40   81.60   83.47   81.33   85.33   81.87   83.20   82.40   81.87   82.67   83.47   85.60   84.27   85.33   85.87   85.60
    61: Sequence72    79.20   76.53   77.87   78.93   73.60   77.07   77.33   77.07   77.60   77.87   84.00   82.67   84.00   84.80   84.27   82.13   83.73   82.13   85.33   82.13   80.80   82.93   81.07   82.40   80.80   83.20   84.00   82.67   82.13   82.13   81.60   83.20   84.53   80.27   82.40   84.00   81.60   83.73   81.60   82.40   81.33   80.27   84.27   83.47   83.73   84.80   81.60   82.67   84.27   83.47   83.47   82.13   82.93   83.20   81.87   82.40   82.93   83.47   83.47   82.40  100.00   84.53   85.87   84.80   83.47   82.93   78.40   85.33   83.47   84.27   83.73   84.80   82.67   84.00   84.27   81.33   83.73   82.93   81.07   81.60   84.27   85.07   83.47   84.27   82.67   85.60   84.27   81.60   82.40   82.67   84.00   83.20   82.13   85.33   82.40   84.00   84.27   85.07   84.80   82.93   84.00   82.67   84.27   86.93   83.47   86.13   84.53   85.07   85.07   84.27   88.27   85.33
    62: Sequence90    77.07   77.87   78.93   77.87   77.87   78.67   76.27   77.60   78.93   78.93   82.67   84.53   84.00   84.00   85.07   83.73   84.80   82.93   84.53   82.40   82.93   80.53   83.20   82.67   81.07   81.33   82.13   83.73   83.73   81.33   82.67   82.93   84.80   80.27   84.00   83.47   81.33   80.53   80.00   81.87   83.20   83.47   84.53   85.60   83.20   80.80   84.27   82.13   83.20   84.00   84.80   83.73   81.87   82.67   84.80   82.67   83.73   83.73   84.27   81.33   84.53  100.00   82.13   84.53   86.13   84.27   81.87   81.33   82.67   82.40   82.93   83.73   83.73   82.13   81.60   82.40   83.20   80.53   82.13   82.93   84.00   82.13   85.07   81.07   81.33   82.93   83.20   83.47   84.27   83.20   84.53   81.87   81.07   84.53   84.27   84.00   84.53   82.93   82.40   84.27   81.07   81.07   82.13   84.27   86.93   82.13   82.67   81.87   81.87   84.00   86.40   85.33
    63: Sequence83    76.27   77.60   77.33   76.80   78.40   79.20   78.40   79.73   77.33   76.53   82.40   81.33   83.20   83.20   82.93   82.40   84.00   80.53   82.40   83.20   80.27   84.27   81.33   84.00   81.87   83.47   84.27   84.80   84.00   84.80   80.00   80.53   81.60   81.33   80.00   80.80   82.40   81.07   83.73   82.40   80.53   83.73   82.93   83.20   82.13   85.33   82.67   82.40   82.13   83.73   83.47   83.20   84.53   82.93   85.87   82.40   84.80   83.73   85.33   84.53   85.87   82.13  100.00   83.47   86.13   81.60   80.00   81.60   84.53   80.53   83.20   82.93   82.93   81.87   82.13   82.93   83.47   81.33   81.87   81.60   80.00   82.67   82.40   82.67   80.80   83.47   82.93   81.87   82.67   80.80   84.80   81.07   81.60   81.33   83.20   83.73   82.67   82.93   82.40   85.87   82.67   82.67   83.20   84.27   83.20   81.33   84.00   83.73   82.67   82.13   86.40   85.07
    64: Sequence36    79.47   76.00   74.93   76.27   77.60   80.00   77.60   77.07   80.27   76.27   82.13   83.73   85.33   84.00   84.80   85.60   84.00   82.67   83.73   83.73   85.07   82.13   82.93   82.40   83.73   83.47   81.60   84.00   86.40   82.93   81.87   84.27   81.60   82.13   83.20   82.67   84.00   83.47   82.93   84.00   84.80   84.27   83.73   86.13   80.27   83.73   84.00   83.73   82.40   83.47   84.27   84.80   85.33   82.13   85.33   86.13   83.47   87.47   85.87   83.47   84.80   84.53   83.47  100.00   87.73   84.00   82.93   84.00   83.47   82.67   81.87   81.33   82.40   80.80   81.60   83.47   83.47   82.40   82.67   83.20   82.40   82.93   80.80   82.67   83.20   84.00   82.67   82.13   82.93   85.07   87.20   83.47   82.67   84.53   84.27   86.93   85.33   84.00   82.67   85.33   84.00   82.40   83.73   84.27   87.20   83.73   85.07   83.20   85.33   85.33   88.53   88.27
    65: Sequence56    79.20   78.93   78.13   73.33   77.87   79.20   74.40   78.13   77.33   76.80   81.33   82.93   82.93   84.53   85.60   84.53   85.87   83.20   82.93   84.27   84.80   81.87   85.60   83.73   82.40   82.93   81.87   85.60   84.80   83.73   83.20   85.07   82.93   82.67   80.53   83.47   81.60   81.33   80.00   82.40   82.40   83.73   82.67   85.60   80.53   82.67   82.13   85.07   84.27   85.07   83.47   84.27   84.53   82.67   85.87   85.33   84.27   84.80   85.07   81.07   83.47   86.13   86.13   87.73  100.00   85.33   83.73   83.47   83.73   83.47   82.40   80.53   84.53   81.87   82.93   82.13   82.13   83.47   83.47   81.87   81.60   80.00   81.60   82.67   82.93   86.67   85.33   84.00   86.67   85.33   85.33   81.60   82.40   84.27   85.87   86.40   84.00   85.07   83.20   86.40   82.13   81.60   84.27   84.27   86.13   84.27   84.80   83.20   85.60   85.87   87.73   88.00
    66: Sequence89    76.53   77.60   76.53   77.87   75.73   77.07   75.47   77.33   77.33   77.33   82.93   82.93   83.73   81.33   83.47   84.53   82.40   82.67   82.67   83.47   84.80   79.20   83.47   83.73   83.73   82.67   83.20   84.53   82.40   84.27   82.93   83.73   84.53   81.87   82.67   83.47   82.93   82.93   81.33   80.80   82.67   82.13   84.27   84.00   82.67   83.47   83.20   81.07   82.13   85.07   84.80   83.20   84.80   83.73   84.53   84.27   84.53   85.33   82.67   82.13   82.93   84.27   81.60   84.00   85.33  100.00   84.27   82.13   85.87   82.67   82.67   82.93   82.93   83.47   83.47   80.80   82.67   81.60   81.60   84.00   85.33   83.20   83.73   83.20   85.07   85.33   83.20   84.00   83.73   83.20   84.27   83.73   82.67   85.07   84.00   84.53   85.33   83.73   85.07   84.27   82.67   84.00   81.87   85.60   83.47   84.53   84.53   84.27   84.00   85.87   86.67   88.00
    67: Sequence79    79.20   78.40   78.67   79.73   78.13   77.07   74.67   77.87   80.53   80.53   82.13   82.13   83.47   84.00   81.60   82.67   81.07   81.60   79.47   80.27   81.60   82.40   83.73   81.33   80.80   84.27   83.73   84.80   84.27   83.20   82.40   82.40   81.33   85.33   84.53   84.00   83.20   84.27   82.93   82.93   82.67   84.53   82.13   82.93   82.40   80.80   80.80   83.20   82.67   83.20   84.00   83.73   85.07   83.47   83.20   83.20   83.47   82.93   81.60   80.80   78.40   81.87   80.00   82.93   83.73   84.27  100.00   79.47   85.33   81.33   81.60   81.87   82.93   84.80   84.53   84.27   82.67   82.67   82.40   84.80   83.73   82.13   84.53   81.87   82.93   83.47   81.87   84.27   84.53   83.47   81.60   84.53   84.53   82.13   86.13   83.73   83.47   85.07   84.53   84.27   86.13   86.13   84.53   82.13   82.93   81.87   84.27   84.00   83.73   84.27   86.40   86.13
    68: Sequence95    78.13   78.40   76.27   74.67   77.07   76.80   76.80   78.40   76.00   78.40   83.20   84.53   84.80   84.00   83.20   81.87   80.27   80.80   81.07   81.60   80.53   81.60   81.33   80.80   83.20   77.87   83.73   81.87   80.80   82.13   82.13   83.20   84.80   81.07   80.80   81.33   82.40   81.87   82.67   83.47   84.27   79.47   80.27   82.67   81.33   82.13   82.40   83.47   82.40   84.80   81.60   80.80   83.20   82.93   84.80   83.73   84.00   81.87   83.20   82.40   85.33   81.33   81.60   84.00   83.47   82.13   79.47  100.00   81.60   85.87   83.20   84.27   81.07   79.47   80.27   81.87   81.87   81.87   80.00   82.67   82.67   81.33   81.33   83.20   80.00   84.00   84.00   81.33   83.47   84.00   85.60   82.13   82.40   83.20   80.80   81.87   82.13   84.27   81.07   81.07   80.53   82.93   81.60   83.73   85.07   85.33   85.07   83.73   84.00   82.13   86.40   84.00
    69: Sequence54    76.80   78.67   76.80   79.20   75.73   77.07   73.33   78.13   78.13   78.93   85.60   82.40   83.47   84.00   84.80   85.33   83.20   82.40   81.60   82.93   81.07   82.40   84.27   82.67   84.27   83.20   84.27   84.27   84.00   85.33   82.13   83.73   82.40   83.73   83.20   85.33   84.53   85.60   82.93   84.27   83.73   83.20   84.27   82.40   84.27   84.27   83.47   85.60   82.67   85.33   82.40   82.93   84.27   84.00   84.80   81.07   85.87   84.53   83.20   81.07   83.47   82.67   84.53   83.47   83.73   85.87   85.33   81.60  100.00   84.27   86.67   84.00   84.00   83.47   84.00   83.73   84.27   84.00   85.87   84.27   84.00   82.67   82.13   82.67   84.27   84.80   84.80   84.27   85.33   87.20   83.20   86.13   82.67   84.00   84.53   84.27   84.80   85.07   82.13   85.33   82.93   85.87   82.13   85.33   83.73   82.93   83.73   85.87   83.20   85.33   88.53   87.47
    70: Sequence21    78.40   78.40   77.60   73.87   79.47   77.07   77.60   80.80   76.80   78.13   84.80   82.93   82.93   82.67   82.93   84.53   82.93   81.60   84.27   85.87   82.67   82.40   82.93   85.33   84.27   82.40   84.80   84.53   82.40   86.13   82.67   83.73   82.13   84.80   83.73   83.20   82.13   82.93   82.93   85.60   82.40   84.00   84.00   82.67   82.13   85.07   80.53   82.93   83.47   84.53   84.27   83.73   82.40   82.93   80.53   83.73   83.47   81.60   80.53   82.40   84.27   82.40   80.53   82.67   83.47   82.67   81.33   85.87   84.27  100.00   87.20   82.93   83.73   83.47   82.40   82.93   81.87   84.80   85.07   84.53   84.27   83.47   84.80   84.80   82.93   85.87   83.47   84.53   83.73   85.33   84.53   81.60   82.40   84.80   83.73   81.87   82.93   82.13   81.07   83.47   82.67   85.60   84.00   83.20   85.07   84.00   85.07   85.60   86.13   85.60   88.00   88.00
    71: Sequence64    77.87   80.53   77.60   74.93   77.87   77.33   76.53   81.07   77.87   76.80   83.47   84.00   84.00   84.27   84.80   85.07   84.27   81.33   82.13   82.67   79.73   81.07   83.73   84.00   83.20   82.13   83.73   83.73   84.27   82.67   82.40   82.67   85.60   82.67   82.67   80.80   82.67   83.73   81.07   85.33   82.13   82.93   83.20   82.67   81.60   85.33   80.00   83.47   81.87   84.53   81.07   80.53   81.87   82.40   80.53   81.33   82.67   81.60   81.60   81.07   83.73   82.93   83.20   81.87   82.40   82.67   81.60   83.20   86.67   87.20  100.00   82.67   83.73   81.07   84.00   80.80   81.33   83.47   82.93   82.67   83.73   81.60   83.73   83.73   84.53   86.67   85.33   84.53   83.20   84.80   82.67   82.13   82.40   83.20   85.07   82.67   83.20   83.20   83.47   82.67   81.07   84.27   82.67   85.87   85.33   81.60   85.33   82.40   83.47   84.53   87.20   86.13
    72: Sequence88    75.73   79.20   78.67   77.87   77.07   78.93   77.87   79.73   77.07   80.80   82.67   81.60   81.87   82.93   82.93   81.33   80.80   79.73   79.47   80.80   80.80   80.80   81.87   82.40   82.67   81.60   84.80   81.87   84.00   83.20   80.27   80.80   81.60   80.53   82.93   82.40   82.13   82.67   81.07   81.60   83.20   82.40   79.47   79.73   80.80   82.40   82.67   82.40   83.73   83.73   82.40   80.27   82.40   82.93   82.67   81.07   81.60   83.47   82.40   85.07   84.80   83.73   82.93   81.33   80.53   82.93   81.87   84.27   84.00   82.93   82.67  100.00   83.73   80.80   83.20   83.20   85.60   79.47   82.13   83.73   84.00   82.93   83.47   83.20   83.47   82.13   82.40   80.80   81.07   81.07   83.20   83.20   81.60   84.53   83.47   82.67   84.27   82.40   83.47   82.67   85.33   84.53   81.60   84.00   83.73   84.27   84.80   84.00   81.87   85.87   86.13   85.07
    73: Sequence78    80.00   79.20   79.47   76.53   79.47   77.07   77.87   80.27   75.47   75.47   83.20   82.67   82.67   84.53   85.33   82.40   84.27   82.67   80.53   82.67   82.93   80.53   82.67   85.07   83.73   80.53   83.20   83.47   85.60   84.27   81.07   81.87   82.40   82.67   81.60   82.40   81.87   82.93   80.80   81.33   82.40   82.40   82.40   83.20   82.67   83.20   82.93   84.53   84.80   82.93   81.60   84.27   82.67   84.53   84.00   82.40   83.73   83.47   84.00   83.47   82.67   83.73   82.93   82.40   84.53   82.93   82.93   81.07   84.00   83.73   83.73   83.73  100.00   84.27   86.67   83.20   84.27   85.07   83.73   83.47   85.33   84.80   82.13   84.80   83.73   82.67   82.93   82.93   83.73   84.00   84.80   83.73   81.60   83.20   86.67   86.13   84.80   85.33   85.33   83.20   83.73   83.20   83.20   85.33   81.87   82.13   83.73   80.80   82.40   83.47   86.40   87.47
    74: Sequence82    78.40   80.00   76.53   78.13   78.40   77.60   77.60   77.07   77.60   77.33   81.60   80.53   81.87   83.20   80.27   82.13   81.60   83.73   82.13   81.87   80.80   81.87   82.93   83.73   83.20   83.73   83.20   84.00   82.40   83.47   81.33   80.80   80.80   81.87   82.67   84.80   81.87   83.47   82.93   81.33   82.93   82.13   81.60   81.60   83.20   83.73   82.40   82.13   83.20   82.40   82.13   84.53   83.20   83.47   82.67   80.53   84.00   84.80   81.33   81.60   84.00   82.13   81.87   80.80   81.87   83.47   84.80   79.47   83.47   83.47   81.07   80.80   84.27  100.00   86.13   82.93   84.00   84.53   83.20   84.80   82.93   82.40   84.00   83.47   82.93   83.47   85.60   83.73   82.13   83.47   80.27   85.33   84.27   81.60   84.00   81.87   82.93   83.47   83.73   81.33   85.60   84.80   85.33   82.93   81.60   83.73   85.33   85.87   84.00   82.93   86.67   86.67
    75: Sequence46    78.93   78.40   76.00   78.93   78.67   77.33   75.73   76.80   79.47   76.27   84.27   82.67   82.67   84.00   83.73   84.00   84.00   82.93   81.33   84.53   84.00   82.40   85.33   82.93   80.27   81.87   82.67   84.27   84.80   82.67   81.87   81.87   82.93   81.60   82.40   83.47   84.80   82.93   80.00   82.40   83.20   82.13   83.73   83.47   83.73   83.47   82.40   83.47   84.27   82.67   83.47   83.47   82.13   85.60   83.20   81.60   84.27   81.87   84.00   82.40   84.27   81.60   82.13   81.60   82.93   83.47   84.53   80.27   84.00   82.40   84.00   83.20   86.67   86.13  100.00   84.80   87.20   84.27   84.27   84.27   86.40   84.53   84.27   84.53   84.27   84.53   84.53   82.67   83.20   82.40   80.80   85.87   85.07   82.67   85.60   84.00   83.20   84.80   86.93   83.73   84.80   85.60   84.27   85.33   81.60   82.67   85.07   84.00   84.80   84.53   88.00   87.73
    76: Sequence34    75.73   79.20   79.47   76.00   78.67   77.07   79.47   78.67   79.20   75.73   84.00   83.20   82.13   83.20   81.87   82.13   82.67   82.67   81.60   80.80   80.80   82.67   84.80   81.60   82.67   84.53   82.13   82.40   85.87   82.67   82.13   82.93   81.33   82.67   83.20   80.80   81.60   82.40   84.27   84.53   84.53   82.40   83.47   82.93   84.80   82.67   82.13   84.80   82.93   83.47   82.93   84.00   83.20   83.20   81.87   81.87   82.93   82.93   85.60   81.07   81.33   82.40   82.93   83.47   82.13   80.80   84.27   81.87   83.73   82.93   80.80   83.20   83.20   82.93   84.80  100.00   86.93   82.67   85.60   85.07   83.20   83.73   85.33   83.20   80.53   82.67   82.93   84.00   84.27   83.73   84.27   84.00   82.13   84.00   83.47   81.07   82.93   82.67   83.20   84.80   86.93   85.60   84.27   85.07   82.67   80.00   84.27   84.53   83.47   82.93   88.00   86.13
    77: Sequence65    78.13   79.20   78.13   77.33   78.93   78.67   77.33   78.67   76.80   77.33   80.53   82.67   82.93   85.60   81.87   83.73   83.20   81.60   81.33   81.33   80.53   82.13   84.00   84.00   79.73   83.47   81.07   83.20   84.00   82.13   83.47   84.80   83.47   83.47   83.47   82.93   85.60   85.87   84.00   83.73   84.80   80.27   83.73   82.40   85.07   83.47   84.53   82.40   83.20   83.20   84.80   84.00   84.27   84.80   84.80   81.87   84.27   83.20   85.07   82.67   83.73   83.20   83.47   83.47   82.13   82.67   82.67   81.87   84.27   81.87   81.33   85.60   84.27   84.00   87.20   86.93  100.00   82.93   86.13   85.33   85.33   85.87   83.20   85.60   82.40   82.67   82.93   84.00   83.47   82.93   83.47   86.93   81.60   82.93   82.93   83.47   82.93   85.87   85.60   85.33   85.33   85.87   83.47   85.60   84.00   81.60   86.40   84.53   82.93   83.73   88.27   86.13
    78: Sequence50    80.27   78.67   76.53   77.33   78.93   77.33   77.07   79.47   75.20   74.67   81.33   81.07   81.87   81.60   83.20   83.47   83.73   82.93   82.67   81.60   80.80   83.20   85.33   86.13   83.20   82.13   82.93   85.07   82.67   82.67   81.87   84.80   83.20   85.07   83.47   81.60   82.93   83.20   81.07   83.73   82.67   82.40   85.07   82.40   82.67   83.20   83.47   82.93   82.67   81.87   81.07   84.53   81.60   84.80   81.33   85.07   84.27   82.13   83.47   82.40   82.93   80.53   81.33   82.40   83.47   81.60   82.67   81.87   84.00   84.80   83.47   79.47   85.07   84.53   84.27   82.67   82.93  100.00   86.67   82.67   85.07   85.33   82.40   86.13   84.00   84.27   85.60   83.47   86.13   84.00   81.33   84.80   82.13   82.67   85.60   81.07   80.27   84.53   83.20   84.53   85.33   85.87   85.07   84.80   82.67   81.33   83.73   83.20   84.53   84.27   87.20   86.67
    79: Sequence68    78.93   78.93   77.33   76.00   79.73   74.93   76.00   78.13   74.93   74.93   81.60   80.80   81.60   84.00   82.67   84.27   84.80   81.60   83.20   84.27   80.53   83.47   84.53   84.27   84.80   81.87   81.33   84.53   83.47   85.07   83.47   84.00   81.60   82.67   85.87   83.73   85.33   83.47   81.87   85.60   81.60   84.00   84.80   81.87   84.80   84.53   85.60   82.67   84.53   84.00   82.40   84.53   82.93   85.07   83.73   82.67   84.80   82.13   83.20   81.33   81.07   82.13   81.87   82.67   83.47   81.60   82.40   80.00   85.87   85.07   82.93   82.13   83.73   83.20   84.27   85.60   86.13   86.67  100.00   84.80   86.13   85.33   81.87   83.47   82.67   82.93   84.27   84.80   83.20   83.20   83.20   85.33   81.33   82.40   83.20   82.93   83.73   83.20   83.73   83.47   87.47   86.93   85.33   84.80   84.53   82.13   82.67   85.33   83.47   84.27   88.27   87.73
    80: Sequence22    78.13   79.47   80.00   77.87   80.27   79.47   76.80   80.00   79.73   77.33   82.13   81.87   82.93   82.40   82.40   84.00   84.80   82.13   82.67   81.87   84.27   83.20   84.53   84.53   83.73   82.93   84.80   84.00   85.60   85.33   81.60   82.93   81.87   82.67   83.47   84.80   82.40   83.47   81.60   81.87   81.07   81.60   85.33   84.00   84.80   82.93   84.00   81.60   82.93   82.40   83.73   81.60   83.47   84.80   85.07   81.60   82.40   84.53   82.67   84.00   81.60   82.93   81.60   83.20   81.87   84.00   84.80   82.67   84.27   84.53   82.67   83.73   83.47   84.80   84.27   85.07   85.33   82.67   84.80  100.00   86.40   84.00   84.53   85.87   82.13   84.27   85.33   86.40   82.67   83.20   84.00   84.27   83.20   81.60   83.20   82.13   81.87   82.93   82.93   83.20   85.33   85.60   83.20   85.33   83.20   81.60   83.20   86.13   84.00   84.27   87.47   86.13
    81: Sequence10    77.60   79.20   78.93   80.00   78.93   74.93   77.07   78.67   77.33   75.73   83.20   83.47   85.07   83.73   85.07   84.00   83.47   82.93   84.53   82.40   83.73   82.67   84.00   83.20   84.53   82.67   85.07   83.47   83.73   81.87   81.60   82.40   85.07   82.93   84.27   83.20   82.40   82.13   80.00   83.47   82.67   81.33   85.60   82.13   85.33   81.07   82.13   80.80   85.07   83.20   84.53   81.87   83.20   85.07   84.53   84.27   84.27   83.47   82.93   83.47   84.27   84.00   80.00   82.40   81.60   85.33   83.73   82.67   84.00   84.27   83.73   84.00   85.33   82.93   86.40   83.20   85.33   85.07   86.13   86.40  100.00   87.47   82.93   84.27   83.20   82.67   84.27   84.53   80.80   82.40   84.80   84.53   83.73   82.93   85.07   81.60   84.00   85.07   84.53   82.93   85.07   85.60   83.20   85.07   83.47   84.53   84.00   83.47   83.73   84.27   88.53   87.20
    82: Sequence55    77.60   76.00   78.13   78.13   76.80   75.73   77.87   78.13   76.80   74.67   84.27   81.87   84.00   83.20   82.93   83.20   82.40   80.53   83.47   81.60   80.80   84.53   82.13   81.87   80.80   82.13   84.00   81.87   81.07   82.13   78.93   84.00   82.40   84.53   84.00   81.33   83.20   83.20   80.53   83.73   84.27   80.80   85.60   83.47   84.80   82.67   81.60   79.73   84.00   85.07   82.93   82.40   85.07   84.80   83.47   81.87   83.73   83.20   84.53   81.33   85.07   82.13   82.67   82.93   80.00   83.20   82.13   81.33   82.67   83.47   81.60   82.93   84.80   82.40   84.53   83.73   85.87   85.33   85.33   84.00   87.47  100.00   82.40   85.33   82.40   81.60   84.53   83.47   81.87   82.93   83.73   84.80   81.33   84.00   82.40   81.87   84.00   84.53   84.80   82.67   86.40   82.13   83.73   84.27   83.73   83.47   82.13   82.67   82.93   82.93   87.20   87.73
    83: Sequence45    78.40   78.13   79.47   80.00   77.87   77.07   77.60   77.87   79.20   77.60   84.53   84.27   82.67   82.40   81.87   81.87   82.13   81.60   84.00   84.00   81.87   86.67   82.67   83.47   83.47   85.60   82.67   83.20   83.20   84.00   84.53   81.87   81.60   80.53   84.80   84.00   81.07   81.87   82.13   81.87   80.00   82.67   84.53   84.80   85.33   82.93   83.47   81.60   82.67   85.07   84.27   81.33   85.07   82.13   81.60   81.87   81.87   82.40   82.40   84.00   83.47   85.07   82.40   80.80   81.60   83.73   84.53   81.33   82.13   84.80   83.73   83.47   82.13   84.00   84.27   85.33   83.20   82.40   81.87   84.53   82.93   82.40  100.00   84.27   85.07   86.13   82.67   82.40   82.40   83.20   83.73   82.40   84.53   83.73   83.47   82.67   84.00   83.73   86.40   83.20   82.93   84.80   86.40   85.87   84.00   85.60   83.20   85.33   85.87   85.33   87.20   86.67
    84: Sequence41    80.53   77.07   75.73   77.07   79.73   77.33   78.13   78.67   76.80   77.33   83.73   82.13   82.40   81.60   81.87   82.13   82.67   81.07   80.80   81.07   80.80   83.73   83.20   85.07   80.53   81.07   83.73   83.20   85.07   84.00   82.40   82.40   82.67   83.47   86.13   82.13   84.00   85.07   80.27   84.00   84.00   79.47   85.87   83.20   82.93   83.20   81.33   82.13   86.13   82.93   81.60   83.73   82.13   82.40   83.20   84.53   82.93   83.47   83.47   81.87   84.27   81.07   82.67   82.67   82.67   83.20   81.87   83.20   82.67   84.80   83.73   83.20   84.80   83.47   84.53   83.20   85.60   86.13   83.47   85.87   84.27   85.33   84.27  100.00   82.67   85.07   85.87   85.60   81.07   84.53   84.53   84.27   82.93   82.13   85.07   83.73   82.40   83.20   84.80   84.00   83.73   85.07   85.33   86.93   80.27   83.73   85.87   81.33   85.60   83.20   87.47   86.40
    85: Sequence67    76.80   76.53   77.07   78.93   75.20   76.53   76.27   78.93   78.67   77.87   83.20   81.33   85.07   83.20   85.33   84.80   83.20   84.53   84.27   85.33   84.27   84.00   83.20   82.40   83.20   85.07   83.20   82.93   82.93   81.33   85.87   83.47   82.67   80.80   82.13   84.80   85.07   84.27   83.73   83.47   81.60   81.60   81.33   84.00   82.67   85.07   82.40   82.67   82.67   81.60   82.93   82.13   83.47   84.53   82.13   82.67   83.47   83.73   80.80   81.60   82.67   81.33   80.80   83.20   82.93   85.07   82.93   80.00   84.27   82.93   84.53   83.47   83.73   82.93   84.27   80.53   82.40   84.00   82.67   82.13   83.20   82.40   85.07   82.67  100.00   84.00   84.27   83.20   82.40   81.07   83.20   85.60   84.53   85.07   83.73   85.07   84.00   85.60   83.47   83.47   84.80   83.20   85.60   85.60   84.80   86.93   85.33   84.53   85.33   87.73   87.73   87.73
    86: Sequence75    78.40   78.67   77.87   77.33   77.60   77.87   76.00   78.13   81.60   77.33   82.40   84.53   82.67   82.93   85.60   82.93   84.00   82.67   81.87   85.07   81.60   83.20   86.13   84.27   83.20   82.67   82.40   84.80   83.20   85.60   84.27   84.80   84.00   85.33   84.00   82.40   82.13   83.73   80.53   84.00   82.13   80.27   83.47   84.27   82.93   83.47   80.80   84.00   82.40   83.20   83.20   81.87   82.40   82.13   82.40   84.27   82.40   82.67   81.33   80.80   85.60   82.93   83.47   84.00   86.67   85.33   83.47   84.00   84.80   85.87   86.67   82.13   82.67   83.47   84.53   82.67   82.67   84.27   82.93   84.27   82.67   81.60   86.13   85.07   84.00  100.00   85.07   86.67   84.80   86.40   85.33   85.07   84.00   84.00   84.27   84.80   84.27   84.00   85.87   83.20   83.20   84.80   84.80   86.93   84.80   84.27   87.73   85.60   87.73   85.07   87.73   87.73
    87: Sequence33    78.13   79.47   77.87   79.20   79.73   77.87   76.00   78.40   76.00   76.53   82.67   82.13   85.33   83.20   84.00   84.00   82.13   82.13   81.87   82.13   81.87   83.20   87.73   84.53   82.67   81.33   81.87   85.60   82.40   83.20   82.93   82.67   84.53   83.73   83.73   82.13   83.73   82.40   80.53   83.73   83.20   81.33   82.93   84.27   83.73   82.93   83.20   83.20   84.00   81.87   82.13   81.87   84.53   83.20   81.60   81.07   83.20   84.53   82.40   82.40   84.27   83.20   82.93   82.67   85.33   83.20   81.87   84.00   84.80   83.47   85.33   82.40   82.93   85.60   84.53   82.93   82.93   85.60   84.27   85.33   84.27   84.53   82.67   85.87   84.27   85.07  100.00   85.60   82.93   85.87   81.87   84.00   83.73   83.47   86.13   81.33   83.20   85.07   83.47   81.33   83.47   84.53   84.00   84.80   84.00   85.60   85.60   84.27   85.33   84.80   88.00   85.87
    88: Sequence29    78.13   79.20   78.13   78.40   79.73   79.20   75.73   79.47   79.47   77.33   82.67   79.73   81.60   80.80   83.20   84.00   83.20   80.80   81.87   82.93   80.80   81.87   84.00   84.27   81.60   81.07   82.93   82.93   83.73   83.47   82.67   83.47   82.93   86.13   84.00   83.20   85.07   85.60   80.80   85.07   82.93   80.00   84.80   82.93   84.53   81.60   82.40   82.93   85.60   84.53   84.27   83.20   84.27   85.33   84.00   84.53   83.47   84.27   81.33   79.20   81.60   83.47   81.87   82.13   84.00   84.00   84.27   81.33   84.27   84.53   84.53   80.80   82.93   83.73   82.67   84.00   84.00   83.47   84.80   86.40   84.53   83.47   82.40   85.60   83.20   86.67   85.60  100.00   85.07   85.87   84.80   86.67   82.93   84.80   85.07   83.20   85.07   82.93   84.27   83.73   82.13   85.33   81.33   83.73   85.07   82.13   85.87   84.27   83.20   82.93   88.00   87.20
    89: Sequence31    77.60   77.87   77.07   76.00   78.93   77.87   74.67   77.87   79.20   77.60   83.47   82.67   83.47   83.47   85.87   84.00   86.13   84.80   82.67   83.47   82.93   83.20   84.80   85.33   81.87   83.20   81.33   83.47   82.40   83.20   83.47   84.80   84.80   83.47   80.00   83.20   81.33   81.33   82.93   81.60   84.27   83.47   85.33   84.00   81.87   83.47   84.27   85.33   81.87   84.00   85.07   85.87   83.73   83.47   84.53   84.80   87.20   83.47   84.53   81.07   82.40   84.27   82.67   82.93   86.67   83.73   84.53   83.47   85.33   83.73   83.20   81.07   83.73   82.13   83.20   84.27   83.47   86.13   83.20   82.67   80.80   81.87   82.40   81.07   82.40   84.80   82.93   85.07  100.00   86.67   84.80   84.53   81.60   83.73   84.00   86.13   82.93   84.80   85.07   85.07   84.00   84.00   84.53   83.73   85.87   83.47   86.93   84.53   84.80   86.13   88.27   88.00
    90: Sequence7     78.40   77.60   75.73   77.87   79.20   77.60   75.47   77.33   80.00   77.60   84.00   82.93   81.33   83.47   83.47   85.87   84.00   83.73   83.47   82.67   81.87   81.60   85.60   82.40   82.93   82.13   81.60   84.53   84.00   83.73   84.27   85.60   84.53   84.27   81.60   83.20   80.80   82.13   81.87   83.47   86.40   83.20   84.27   83.73   83.20   82.13   84.00   85.07   82.13   84.80   85.07   84.27   82.93   82.40   82.93   82.93   86.67   85.60   84.80   80.27   82.67   83.20   80.80   85.07   85.33   83.20   83.47   84.00   87.20   85.33   84.80   81.07   84.00   83.47   82.40   83.73   82.93   84.00   83.20   83.20   82.40   82.93   83.20   84.53   81.07   86.40   85.87   85.87   86.67  100.00   82.93   83.73   83.20   84.27   82.40   86.67   84.53   83.20   85.07   85.87   83.20   84.53   83.47   84.27   85.07   83.47   85.07   84.80   84.27   84.27   88.53   88.27
    91: Sequence51    78.67   76.27   76.53   74.13   78.13   77.07   78.13   79.73   78.40   76.27   84.53   83.47   84.80   84.27   85.33   83.20   83.73   80.53   81.87   83.73   84.00   82.40   83.47   83.73   84.27   84.00   83.47   85.07   85.33   83.73   80.80   82.93   83.73   82.13   84.00   81.60   84.80   82.40   82.40   84.53   85.07   80.80   82.40   84.80   83.20   82.40   83.47   83.47   85.60   85.07   83.20   83.73   85.60   81.87   86.93   83.47   83.20   84.00   83.47   83.73   84.00   84.53   84.80   87.20   85.33   84.27   81.60   85.60   83.20   84.53   82.67   83.20   84.80   80.27   80.80   84.27   83.47   81.33   83.20   84.00   84.80   83.73   83.73   84.53   83.20   85.33   81.87   84.80   84.80   82.93  100.00   84.00   84.27   86.67   84.80   84.80   87.20   82.93   85.07   85.07   84.27   84.53   83.47   85.60   85.87   84.53   85.60   83.20   85.07   85.07   88.27   88.53
    92: Sequence43    75.73   76.53   74.40   78.40   77.87   77.07   78.13   77.60   78.93   77.07   83.47   83.47   82.93   83.47   83.47   82.67   82.40   82.13   81.60   82.40   80.80   84.27   83.47   82.13   81.60   82.67   82.67   82.40   82.67   83.47   82.13   84.00   82.67   83.47   83.20   83.47   84.27   86.40   81.60   85.33   84.00   80.53   84.27   81.87   85.07   84.00   85.07   82.40   81.60   82.93   82.67   84.00   82.13   85.07   84.00   83.73   86.13   81.33   83.73   83.20   83.20   81.87   81.07   83.47   81.60   83.73   84.53   82.13   86.13   81.60   82.13   83.20   83.73   85.33   85.87   84.00   86.93   84.80   85.33   84.27   84.53   84.80   82.40   84.27   85.60   85.07   84.00   86.67   84.53   83.73   84.00  100.00   82.93   83.47   85.87   84.27   84.80   85.87   85.07   85.60   84.53   86.13   84.80   84.53   83.47   83.20   85.60   86.13   82.93   85.60   88.53   88.80
    93: Sequence27    78.93   77.07   75.47   78.40   78.67   77.60   76.00   76.27   78.93   77.87   85.60   83.20   84.80   84.00   82.40   85.33   84.53   86.93   83.20   83.20   84.00   82.93   83.20   81.33   82.40   85.87   82.13   84.53   85.33   84.53   86.13   83.20   83.20   83.47   83.73   85.07   86.13   84.27   85.60   82.67   81.60   81.60   81.07   83.47   84.00   81.07   82.67   84.27   81.87   84.53   84.27   82.67   81.60   83.47   84.80   80.80   83.47   83.73   81.60   84.53   82.13   81.07   81.60   82.67   82.40   82.67   84.53   82.40   82.67   82.40   82.40   81.60   81.60   84.27   85.07   82.13   81.60   82.13   81.33   83.20   83.73   81.33   84.53   82.93   84.53   84.00   83.73   82.93   81.60   83.20   84.27   82.93  100.00   84.53   85.07   82.40   82.40   84.80   84.00   84.27   86.13   83.20   86.13   83.73   83.47   86.93   86.13   86.40   86.67   83.73   88.80   86.93
    94: Sequence25    79.20   77.07   76.53   75.73   76.53   78.40   78.13   77.60   77.33   78.93   85.87   84.53   83.73   84.00   85.60   82.93   84.00   83.73   85.07   84.80   81.87   83.73   82.93   84.00   84.27   86.13   82.13   84.00   84.80   84.00   82.67   84.00   83.20   84.27   83.20   84.00   85.87   84.80   85.33   82.93   82.40   82.40   81.87   84.00   83.47   82.93   81.87   82.93   84.27   85.87   83.73   82.67   83.73   84.80   82.13   80.27   84.00   84.80   82.67   81.87   85.33   84.53   81.33   84.53   84.27   85.07   82.13   83.20   84.00   84.80   83.20   84.53   83.20   81.60   82.67   84.00   82.93   82.67   82.40   81.60   82.93   84.00   83.73   82.13   85.07   84.00   83.47   84.80   83.73   84.27   86.67   83.47   84.53  100.00   85.60   84.53   86.40   84.80   85.60   82.67   82.93   84.00   83.47   84.53   87.20   86.40   83.47   84.53   82.93   84.80   88.80   87.73
    95: Sequence9     80.00   78.13   77.33   79.20   80.00   79.73   76.80   78.67   76.00   77.07   82.40   83.47   83.47   83.47   85.33   86.13   85.33   82.67   83.73   84.80   81.07   82.93   84.53   85.33   83.73   84.80   85.07   84.80   86.93   85.87   83.20   82.40   83.20   85.07   85.60   84.53   84.53   85.33   82.40   84.27   81.87   83.20   82.67   83.20   83.20   84.00   82.40   84.27   83.73   85.07   82.67   84.27   84.00   85.60   83.47   81.60   83.47   83.47   82.67   83.20   82.40   84.27   83.20   84.27   85.87   84.00   86.13   80.80   84.53   83.73   85.07   83.47   86.67   84.00   85.60   83.47   82.93   85.60   83.20   83.20   85.07   82.40   83.47   85.07   83.73   84.27   86.13   85.07   84.00   82.40   84.80   85.87   85.07   85.60  100.00   84.27   84.27   86.93   84.53   85.07   84.00   85.07   84.00   84.80   82.93   85.07   83.47   82.40   84.27   85.60   88.27   88.80
    96: Sequence63    77.87   78.40   74.93   76.53   78.13   78.67   77.07   78.13   78.93   76.27   82.13   81.33   83.20   84.80   86.67   83.73   84.80   82.67   81.60   86.40   84.53   82.93   81.87   84.80   82.67   83.73   80.53   84.00   83.73   82.93   84.00   82.67   83.20   83.73   82.93   84.27   84.00   84.00   81.87   82.40   83.47   84.00   84.80   84.00   81.33   85.07   84.27   83.47   82.40   83.20   84.53   86.13   83.73   82.93   86.13   86.40   84.80   86.40   85.07   83.20   84.00   84.00   83.73   86.93   86.40   84.53   83.73   81.87   84.27   81.87   82.67   82.67   86.13   81.87   84.00   81.07   83.47   81.07   82.93   82.13   81.60   81.87   82.67   83.73   85.07   84.80   81.33   83.20   86.13   86.67   84.80   84.27   82.40   84.53   84.27  100.00   86.40   83.47   84.53   85.60   84.27   82.13   84.00   85.87   85.60   84.53   83.73   82.93   83.73   86.67   88.27   88.80
    97: Sequence20    77.07   77.60   76.80   74.40   76.00   77.60   76.53   76.53   77.60   76.80   83.73   82.40   84.27   84.80   84.80   83.73   82.67   80.80   82.67   83.20   84.00   81.07   83.47   83.73   83.20   82.93   82.40   83.47   83.73   84.53   82.40   83.73   82.93   82.93   84.27   84.00   84.00   84.27   81.60   84.00   85.07   80.80   81.07   84.80   82.40   82.40   82.67   82.13   85.07   85.60   84.53   82.40   85.87   84.80   83.20   81.33   82.40   85.33   82.40   82.40   84.27   84.53   82.67   85.33   84.00   85.33   83.47   82.13   84.80   82.93   83.20   84.27   84.80   82.93   83.20   82.93   82.93   80.27   83.73   81.87   84.00   84.00   84.00   82.40   84.00   84.27   83.20   85.07   82.93   84.53   87.20   84.80   82.40   86.40   84.27   86.40  100.00   84.27   84.80   84.53   82.93   84.80   84.27   85.07   84.80   87.20   85.33   84.80   88.00   85.07   89.07   89.07
    98: Sequence18    80.27   80.53   78.13   78.67   77.07   77.07   76.53   76.80   76.53   77.33   82.40   83.73   84.00   83.47   85.87   83.47   84.00   84.80   84.80   82.13   82.67   84.53   83.73   81.87   84.27   82.67   83.20   83.47   83.47   82.67   84.27   84.27   81.87   82.13   82.13   81.07   84.80   84.00   83.20   82.13   83.20   81.60   83.73   82.93   82.93   82.93   81.60   85.60   84.00   82.93   83.20   83.47   84.00   84.53   85.60   82.93   85.07   85.33   84.80   81.60   85.07   82.93   82.93   84.00   85.07   83.73   85.07   84.27   85.07   82.13   83.20   82.40   85.33   83.47   84.80   82.67   85.87   84.53   83.20   82.93   85.07   84.53   83.73   83.20   85.60   84.00   85.07   82.93   84.80   83.20   82.93   85.87   84.80   84.80   86.93   83.47   84.27  100.00   86.67   84.80   85.60   84.80   87.73   85.33   83.73   86.93   85.60   84.00   85.07   85.33   88.80   86.13
    99: Sequence15    81.07   78.13   77.87   78.67   78.13   79.20   77.87   77.33   78.13   74.67   84.80   84.80   82.67   83.47   82.40   82.40   83.20   81.87   84.00   84.53   83.73   82.67   85.33   82.40   81.07   83.47   81.07   84.00   83.47   81.33   82.40   81.87   82.67   82.93   84.27   83.20   84.80   85.60   82.93   82.13   82.67   80.80   85.07   84.80   85.33   82.13   83.47   83.73   84.27   85.60   84.80   82.93   83.20   86.13   82.67   81.60   84.53   81.87   82.40   83.47   84.80   82.40   82.40   82.67   83.20   85.07   84.53   81.07   82.13   81.07   83.47   83.47   85.33   83.73   86.93   83.20   85.60   83.20   83.73   82.93   84.53   84.80   86.40   84.80   83.47   85.87   83.47   84.27   85.07   85.07   85.07   85.07   84.00   85.60   84.53   84.53   84.80   86.67  100.00   83.73   85.87   84.53   86.13   85.33   83.73   84.53   84.53   82.93   85.33   85.33   88.80   88.00
   100: Sequence12    78.40   76.00   75.73   76.27   77.87   79.47   79.47   77.87   78.93   77.07   84.53   85.07   84.00   85.33   82.93   85.87   85.87   81.87   83.47   83.73   84.53   81.60   83.20   83.20   80.80   84.53   84.27   84.53   85.87   86.67   83.47   82.93   81.60   81.87   82.13   83.73   83.73   84.00   81.87   85.60   83.47   84.53   85.33   84.00   82.93   84.53   84.00   84.53   84.27   85.60   86.67   85.60   84.80   84.80   85.60   87.20   85.07   85.33   86.40   81.33   82.93   84.27   85.87   85.33   86.40   84.27   84.27   81.07   85.33   83.47   82.67   82.67   83.20   81.33   83.73   84.80   85.33   84.53   83.47   83.20   82.93   82.67   83.20   84.00   83.47   83.20   81.33   83.73   85.07   85.87   85.07   85.60   84.27   82.67   85.07   85.60   84.53   84.80   83.73  100.00   85.33   85.07   83.73   85.60   85.33   83.47   86.40   83.47   85.60   84.00   89.07   88.00
   101: Sequence2     78.40   78.13   77.07   78.93   77.87   77.07   78.93   79.20   78.93   78.13   84.00   82.40   84.53   85.33   81.60   83.47   83.47   83.47   82.93   82.40   82.40   84.00   84.80   82.93   82.93   87.20   84.00   85.33   84.53   84.00   81.87   84.27   82.13   83.47   84.53   81.60   83.47   83.73   83.73   85.33   84.80   85.33   84.27   82.40   85.87   83.73   83.47   83.73   83.20   82.67   84.27   84.00   81.33   85.33   84.53   83.47   86.40   85.07   85.60   85.33   84.00   81.07   82.67   84.00   82.13   82.67   86.13   80.53   82.93   82.67   81.07   85.33   83.73   85.60   84.80   86.93   85.33   85.33   87.47   85.33   85.07   86.40   82.93   83.73   84.80   83.20   83.47   82.13   84.00   83.20   84.27   84.53   86.13   82.93   84.00   84.27   82.93   85.60   85.87   85.33  100.00   86.13   88.53   86.67   83.73   84.80   85.60   85.87   87.47   85.33   89.07   87.73
   102: Sequence19    79.47   77.07   76.27   77.60   78.93   76.27   76.80   80.00   76.00   76.27   84.27   83.47   82.40   82.13   81.87   82.93   82.40   81.33   81.60   82.93   83.73   82.13   84.53   84.27   82.93   84.27   83.20   85.33   83.73   84.80   82.13   83.47   82.67   83.20   85.33   83.73   85.33   84.80   83.20   84.80   82.40   84.27   84.80   82.93   85.60   83.20   81.87   84.00   85.33   84.53   84.27   85.60   85.07   85.07   84.53   81.87   85.87   83.20   83.73   81.87   82.67   81.07   82.67   82.40   81.60   84.00   86.13   82.93   85.87   85.60   84.27   84.53   83.20   84.80   85.60   85.60   85.87   85.87   86.93   85.60   85.60   82.13   84.80   85.07   83.20   84.80   84.53   85.33   84.00   84.53   84.53   86.13   83.20   84.00   85.07   82.13   84.80   84.80   84.53   85.07   86.13  100.00   83.20   85.07   84.80   83.20   85.60   85.33   85.07   85.60   89.07   88.80
   103: Sequence1     80.00   77.33   76.27   78.40   78.13   77.07   77.33   76.53   78.67   79.20   84.00   83.73   84.27   84.27   84.27   82.40   82.13   85.87   83.73   84.53   83.20   86.40   84.00   85.07   83.73   87.20   82.40   84.80   84.00   85.87   82.93   82.93   81.60   80.80   83.73   82.93   82.93   84.00   84.00   82.93   84.27   84.27   82.93   82.40   84.80   85.60   84.80   84.00   81.60   82.93   83.47   83.20   84.00   82.40   82.67   83.20   84.80   84.27   85.60   83.20   84.27   82.13   83.20   83.73   84.27   81.87   84.53   81.60   82.13   84.00   82.67   81.60   83.20   85.33   84.27   84.27   83.47   85.07   85.33   83.20   83.20   83.73   86.40   85.33   85.60   84.80   84.00   81.33   84.53   83.47   83.47   84.80   86.13   83.47   84.00   84.00   84.27   87.73   86.13   83.73   88.53   83.20  100.00   84.80   82.67   85.60   86.67   86.67   86.93   86.40   89.33   87.20
   104: Sequence11    77.33   76.53   77.07   76.80   77.33   77.87   76.53   76.80   78.67   78.13   84.00   83.47   85.07   85.87   86.40   82.93   84.53   83.20   83.47   84.27   85.87   83.20   83.47   83.73   85.33   83.20   85.07   84.27   86.13   83.47   82.40   84.80   85.07   81.87   83.47   82.93   85.60   84.27   82.40   83.47   85.33   81.07   85.60   85.87   85.33   85.07   83.47   82.93   85.87   84.27   85.07   82.93   83.73   85.87   84.00   82.93   85.87   84.53   86.13   82.40   86.93   84.27   84.27   84.27   84.27   85.60   82.13   83.73   85.33   83.20   85.87   84.00   85.33   82.93   85.33   85.07   85.60   84.80   84.80   85.33   85.07   84.27   85.87   86.93   85.60   86.93   84.80   83.73   83.73   84.27   85.60   84.53   83.73   84.53   84.80   85.87   85.07   85.33   85.33   85.60   86.67   85.07   84.80  100.00   83.47   84.80   86.40   83.47   86.13   85.87   89.60   88.53
   105: Sequence4     78.93   77.33   74.67   74.67   77.33   77.87   78.67   76.80   77.33   78.40   85.07   83.73   87.47   86.13   87.20   86.67   85.33   83.73   83.47   83.47   82.67   85.07   83.73   81.87   81.87   84.27   84.00   84.53   82.93   84.27   84.80   83.20   84.27   83.47   84.00   83.20   84.00   83.47   82.93   82.40   81.33   84.53   85.07   86.67   81.87   83.73   84.00   83.47   85.07   85.60   86.13   85.33   84.00   84.53   84.53   83.47   87.20   85.60   84.80   81.87   83.47   86.93   83.20   87.20   86.13   83.47   82.93   85.07   83.73   85.07   85.33   83.73   81.87   81.60   81.60   82.67   84.00   82.67   84.53   83.20   83.47   83.73   84.00   80.27   84.80   84.80   84.00   85.07   85.87   85.07   85.87   83.47   83.47   87.20   82.93   85.60   84.80   83.73   83.73   85.33   83.73   84.80   82.67   83.47  100.00   84.27   84.53   85.33   84.80   84.53   89.33   88.53
   106: Sequence39    77.60   77.07   76.27   77.87   76.27   76.27   76.27   76.00   78.93   76.27   85.33   83.73   83.47   82.93   83.20   85.87   83.47   83.73   86.40   85.33   84.00   84.53   83.20   82.93   85.33   84.53   84.00   84.53   82.13   83.47   85.07   84.27   81.60   83.20   84.80   86.40   84.00   82.93   84.53   82.93   83.20   82.40   82.13   84.27   84.80   81.87   85.07   84.27   83.20   83.47   84.27   82.67   84.27   85.07   82.40   81.60   83.47   85.87   82.93   82.67   86.13   82.13   81.33   83.73   84.27   84.53   81.87   85.33   82.93   84.00   81.60   84.27   82.13   83.73   82.67   80.00   81.60   81.33   82.13   81.60   84.53   83.47   85.60   83.73   86.93   84.27   85.60   82.13   83.47   83.47   84.53   83.20   86.93   86.40   85.07   84.53   87.20   86.93   84.53   83.47   84.80   83.20   85.60   84.80   84.27  100.00   85.60   85.87   86.93   86.93   88.80   89.60
   107: Sequence3     77.60   78.40   76.27   75.47   79.73   77.87   76.53   79.47   79.20   78.13   84.53   84.27   85.87   86.67   83.73   85.07   84.27   83.73   82.93   81.87   84.53   81.87   85.87   84.00   82.93   84.27   84.27   85.07   84.27   84.00   85.87   85.87   84.80   85.07   82.40   83.47   84.80   84.80   82.67   85.07   87.47   82.13   82.40   83.20   84.00   84.27   83.47   86.13   82.93   85.33   84.53   86.93   85.07   85.07   84.53   85.87   84.80   84.80   84.53   83.47   84.53   82.67   84.00   85.07   84.80   84.53   84.27   85.07   83.73   85.07   85.33   84.80   83.73   85.33   85.07   84.27   86.40   83.73   82.67   83.20   84.00   82.13   83.20   85.87   85.33   87.73   85.60   85.87   86.93   85.07   85.60   85.60   86.13   83.47   83.47   83.73   85.33   85.60   84.53   86.40   85.60   85.60   86.67   86.40   84.53   85.60  100.00   86.40   88.80   85.60   89.60   88.27
   108: Sequence6     76.00   78.93   76.53   77.87   77.60   77.87   75.47   81.07   78.13   79.73   85.07   81.07   85.60   85.33   82.93   85.07   84.53   85.87   84.27   83.73   83.20   83.73   82.93   84.00   83.73   84.27   84.00   84.00   84.00   85.33   82.93   85.33   85.07   82.93   82.40   85.07   84.53   84.80   83.73   84.27   81.07   83.47   82.93   83.20   85.60   86.93   83.47   82.93   80.00   84.53   83.73   85.33   82.67   82.40   82.93   82.40   84.00   84.53   84.27   85.60   85.07   81.87   83.73   83.20   83.20   84.27   84.00   83.73   85.87   85.60   82.40   84.00   80.80   85.87   84.00   84.53   84.53   83.20   85.33   86.13   83.47   82.67   85.33   81.33   84.53   85.60   84.27   84.27   84.53   84.80   83.20   86.13   86.40   84.53   82.40   82.93   84.80   84.00   82.93   83.47   85.87   85.33   86.67   83.47   85.33   85.87   86.40  100.00   85.87   85.60   90.40   87.73
   109: Sequence5     81.60   77.60   76.80   75.73   78.13   78.40   76.53   78.40   77.60   76.27   85.87   82.13   82.67   87.20   82.93   85.60   84.80   81.87   82.40   83.20   85.60   82.93   84.80   85.33   82.13   84.00   84.80   85.87   85.60   85.87   86.13   83.20   82.13   83.47   82.93   83.47   83.47   83.20   83.20   85.07   84.27   81.33   83.47   86.67   83.20   82.67   82.93   85.07   84.53   82.93   85.60   82.93   83.73   84.27   85.07   82.67   84.00   84.27   83.47   84.27   85.07   81.87   82.67   85.33   85.60   84.00   83.73   84.00   83.20   86.13   83.47   81.87   82.40   84.00   84.80   83.47   82.93   84.53   83.47   84.00   83.73   82.93   85.87   85.60   85.33   87.73   85.33   83.20   84.80   84.27   85.07   82.93   86.67   82.93   84.27   83.73   88.00   85.07   85.33   85.60   87.47   85.07   86.93   86.13   84.80   86.93   88.80   85.87  100.00   85.60   90.13   88.27
   110: Sequence13    76.53   76.27   75.73   76.53   78.93   75.47   75.73   79.20   76.53   78.13   84.27   83.47   85.33   82.67   85.60   86.93   86.93   84.53   84.53   85.87   83.73   84.53   83.47   84.80   84.27   84.53   82.40   86.67   83.20   84.53   83.47   85.60   82.13   85.60   84.27   85.60   83.20   83.47   83.73   83.20   85.33   83.47   84.00   82.40   84.27   84.00   84.00   85.07   84.27   84.53   85.87   83.47   84.53   83.47   83.47   82.40   84.53   85.07   83.47   85.33   84.27   84.00   82.13   85.33   85.87   85.87   84.27   82.13   85.33   85.60   84.53   85.87   83.47   82.93   84.53   82.93   83.73   84.27   84.27   84.27   84.27   82.93   85.33   83.20   87.73   85.07   84.80   82.93   86.13   84.27   85.07   85.60   83.73   84.80   85.60   86.67   85.07   85.33   85.33   84.00   85.33   85.60   86.40   85.87   84.53   86.93   85.60   85.60   85.60  100.00   89.87   90.67
   111: Sequence111   78.67   78.40   76.27   77.33   78.67   78.40   77.07   78.67   78.67   78.13   88.00   86.40   88.53   88.00   87.73   88.27   87.73   86.67   87.47   87.47   86.67   87.20   87.47   87.20   86.40   87.47   86.67   88.27   88.53   88.00   87.47   87.47   86.67   86.67   86.93   87.47   87.73   87.73   86.13   88.00   86.93   86.40   87.20   88.00   86.93   88.27   86.67   87.47   88.27   88.53   88.00   87.73   88.00   88.27   88.53   86.40   88.53   89.07   88.53   85.87   88.27   86.40   86.40   88.53   87.73   86.67   86.40   86.40   88.53   88.00   87.20   86.13   86.40   86.67   88.00   88.00   88.27   87.20   88.27   87.47   88.53   87.20   87.20   87.47   87.73   87.73   88.00   88.00   88.27   88.53   88.27   88.53   88.80   88.80   88.27   88.27   89.07   88.80   88.80   89.07   89.07   89.07   89.33   89.60   89.33   88.80   89.60   90.40   90.13   89.87  100.00   94.67
   112: Sequence112   78.40   77.07   75.47   76.80   79.20   78.67   76.80   78.67   78.40   76.00   87.47   86.40   88.00   86.93   86.93   88.00   87.47   85.87   86.93   89.60   86.67   87.73   88.00   86.40   86.40   87.20   85.87   89.33   88.00   87.73   86.67   88.53   85.33   86.93   86.67   87.47   86.13   86.13   85.60   87.20   87.73   86.67   85.87   86.67   85.33   87.47   86.67   86.93   87.47   88.80   86.93   88.00   87.47   88.53   87.47   85.60   87.73   87.73   87.73   85.60   85.33   85.33   85.07   88.27   88.00   88.00   86.13   84.00   87.47   88.00   86.13   85.07   87.47   86.67   87.73   86.13   86.13   86.67   87.73   86.13   87.20   87.73   86.67   86.40   87.73   87.73   85.87   87.20   88.00   88.27   88.53   88.80   86.93   87.73   88.80   88.80   89.07   86.13   88.00   88.00   87.73   88.80   87.20   88.53   88.53   89.60   88.27   87.73   88.27   90.67   94.67  100.00
```
