## Supplemental Figure 3 for "A selectable system to evaluate synthetic gene optimization features"

```
Supplemental Figure 3. Codon variants of sh ble evaluated in E.coli. (A) nucleotide sequences, (B) multiple sequence alignment using clustal Omega, and (C) percent identity matrix of the sequences, (D) Pairwise alignment of Seq7 and Seq109.
A.

>sh ble wt
ATGGCCAAGTTGACCAGTGCCGTTCCGGTGCTCACCGCGCGCGACGTCGCCGGAGCGGTCGAGTTCTGGACCGACCGGCTCGGGTTCTCCCGGGACTTCGTGGAGGACGACTTCGCCGGTGTGGTCCGGGACGACGTGACCCTGTTCATCAGCGCGGTCCAGGACCAGGTGGTGCCGGACAACACCCTGGCCTGGGTGTGGGTGCGCGGCCTGGACGAGCTGTACGCCGAGTGGTCGGAGGTCGTGTCCACGAACTTCCGGGACGCCTCCGGGCCGGCCATGACCGAGATCGGCGAGCAGCCGTGGGGGCGGGAGTTCGCCCTGCGCGACCCGGCCGGCAACTGCGTGCACTTCGTGGCCGAGGAGCAGGACTGA

>Seq111
ATGGCGAAACTGACCAGCGCGGTGCCGGTGCTGACCGCGCGCGATGTGGCGGGCGCGGTGGAATTTTGGACCGATCGCCTGGGCTTTAGCCGCGATTTTGTGGAAGATGATTTTGCGGGCGTGGTGCGCGATGATGTGACCCTGTTTATTAGCGCGGTGCAGGATCAGGTGGTGCCGGATAACACCCTGGCGTGGGTGTGGGTGCGCGGCCTGGATGAACTGTATGCGGAATGGAGCGAAGTGGTGAGCACCAACTTTCGCGATGCGAGCGGCCCGGCGATGACCGAAATTGGCGAACAGCCGTGGGGCCGCGAATTTGCGCTGCGCGATCCGGCGGGCAACTGCGTGCATTTTGTGGCGGAAGAACAGGATTAA

>Seq112
ATGGCCAAACTGACCAGCGCCGTTCCGGTGCTGACCGCGCGCGATGTGGCCGGCGCGGTTGAATTTTGGACCGATCGCCTGGGCTTTAGCCGCGATTTTGTGGAAGATGATTTTGCCGGTGTGGTTCGTGATGATGTGACCCTGTTTATCAGCGCGGTGCAGGATCAGGTGGTGCCGGATAACACCCTGGCCTGGGTGTGGGTGCGCGGCCTGGATGAACTGTATGCCGAATGGAGCGAAGTGGTGAGCACCAACTTTCGTGATGCCAGCGGCCCGGCCATGACCGAAATCGGCGAACAGCCGTGGGGCCGCGAATTTGCCCTGCGCGATCCGGCCGGCAACTGCGTGCATTTTGTGGCCGAAGAACAGGATTGA

>Seq103
ATGGCGAAACTTACTTCCGCGGTTCCGGTCCTAACCGCTCGCGATGTCGCGGGTGCTGTGGAGTTCTGGACCGACCGTCTCGGCTTCTCGCGTGATTTCGTGGAGGATGACTTCGCCGGAGTGGTCCGTGATGACGTAACGCTGTTCATTTCTGCAGTACAAGACCAGGTCGTGCCCGACAACACATTGGCCTGGGTTTGGGTACGGGGCCTCGATGAGTTATATGCCGAATGGAGTGAAGTCGTAAGCACCAATTTTCGCGATGCCTCAGGACCAGCAATGACGGAAATTGGCGAGCAACCGTGGGGGCGAGAATTCGCTCTGCGCGACCCTGCAGGCAATTGCGTACATTTTGTCGCCGAAGAGCAGGATTAA

>Seq109
ATGGCTAAATTGACGAGTGCGGTACCAGTTTTAACGGCACGTGACGTTGCAGGGGCTGTAGAATTCTGGACCGATCGGCTCGGCTTTTCACGGGATTTTGTAGAGGACGATTTTGCAGGGGTGGTCAGGGATGACGTTACCCTTTTCATCAGCGCGGTACAAGATCAAGTCGTGCCGGACAATACGTTAGCCTGGGTGTGGGTTCGCGGTCTAGATGAGCTGTATGCAGAATGGTCCGAGGTCGTATCTACTAACTTTCGCGACGCTTCGGGTCCCGCGATGACAGAAATCGGCGAACAACCGTGGGGCCGAGAATTCGCCTTACGTGATCCTGCGGGAAATTGCGTTCATTTCGTAGCGGAAGAACAGGACTAA

>Seq20
ATGGCCAAACTGACCTCAGCCGTGCCGGTGCTCACCGCCCGTGATGTGGCGGGCGCGGTTGAGTTTTGGACTGACCGTCTGGGCTTTAGCCGCGACTTTGTAGAAGACGATTTCGCGGGTGTTGTTCGTGATGATGTGACATTATTTATCAGCGCGGTCCAGGACCAGGTGGTGCCGGATAACACCCTGGCGTGGGTGTGGGTGCGCGGCCTGGATGAACTGTATGCGGAATGGAGCGAAGTTGTGAGCACCAACTTCCGTGATGCGAGCGGCCCGGCGATGACCGAAATTGGTGAGCAACCGTGGGGTCGCGAATTCGCCCTTCGCGATCCGGCAGGGAACTGCGTTCATTTTGTGGCAGAAGAACAGGATTAA

>Seq51
ATGGCCAAACTGACCTCAGCGGTTCCGGTGCTGACCGCCCGCGACGTGGCGGGAGCGGTTGAATTTTGGACCGACCGCCTGGGTTTCAGCCGCGACTTTGTGGAAGATGACTTTGCGGGCGTGGTTCGTGATGATGTGACACTGTTTATTAGTGCTGTTCAAGATCAGGTCGTACCGGATAACACCCTGGCGTGGGTCTGGGTACGTGGCCTGGATGAACTGTATGCCGAGTGGAGCGAGGTGGTGAGCACCAATTTTCGTGATGCAAGCGGCCCGGCTATGACCGAAATTGGCGAACAGCCGTGGGGTCGCGAATTCGCCCTCCGTGATCCGGCGGGGAACTGCGTACATTTCGTTGCAGAAGAACAGGATTGA

>Seq95
ATGGCCAAACTTACGTCGGCGGTGCCGGTCCTGACTGCCCGCGACGTGGCGGGCGCAGTGGAGTTTTGGACGGATCGCTTAGGTTTCAGCCGTGACTTTGTGGAGGATGATTTTGCGGGCGTTGTGCGCGACGATGTAACCCTGTTTATCTCAGCCGTTCAAGATCAGGTTGTTCCGGATAACACCCTGGCGTGGGTCTGGGTCCGCGGTCTCGATGAATTGTATGCCGAATGGAGCGAAGTTGTTAGCACCAATTTCCGCGACGCTAGCGGCCCGGCGATGACCGAAATCGGCGAACAGCCGTGGGGTCGTGAATTTGCGCTGCGTGATCCTGCGGGCAACTGCGTGCATTTCGTGGCGGAAGAACAGGATTGA

>Seq7
ATGGCCAAACTGACGAGCGCGGTTCCGGTCCTGACCGCGCGTGATGTCGCAGGCGCGGTGGAATTTTGGACCGATCGCCTGGGCTTTAGTCGCGACTTTGTTGAGGACGATTTTGCCGGCGTGGTGCGCGATGATGTAACCTTATTTATTTCAGCGGTGCAGGATCAGGTGGTTCCGGATAACACCCTGGCCTGGGTTTGGGTTCGCGGCCTCGATGAACTGTATGCGGAATGGAGCGAAGTGGTATCGACCAATTTTCGTGATGCTAGCGGTCCGGCGATGACGGAAATCGGTGAACAGCCGTGGGGGCGCGAATTTGCCCTGCGTGATCCGGCAGGTAACTGCGTTCATTTTGTGGCCGAAGAACAAGATTAA


B.
CLUSTAL O(1.2.4) multiple sequence alignment


Seq109      ATGGCTAAATTGACGAGTGCGGTACCAGTTTTAACGGCACGTGACGTTGCAGGGGCTGTA	60
Seq103      ATGGCGAAACTTACTTCCGCGGTTCCGGTCCTAACCGCTCGCGATGTCGCGGGTGCTGTG	60
wt          ATGGCCAAGTTGACCAGTGCCGTTCCGGTGCTCACCGCGCGCGACGTCGCCGGAGCGGTC	60
Seq95       ATGGCCAAACTTACGTCGGCGGTGCCGGTCCTGACTGCCCGCGACGTGGCGGGCGCAGTG	60
Seq7        ATGGCCAAACTGACGAGCGCGGTTCCGGTCCTGACCGCGCGTGATGTCGCAGGCGCGGTG	60
Seq51       ATGGCCAAACTGACCTCAGCGGTTCCGGTGCTGACCGCCCGCGACGTGGCGGGAGCGGTT	60
Seq20       ATGGCCAAACTGACCTCAGCCGTGCCGGTGCTCACCGCCCGTGATGTGGCGGGCGCGGTT	60
Seq111      ATGGCGAAACTGACCAGCGCGGTGCCGGTGCTGACCGCGCGCGATGTGGCGGGCGCGGTG	60
Seq112      ATGGCCAAACTGACCAGCGCCGTTCCGGTGCTGACCGCGCGCGATGTGGCCGGCGCGGTT	60
            ***** **  * **    ** ** ** **  * ** ** ** ** ** ** ** ** **

Seq109      GAATTCTGGACCGATCGGCTCGGCTTTTCACGGGATTTTGTAGAGGACGATTTTGCAGGG	120
Seq103      GAGTTCTGGACCGACCGTCTCGGCTTCTCGCGTGATTTCGTGGAGGATGACTTCGCCGGA	120
wt          GAGTTCTGGACCGACCGGCTCGGGTTCTCCCGGGACTTCGTGGAGGACGACTTCGCCGGT	120
Seq95       GAGTTTTGGACGGATCGCTTAGGTTTCAGCCGTGACTTTGTGGAGGATGATTTTGCGGGC	120
Seq7        GAATTTTGGACCGATCGCCTGGGCTTTAGTCGCGACTTTGTTGAGGACGATTTTGCCGGC	120
Seq51       GAATTTTGGACCGACCGCCTGGGTTTCAGCCGCGACTTTGTGGAAGATGACTTTGCGGGC	120
Seq20       GAGTTTTGGACTGACCGTCTGGGCTTTAGCCGCGACTTTGTAGAAGACGATTTCGCGGGT	120
Seq111      GAATTTTGGACCGATCGCCTGGGCTTTAGCCGCGATTTTGTGGAAGATGATTTTGCGGGC	120
Seq112      GAATTTTGGACCGATCGCCTGGGCTTTAGCCGCGATTTTGTGGAAGATGATTTTGCCGGT	120
            ** ** ***** ** **  * ** **    ** ** ** ** ** ** ** ** ** **

Seq109      GTGGTCAGGGATGACGTTACCCTTTTCATCAGCGCGGTACAAGATCAAGTCGTGCCGGAC	180
Seq103      GTGGTCCGTGATGACGTAACGCTGTTCATTTCTGCAGTACAAGACCAGGTCGTGCCCGAC	180
wt          GTGGTCCGGGACGACGTGACCCTGTTCATCAGCGCGGTCCAGGACCAGGTGGTGCCGGAC	180
Seq95       GTTGTGCGCGACGATGTAACCCTGTTTATCTCAGCCGTTCAAGATCAGGTTGTTCCGGAT	180
Seq7        GTGGTGCGCGATGATGTAACCTTATTTATTTCAGCGGTGCAGGATCAGGTGGTTCCGGAT	180
Seq51       GTGGTTCGTGATGATGTGACACTGTTTATTAGTGCTGTTCAAGATCAGGTCGTACCGGAT	180
Seq20       GTTGTTCGTGATGATGTGACATTATTTATCAGCGCGGTCCAGGACCAGGTGGTGCCGGAT	180
Seq111      GTGGTGCGCGATGATGTGACCCTGTTTATTAGCGCGGTGCAGGATCAGGTGGTGCCGGAT	180
Seq112      GTGGTTCGTGATGATGTGACCCTGTTTATCAGCGCGGTGCAGGATCAGGTGGTGCCGGAT	180
            ** **  * ** ** ** **  * ** **    ** ** ** ** ** ** ** ** **

Seq109      AATACGTTAGCCTGGGTGTGGGTTCGCGGTCTAGATGAGCTGTATGCAGAATGGTCCGAG	240
Seq103      AACACATTGGCCTGGGTTTGGGTACGGGGCCTCGATGAGTTATATGCCGAATGGAGTGAA	240
wt          AACACCCTGGCCTGGGTGTGGGTGCGCGGCCTGGACGAGCTGTACGCCGAGTGGTCGGAG	240
Seq95       AACACCCTGGCGTGGGTCTGGGTCCGCGGTCTCGATGAATTGTATGCCGAATGGAGCGAA	240
Seq7        AACACCCTGGCCTGGGTTTGGGTTCGCGGCCTCGATGAACTGTATGCGGAATGGAGCGAA	240
Seq51       AACACCCTGGCGTGGGTCTGGGTACGTGGCCTGGATGAACTGTATGCCGAGTGGAGCGAG	240
Seq20       AACACCCTGGCGTGGGTGTGGGTGCGCGGCCTGGATGAACTGTATGCGGAATGGAGCGAA	240
Seq111      AACACCCTGGCGTGGGTGTGGGTGCGCGGCCTGGATGAACTGTATGCGGAATGGAGCGAA	240
Seq112      AACACCCTGGCCTGGGTGTGGGTGCGCGGCCTGGATGAACTGTATGCCGAATGGAGCGAA	240
            ** **  * ** ***** ***** ** ** ** ** **  * ** ** ** ***   **

Seq109      GTCGTATCTACTAACTTTCGCGACGCTTCGGGTCCCGCGATGACAGAAATCGGCGAACAA	300
Seq103      GTCGTAAGCACCAATTTTCGCGATGCCTCAGGACCAGCAATGACGGAAATTGGCGAGCAA	300
wt          GTCGTGTCCACGAACTTCCGGGACGCCTCCGGGCCGGCCATGACCGAGATCGGCGAGCAG	300
Seq95       GTTGTTAGCACCAATTTCCGCGACGCTAGCGGCCCGGCGATGACCGAAATCGGCGAACAG	300
Seq7        GTGGTATCGACCAATTTTCGTGATGCTAGCGGTCCGGCGATGACGGAAATCGGTGAACAG	300
Seq51       GTGGTGAGCACCAATTTTCGTGATGCAAGCGGCCCGGCTATGACCGAAATTGGCGAACAG	300
Seq20       GTTGTGAGCACCAACTTCCGTGATGCGAGCGGCCCGGCGATGACCGAAATTGGTGAGCAA	300
Seq111      GTGGTGAGCACCAACTTTCGCGATGCGAGCGGCCCGGCGATGACCGAAATTGGCGAACAG	300
Seq112      GTGGTGAGCACCAACTTTCGTGATGCCAGCGGCCCGGCCATGACCGAAATCGGCGAACAG	300
            ** **    ** ** ** ** ** **    ** ** ** ***** ** ** ** ** **

Seq109      CCGTGGGGCCGAGAATTCGCCTTACGTGATCCTGCGGGAAATTGCGTTCATTTCGTAGCG	360
Seq103      CCGTGGGGGCGAGAATTCGCTCTGCGCGACCCTGCAGGCAATTGCGTACATTTTGTCGCC	360
wt          CCGTGGGGGCGGGAGTTCGCCCTGCGCGACCCGGCCGGCAACTGCGTGCACTTCGTGGCC	360
Seq95       CCGTGGGGTCGTGAATTTGCGCTGCGTGATCCTGCGGGCAACTGCGTGCATTTCGTGGCG	360
Seq7        CCGTGGGGGCGCGAATTTGCCCTGCGTGATCCGGCAGGTAACTGCGTTCATTTTGTGGCC	360
Seq51       CCGTGGGGTCGCGAATTCGCCCTCCGTGATCCGGCGGGGAACTGCGTACATTTCGTTGCA	360
Seq20       CCGTGGGGTCGCGAATTCGCCCTTCGCGATCCGGCAGGGAACTGCGTTCATTTTGTGGCA	360
Seq111      CCGTGGGGCCGCGAATTTGCGCTGCGCGATCCGGCGGGCAACTGCGTGCATTTTGTGGCG	360
Seq112      CCGTGGGGCCGCGAATTTGCCCTGCGCGATCCGGCCGGCAACTGCGTGCATTTTGTGGCC	360
            ******** ** ** ** **  * ** ** ** ** ** ** ***** ** ** ** **

Seq109      GAAGAACAGGACTAA	375
Seq103      GAAGAGCAGGATTAA	375
sh          GAGGAGCAGGACTGA	375
Seq95       GAAGAACAGGATTGA	375
Seq7        GAAGAACAAGATTAA	375
Seq51       GAAGAACAGGATTGA	375
Seq20       GAAGAACAGGATTAA	375
Seq111      GAAGAACAGGATTAA	375
Seq112      GAAGAACAGGATTGA	375
            ** ** ** ** * *


C.

#
#
#  Percent Identity  Matrix - created by Clustal2.1
#
#
		 Seq109	Seq103	wt     Seq95   Seq7    Seq51   Seq20   Seq111  Seq112
     1: Seq109      100.00   76.53   78.13   74.67   77.87   74.13   74.40   77.33   76.80
     2: Seq103       76.53  100.00   78.40   78.13   78.40   78.67   77.07   78.67   78.40
     3: wt           78.13   78.40  100.00   75.47   76.53   78.40   79.47   79.20   83.47
     4: Seq95        74.67   78.13   75.47  100.00   84.00   85.60   82.13   86.40   84.00
     5: Seq7         77.87   78.40   76.53   84.00  100.00   82.93   84.53   88.53   88.27
     6: Seq51        74.13   78.67   78.40   85.60   82.93  100.00   87.20   88.27   88.53
     7: Seq20        74.40   77.07   79.47   82.13   84.53   87.20  100.00   89.07   89.07
     8: Seq111       77.33   78.67   79.20   86.40   88.53   88.27   89.07  100.00   94.67
     9: Seq112       76.80   78.40   83.47   84.00   88.27   88.53   89.07   94.67  100.00


D.


########################################
# Program: needle
# Rundate: Wed 30 Oct 2019 23:27:24
# Commandline: needle
#    -auto
#    -stdout
#    -asequence emboss_needle-I20191030-232721-0329-31199833-p2m.asequence
#    -bsequence emboss_needle-I20191030-232721-0329-31199833-p2m.bsequence
#    -datafile EDNAFULL
#    -gapopen 10.0
#    -gapextend 0.5
#    -endopen 10.0
#    -endextend 0.5
#    -aformat3 pair
#    -snucleotide1
#    -snucleotide2
# Align_format: pair
# Report_file: stdout
########################################

#=======================================
#
# Aligned_sequences: 2
# 1: Seq7
# 2: Seq109
# Matrix: EDNAFULL
# Gap_penalty: 10.0
# Extend_penalty: 0.5
#
# Length: 380
# Identity:     293/380 (77.1%)
# Similarity:   293/380 (77.1%)
# Gaps:          10/380 ( 2.6%)
# Score: 1133.0
#
#
#=======================================

Seq7               1 ATGGCCAAACTGACGAGCGCGGTTCCGGTCCTGACCGCGCGTGATGTCGC     50
                     |||||.|||.|||||||.|||||.||.||..|.||.||.|||||.||.||
Seq109             1 ATGGCTAAATTGACGAGTGCGGTACCAGTTTTAACGGCACGTGACGTTGC     50

Seq7              51 AGGCGCGGTGGAATTTTGGACCGATCGCCTGGGCTTTAGTCGCGACTTTG    100
                     |||.||.||.|||||.|||||||||||.||.||||||...||.||.||||
Seq109            51 AGGGGCTGTAGAATTCTGGACCGATCGGCTCGGCTTTTCACGGGATTTTG    100

Seq7             101 TTGAGGACGATTTTGCCGGCGTGGTGCGCGATGATGTAACCTTATTTATT    150
                     |.||||||||||||||.||.|||||..|.|||||.||.|||.|     ||
Seq109           101 TAGAGGACGATTTTGCAGGGGTGGTCAGGGATGACGTTACCCT-----TT    145

Seq7             151 TCA-----GCGGTGCAGGATCAGGTGGTTCCGGATAACACCCTGGCCTGG    195
                     |||     |||||.||.|||||.||.||.|||||.||.||..|.||||||
Seq109           146 TCATCAGCGCGGTACAAGATCAAGTCGTGCCGGACAATACGTTAGCCTGG    195

Seq7             196 GTTTGGGTTCGCGGCCTCGATGAACTGTATGCGGAATGGAGCGAAGTGGT    245
                     ||.|||||||||||.||.|||||.||||||||.||||||..|||.||.||
Seq109           196 GTGTGGGTTCGCGGTCTAGATGAGCTGTATGCAGAATGGTCCGAGGTCGT    245

Seq7             246 ATCGACCAATTTTCGTGATGCTAGCGGTCCGGCGATGACGGAAATCGGTG    295
                     |||.||.||.|||||.||.|||...|||||.||||||||.||||||||.|
Seq109           246 ATCTACTAACTTTCGCGACGCTTCGGGTCCCGCGATGACAGAAATCGGCG    295

Seq7             296 AACAGCCGTGGGGGCGCGAATTTGCCCTGCGTGATCCGGCAGGTAACTGC    345
                     ||||.||||||||.||.|||||.|||.|.||||||||.||.||.||.|||
Seq109           296 AACAACCGTGGGGCCGAGAATTCGCCTTACGTGATCCTGCGGGAAATTGC    345

Seq7             346 GTTCATTTTGTGGCCGAAGAACAAGATTAA    375
                     ||||||||.||.||.||||||||.||.|||
Seq109           346 GTTCATTTCGTAGCGGAAGAACAGGACTAA    375


#---------------------------------------
#---------------------------------------
```
