## Supplementary figures and images for "A selectable system to evaluate synthetic gene optimization features"

### Supplemental Figure 4

**Supplemental Figure XX.** Map of pTWIST-CMV-sh ble.

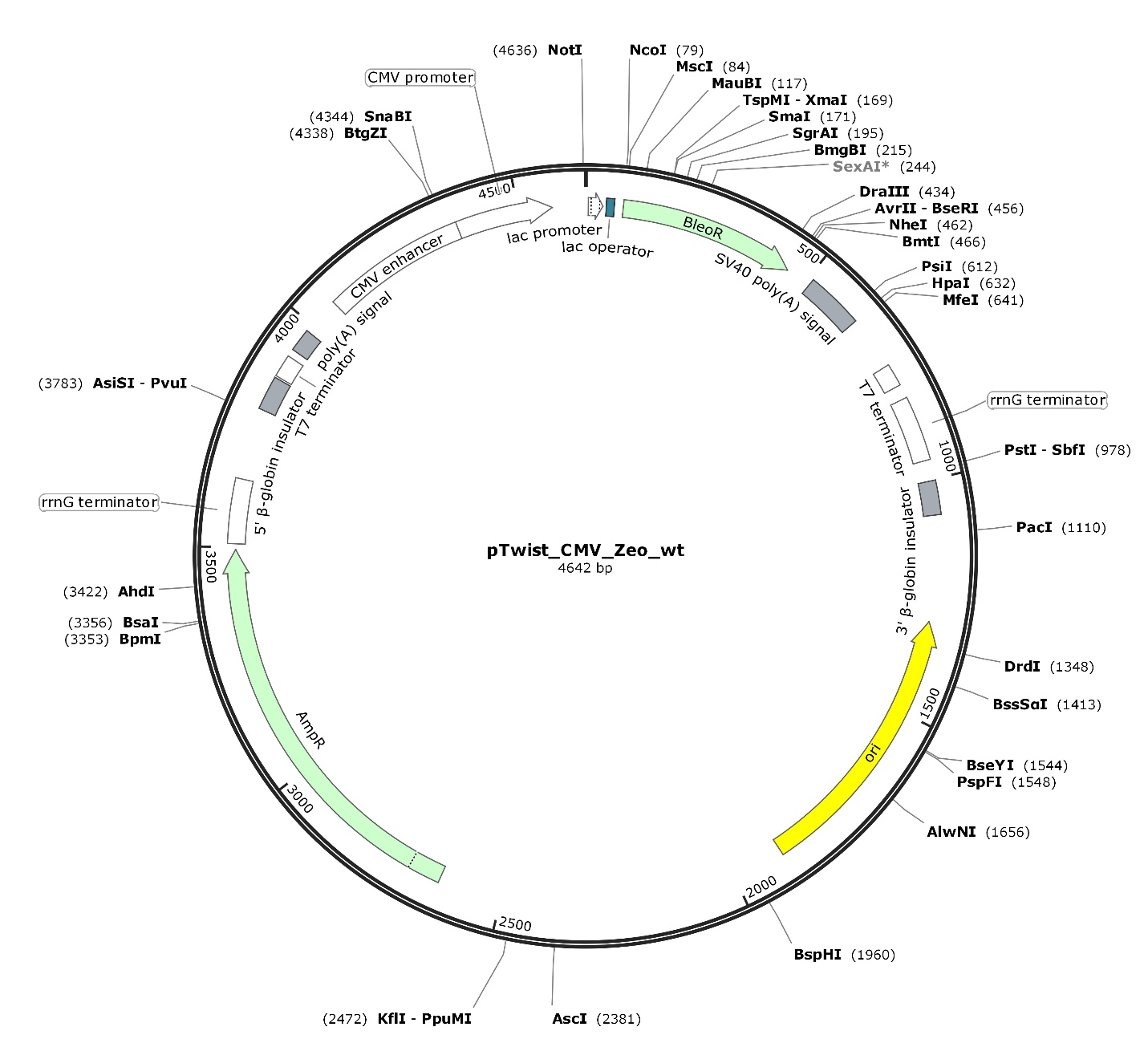

### Supplemental Figure 5

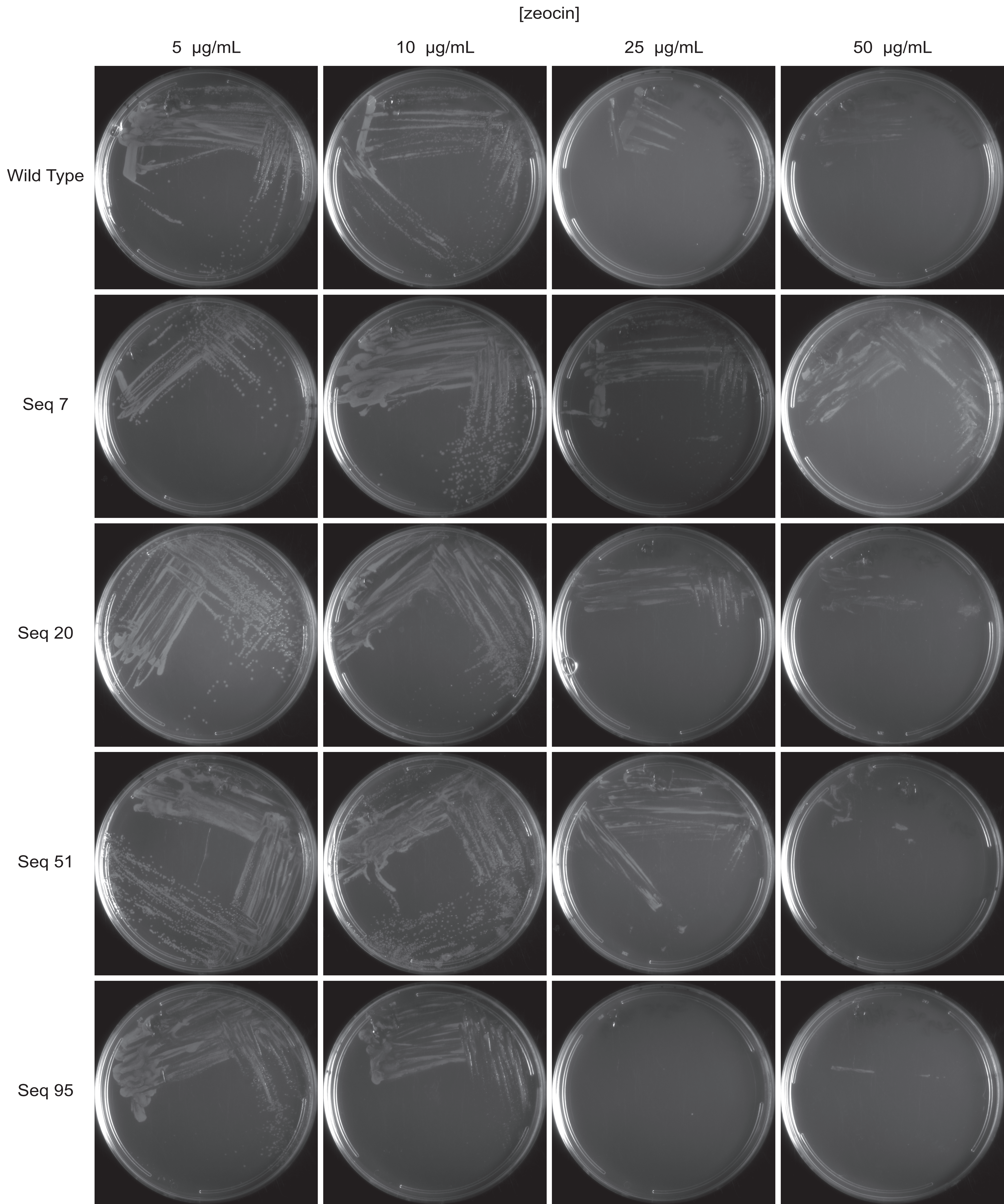

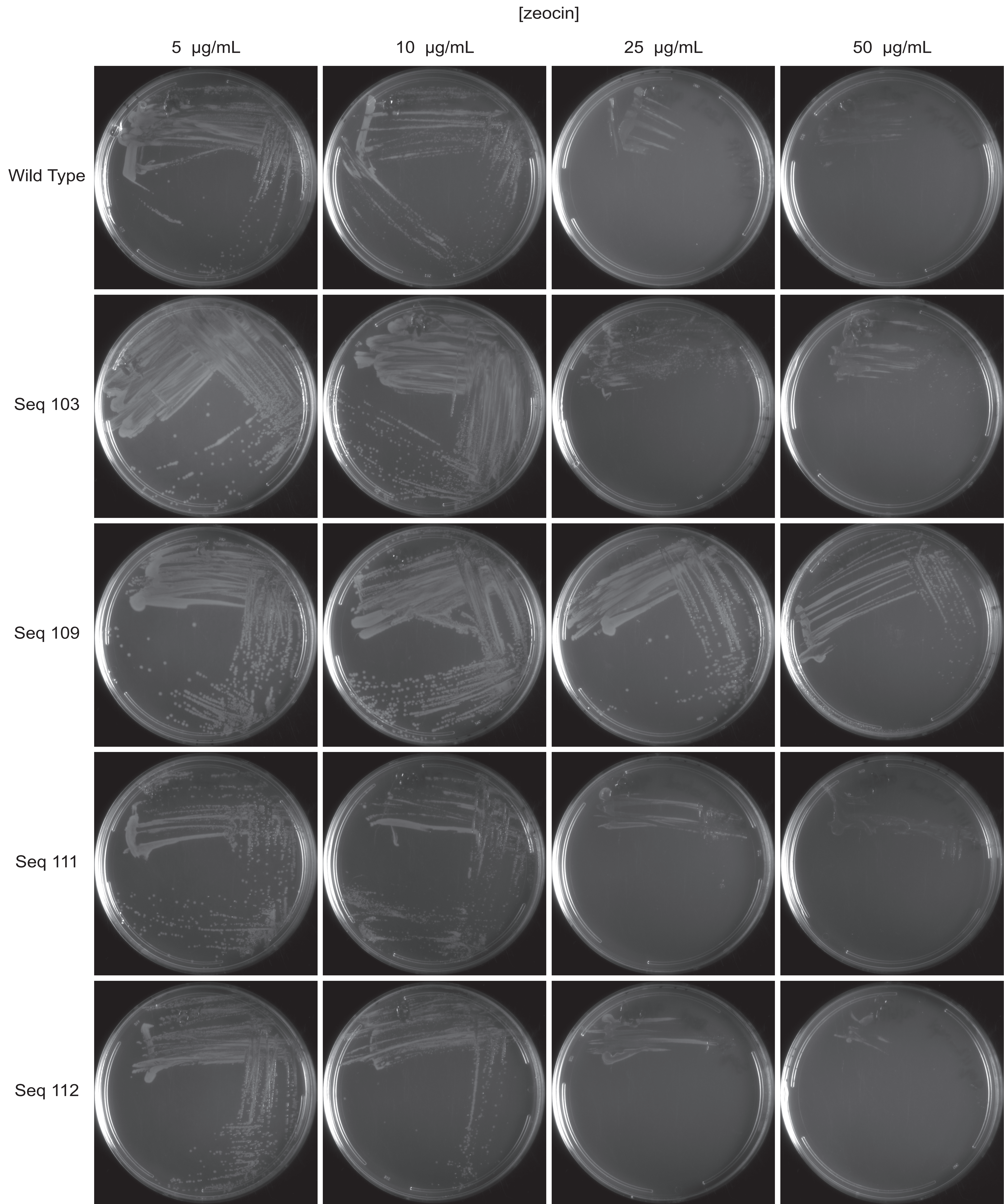
