## Supplemental Methods for "A selectable system to evaluate synthetic gene optimization features"

*Gene sequence genesis*

This sequence genesis program was written in JAVA and contains three algorithms which were designed to generate DNA sequences in order to select genes with enhanced expression. The three different optimization algorithms are random generation, high frequency generation, and codon replacement. Each algorithm has different inputs that must be met in order to execute the program. All three algorithms require a codon frequency chart for the species the gene will be expressed, which can be found at <https://www.kazusa.or.jp/codon/>. Under the format section, one must select “standard” and “a style like CodonFrequency output in GCG [Wisconsin Package^TM^](http://www.gcg.com/products/software.html)”. In addition to the codon frequency chart, the random and high frequency generation algorithms also require a protein sequence which will be used for the optimization. The random generation algorithm will also require one to enter the amount of sequences that should be generated. Lastly, the codon replacement algorithm requires a DNA sequence instead of a protein sequence to execute properly. Once the necessary information is filled in, the code will be able to execute without error as long as the input formatting was correct.

The first task the code accomplishes when it is run is the collection of data. The following information is gathered during the data collection: codon, amino acid, and sequence. First, the codon frequency chart is used to collect the codon data which consists of the codon sequence and the frequency of the codon out of 1000. Second, a codon table is used to gather the amino acid data which consists of the amino acid name, abbreviation, and codon data. Third, the protein sequence is gathered and stored. Lastly, the amount of each amino acid is gathered and individually stored.

After the necessary data is collected, codon evaluation takes place which calculates a new frequency and determines whether a codon is favored or unfavored. The new codon frequency is the percentage of how often the codon appears for an amino acid out of 100. When all of the codon percentages, or new frequencies, for a specific amino acid are added up the sum will equal 1.0 (or 100%). A codon’s old frequency is the frequency gathered from the codon frequency chart when the data is initially collected. In order to determine the codon percentage, all of the old frequencies of the codons of an amino acid are added up. Each codon is then assigned a new frequency which can be determined by dividing the codon’s old frequency by the sum of codon old frequencies (New Frequency = Old Frequency / Sum Old Frequencies). After the new frequencies are calculated, codons are classified as favored or unfavored. The favoring system is essential to the random generation algorithm and is utilized in the majority of the codon generation process. The first step to classify the codons as favored or unfavored is to find the average codon frequency for the amino acid. This can be found by summing the new frequencies and dividing by the amount of codons the amino acid has (Sum New Frequencies / Codon Amount). Each individual codon is then classified as favored or unfavored based on the following conditions: a codon is favored if the new frequency is greater than or equal to the average frequency and a codon is unfavored if the new frequency is less than the average frequency.

The most complex codon optimization algorithm is the random generation. In short, this algorithm generates codons for an amino acid before cycling on to the next one. The amino acids are cycled alphabetically, so the first codons that are generated belong to Alanine. Once an amino acid has been selected for generation, the program cycles through the protein sequence and if there is the same amino acid in the sequence as the one selected, a codon is generated. Codon generation begins by generating a random decimal between zero and one. This number can then be used to randomly select a codon because the sum of new frequencies of codons for an amino acid add up to one. The codon is then selected based on which frequency range the randomly generated number is in. Consider the following scenario; there are four codons (with new frequencies of 0.10/0.20/0.30/0.40) and the randomly generated number was 0.55. In order to determine which codon is generated, the program must find which range the randomly generated lies under and what codon the range belongs to. Finding the range requires addition and starts at 0. The ranges for the scenario described above would be:

0.0 ≤ Codon 1 < 0.10, 0.10 ≤ Codon 2 < 0.30, 0.30 ≤ Codon 3 < 0.60, 0.60 ≤ Codon 4 < 1.0. The range can be found by adding the codon’s new frequency to the sum of the new frequencies of the codons before it. For example, the range for Codon 3 is 0.30 ≤ Codon 3 < 0.60. The minimum value of the range, 0.30, can be found by summing the new frequencies of the codons before it. In this case, the new frequencies of Codons 1 and 2, which add up to 0.30 (0.10 + 0.20 = 0.30). The maximum value of the range, 0.60, can be found by adding that codon’s new frequency to the sum of the new frequencies of the codons before it. In this case, adding the frequency of Codon 3, 0.30, to the sum of the new frequencies of the codons before it, 0.30. This results in the maximum range of 0.60 (0.30 + 0.30 = 0.60). Now the codon that should be generated can be identified based on the ranges of the codons and the randomly generated number. Since 0.55 lies in the range of Codon 3, 0.30 ≤ Codon 3 < 0.60, the codon that is generated is Codon 3. This codon is not directly added to the sequence, it is instead stored for later and will be implemented into the sequence after all the codon generation has completed. This process then repeats for every amino acid located in the sequence that is the same as the selected amino acid. However, a different generation process is used if there are more codons for the selected amino acid than the amount of the same amino acid in the sequence. For example, if an amino acid has four codons, but that amino acid only occurs three times in the protein sequence. This new generation process involves only generating favored codons, however, it still uses the range method that was mentioned in the previous process. The only difference is that only favored codons are involved in the range method. For example, if there were four codons and codons 1 and 3 are favored, then the range process would only involve the new frequencies of those codons. Additionally, the randomly generated number is between 0 and the sum of the new frequencies of the favored codons.
